# Plasticity-induced actin polymerization in the dendritic shaft regulates intracellular AMPA receptor trafficking

**DOI:** 10.1101/2022.05.29.493906

**Authors:** V. C. Wong, P.R. Houlihan, H. Liu, D. Walpita, M.C. DeSantis, Z. Liu, E. K. O’Shea

**Affiliations:** Janelia Research Campus, Howard Hughes Medical Institute, Ashburn, VA, 20147

## Abstract

AMPA-type receptors (AMPARs) are rapidly inserted into synapses undergoing long-term potentiation (LTP) to increase synaptic transmission, but how AMPAR-containing vesicles are selectively trafficked to these synapses during LTP is not known. Here we developed a strategy to label AMPAR GluA1 subunits expressed from the endogenous loci of rat hippocampal neurons such that the motion of GluA1-containing vesicles in time-lapse sequences can be characterized using single-particle tracking and mathematical modeling. We find that GluA1- containing vesicles are confined and concentrated near sites of stimulation-induced plasticity. We show that confinement is mediated by actin polymerization, which hinders the active transport of GluA1-containing vesicles along the length of the dendritic shaft by modulating the rheological properties of the cytoplasm. Actin polymerization also facilitates myosin-mediated transport of GluA1-containing vesicles to exocytic sites. We conclude that neurons utilize F-actin to increase vesicular GluA1 reservoirs and promote exocytosis proximal to the sites of neuronal activity.

## Introduction

Synaptic plasticity – the modulation of synaptic connections in response to changes in neuronal activity – is regarded as the cellular basis for learning and memory (Abbot and Nelson, 2000; Citri and Malenka, 2008; Magee and Greinberger, 2020). Long-term potentiation (LTP) is a lasting form of synaptic plasticity in which a synapse is strengthened through persistent activity (Citri and Malenka, 2008). Canonical LTP – the most well-characterized form of LTP – is triggered by activation of the N-methyl-D-aspartate receptors (NMDARs), which facilitate the influx of calcium ions from the synaptic cleft into the postsynaptic compartment (Bliss, 2011).

Calcium binds to calcium/calmodulin-dependent protein kinase II (CaMKII), triggering signaling pathways that lead to LTP (Lisman et al., 2012). Although the molecular mechanism mediating LTP is still under debate (Bliss and Collingridge, 2013), substantial evidence suggests that α- amino-3-hydroxy-5-methyl-4-isoxazolepropionic acid receptors (AMPARs) play a critical role (Park, 2018). AMPARs are a class of ionotropic glutamate receptors that permit the influx of sodium, potassium, and in certain cases calcium ions (Chater and Goda, 2014). The activation of CaMKII signaling results in a rapid increase in the abundance and conductance of AMPARs at the synapse, increasing the inward flow of ions in response to stimulation (Lisman et al., 2012). This increase in AMPAR-mediated current is thought to underlie LTP (Herring and Nicoll, 2016).

Given that the number of AMPARs at synapses is tightly regulated during LTP and other forms of plasticity, AMPAR trafficking has been studied extensively (Groc and Choquet, 2020). The majority of AMPARs are synthesized in the cell body and then enter the secretory pathway, where they can be trafficked in vesicles and ultimately inserted into the neuronal membrane via exocytosis (Shepard and Huganir, 2007). AMPARs on the membrane can diffuse into synapses and can also be endocytosed and delivered to lysosomes for degradation or recycled through the endocytic pathway (Choquet, 2022; Goo et al., 2017). An important feature of LTP is input specificity – LTP stimulated at one synapse does not spread to inactive synapses (Kandel et al., 2013) – suggesting that AMPAR trafficking is regulated such that AMPARs are inserted selectively in synapses that are undergoing LTP. The mechanisms that regulate how AMPARs are delivered to synapses undergoing LTP are not fully understood, but evidence points to two models: (1) AMPARs in vesicles may be trafficked and exocytosed directly into synapses undergoing LTP; and (2) AMPARs diffusing in the neuronal membrane may be selectively trapped at synapses undergoing LTP.

AMPARs in vesicles are trafficked through dendrites by both microtubule- and actin-based active transport (Setou et al., 2002; Hoogenraad et al., 2005; Correia et al., 2008; Wang et al., 2008; Hoerndli et al., 2013; Wagner et al., 2019). Importantly, disrupting either microtubule- or actin-based transport inhibits AMPAR-mediated currents and also interferes with LTP (Correia et al., 2008; Wang et al., 2008; Hoerndli et al., 2013), demonstrating that the active transport of AMPARs plays a role in plasticity. Nevertheless, whether AMPAR-containing vesicles (hereafter referred to as AMPAR vesicles) are delivered to loci undergoing neuronal activity is unclear, due in large part to limitations in imaging AMPAR vesicles. Efforts to capture the motion of AMPAR vesicles have been hindered by obstacles in receptor tagging (Groc and Choquet, 2020). One approach utilizes chemically inducible dimerization to temporally control the release of exogenous AMPARs (i.e., AMPARs overexpressed from plasmid DNA) from the endoplasmic reticulum into the secretory pathway, followed by tracking AMPAR vesicles as they traverse photobleached sections of dendrite (Hangen et al., 2018). Using this technique, Hangen et al. (2018) found AMPAR vesicles slow down and pause during the influx of calcium, and hypothesized that a calcium-mediated mechanism primes AMPAR vesicles for exocytosis. However, it is unclear whether AMPAR vesicle pausing is directly linked to increased AMPAR exocytosis at or near synapses undergoing LTP. Additional studies have demonstrated that endosomes containing exogenous AMPARs can enter dendritic spines (Esteves da Silva et al., 2017), raising the possibility of direct AMPAR exocytosis into synapses. However, scanning electron micrographs of immunogold-labelled AMPARs fail to reveal a substantial fraction of AMPAR vesicles in dendritic spines (Tao-Cheng et al., 2011). Moreover, imaging exogenous AMPARs tagged with super ecliptic pHluorin (SEP) shows that exocytosis occurs largely at extrasynaptic sites (Lin et al., 2007; Makino and Malinow, 2009; Patterson et al., 2010), favoring a model in which receptors diffuse into synapses after exocytosis.

Much research has been focused on understanding how synapses capture AMPARs as they diffuse through the neuronal membrane (Choquet and Opazo, 2022). After exocytosis, AMPARs diffuse freely in random directions, but this diffusion decreases precipitously at synapses because AMPARs are anchored there by postsynaptic density (PSD) proteins, such as PSD-95 (Tardin et al., 2003; Makino and Malinow, 2009; Opazo et al., 2010). CaMKII-mediated signaling during LTP changes both the composition of proteins in the synapse and posttranslational modifications on AMPARs, further enhancing receptor anchoring (Opazo et al., 2010; Opazo et al., 2012). These observations support a model where AMPARs may not be trafficked to specific loci, but rather diffuse in random directions, only to be concentrated in synapses undergoing LTP as a consequence of their increased residence time. Importantly, crosslinking AMPARs on the neuronal membrane to prevent their diffusion impairs synaptic potentiation in vivo (Penn et al., 2017). Nevertheless, the net distance a receptor can travel via diffusion is limited (Groc and Choquet, 2020). Consequently, the viability of this model depends on the presence of nearby extrasynaptic reservoirs from which synapses can draw AMPARs during LTP. How these reservoirs are established and maintained is not fully understood, but given that synapses can be located hundreds of microns from the cell body, it is probable that receptors are actively transported to these locations.

To address whether and how neurons specify the location to which AMPAR vesicles are delivered, we developed a method to identify vesicles containing AMPAR GluA1 subunits expressed at native levels from endogenous loci and characterize the motion of these vesicles in cultured rat hippocampal neurons. Using this technique, we identify previously undescribed motion behaviors for GluA1-containing vesicles (hereafter referred to as GluA1 vesicles). We show that stimulating neuronal activity with glycine-induced chemical LTP (cLTP) or glutamate uncaging-evoked structural LTP (sLTP) results in the local confinement of GluA1 vesicles in the dendritic shaft. We find that confinement concentrates GluA1 vesicles near sites of stimulation, thereby increasing the size of GluA1 reservoirs near these sites. GluA1 vesicle confinement is the result of stimulation-induced actin polymerization in the dendritic shaft, which changes the rheological properties of the dendritic cytoplasm in a manner that inhibits transport along the length of the dendrite and inhibits diffusion of GluA1 vesicles. Finally, we show that actin polymerization in the dendritic shaft near the sites of stimulation facilitates myosin-mediated transport of GluA1 vesicles from intracellular reservoirs to sites of exocytosis. In sum, our results suggest that neurons enhance the delivery AMPARs to synapses undergoing LTP by restricting the motion of AMPAR vesicles away from these synapses while simultaneously promoting AMPAR exocytosis near these synapses.

## Results

### Neuronal activity confines GluA1 vesicles by disrupting vesicle motion in the dendritic shaft of cultured rat hippocampal neurons

To study the intracellular transport of AMPARs during neuronal activity, we developed a method to tag and sparsely label endogenous AMPAR GluA1 subunits (encoded by *Gria1*) in cultured rat hippocampal neurons, and then identify and track GluA1 vesicles in live-cell time-lapse sequences. We tagged endogenous GluA1 using homology-independent targeted integration (HITI) to circumvent issues associated with overexpressing AMPARs, such as mislocalization and the formation of non-native tetramers (Diering and Huganir, 2016). We selected HaloTag (HT) as the attachment strategy because it provides us with a versatile platform to label GluA1 with bright and photostable Janelia Fluor (JF) dyes (Grimm et al., 2015), as well as other functional dyes, which can be conjugated to the HaloTag ligand (HTL).

HITI is a Cas9-based genetic editing strategy that relies on non-homologous end joining (NHEJ) to insert a donor sequence into a target site (Suzuki et al., 2016). This strategy involves co-transfecting or co-transducing neurons with a donor plasmid containing HaloTag flanked on each side by one copy of the *Gria1* sequence to be targeted by Cas9 and a second plasmid containing *Cas9* (**Figure 1A**). To identify a viable insertion site for HaloTag in the extracellular amino-terminal domain (NTD) of GluA1, we tested five Cas9 target sites and three glycine-serine (GS) linker lengths (**Figure 1-figure supplement 1A-B**). Inserting HaloTag at R280 flanked by 5 amino acid GS linkers results in the highest knock-in efficiency (**Figure 1-figure supplement 1B**). GluA1 edited at this position with HaloTag (GluA1-HT) and labeled with JF_549_-HaloTag ligand (JF_549_-HTL) is concentrated in the dendritic spines, similar to endogenous GluA1 (**Figure 1B**; Craig et al., 1993). Overall, this tagging strategy has a relative knock-in efficiency of 1.5-3%, depending on the method of plasmid delivery (**Figure 1C**). Using adeno-associated virus (AAV)-mediated transduction, we can achieve higher knock-in efficiencies, especially when plasmids are packaged into AAV with an rh10 capsid. Critically, this strategy enables us to label neurons with the necessary density to readily identify edited cells under high magnification, but also with sufficient sparsity that processes from different neurons can be easily distinguished (**Figure 1C**, images).

**Figure 1.**
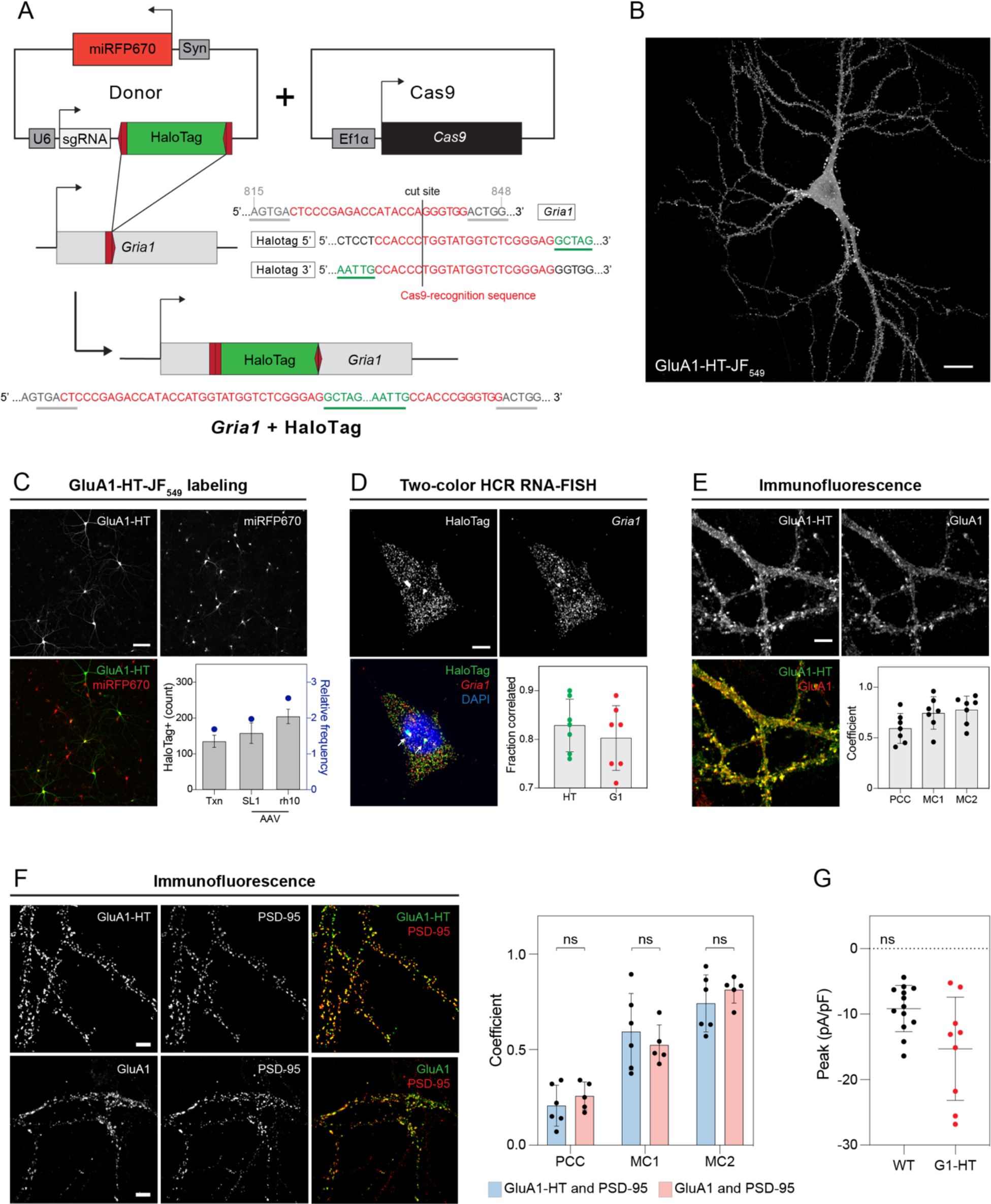
Endogenous GluA1 labeled with HaloTag via homology-independent targeted integration is trafficked to postsynaptic densities and responds to stimulation. (A) Schematic of *Gria1* gene targeting with HaloTag (HT) by homology-independent targeted integration (HITI). A donor construct for HaloTag integration (Donor) is co-transfected into rat hippocampal neurons with a construct expressing Cas9 (Cas9). The donor contains HaloTag flanked on each side by one copy of the *Gria1* sequence to be targeted by Cas9, a single guide RNA (sgRNA), and neuronal transfection marker (e.g., miRFP670). In neurons transfected with both the Cas9 and Donor constructs, Cas9 creates a double strand break in the genomic copy of *Gria1* and also excises the HaloTag sequence from the donor construct. The freed HaloTag sequence can be inserted into the genomic cut site when the double strand break is repaired by non-homologous end joining (NHEJ). (B) Representative confocal image of a cultured rat hippocampal neuron expressing endogenous GluA1 tagged with HaloTag and labeled with JF_549_-HaloTag ligand (GluA1-HT-JF549). Scale bar, 20 μm. (C) Images: representative widefield images of electroporated neurons expressing endogenous GluA1 tagged with HaloTag and labeled with JF_549_-HaloTag ligand (GluA1-HT) or an miRFP670 neuronal transfection marker (miRFP670). Scale bar, 100 μm. Bar graph: efficiencies for HaloTag insertion for electroporation (txn) vs AAV-infection with SL1 or rh10 pseudotyped vectors. Bars represent the mean count and standard deviation for HaloTag positive neurons per dish. Blue dots represent relative efficiencies (HaloTag positive cells divided by cells transfected with both Donor and Cas9). n=6 transfections or infections for each condition. (D) Images: representative confocal images of a two-color hybridization chain reaction RNA fluorescence in situ hybridization (HCR RNA-FISH) labeling experiment with probes targeting HaloTag mRNA (HaloTag) or *Gria1* mRNA (*Gria1*). Arrows indicate active transcription sites (ATS). Scale bar, 5 μm. Bar graph: fraction of HaloTag mRNA spots located within 0.5 μm of *Gria1* mRNA spots and vice versa. Each dot represents a neuron. Correlated localization was used to establish intramolecular labeling between two probe sets (Figure 1**-figure supplement 2A** and **Methods** for details). (E) Images: representative confocal images of a dendrite labeled with anti-HaloTag antibody (GluA1-HT) and anti-GluA1 antibody (GluA1). Scale bar, 5 μm. Bar graph: Pearson correlation coefficients (PCC) and Manders’ coefficients (MC1 and MC2) between HaloTag and GluA1 labeling. Bars represent mean and standard deviation. Each dot represents a neuron. (F) Images: representative confocal images of dendrites labeled with anti-HaloTag antibody (GluA1-HT), anti-GluA1 antibody (GluA1), and anti-PSD-95 antibody (PSD-95) from the same dish. Scale bars, 5 μm. Bar graph: PCCs and MCs between GluA1-HT and PSD-95 labeling vs GluA1 and PSD-95 labeling. Bars represent mean and standard deviation. Each dot represents a neuron. Significance was determined by Mann-Whitney test. (G) Currents elicited by GluA1-HT and unlabeled neurons in response to locally perfused glutamate. Crosshairs represent mean and standard deviation of peak current densities (pA/pF) for each condition (Figure 1**-figure supplement 3C** for current density over time). Each dot represents a single recording. GluA1-HT, n=9; unlabeled, n=13. Significance was determined by Mann-Whitney test. Current densities for GluA1-HT neurons and unlabeled neurons were recorded following a 5 sec 100 µM glutamate stimulation at a –70 mV holding potential.

We first assessed whether HaloTag correctly inserts into GluA1 by comparing the localization of HaloTag mRNA and *Gria1* mRNA using two-color hybridization chain reaction RNA fluorescence in situ hybridization (HCR RNA-FISH) labeling. mRNA labeled by HCR RNA-FISH can be detected and quantified as single puncta (Choi et al., 2018). By using probes conjugated with Alexa Fluor 488 to target HaloTag mRNA and probes conjugated with Alexa Fluor 546 to target *Gria1* mRNA, we find a close correspondence in the localization of these two signals, particularly at active transcription sites (ATS), suggesting that HaloTag inserts into *Gria1* and that *Gria1*-HaloTag is transcribed as a single mRNA molecule (**Figure 1D** and **Figure 1-figure supplement 2A**). We then examined whether HaloTag protein colocalizes with GluA1 protein by immunofluorescence staining. GluA1 labeling and HaloTag labeling are correlated as evidenced by strong Pearson correlation and Manders’ coefficients (**Figure 1E**). To confirm this result, we labeled GluA1 with an antibody that targets an epitope on GluA1 that is disrupted by the HaloTag insertion (**Figure 1-figure supplement 3A**). We observe only weak GluA1 labeling in cells that label for HaloTag, demonstrating that HaloTag correctly inserts into the NTD of GluA1.

Having established a site in *Gria1* where HaloTag can be reliably inserted, we next sought to demonstrate that editing *Gria1* did not dramatically alter its role during neuronal activity by disrupting *Gria1* expression or GluA1 trafficking and function. We first used HCR RNA-FISH to quantify HaloTag mRNA expression versus endogenous *Gria1* expression (**Figure 1-figure supplement 2B-C**). We identified and segmented cells using the far-red-excited and fluorogenic JF_646_-HaloTag ligand (JF_646_-HTL; Grimm et al., 2015) and then detected mRNA spots using FISH-Quant (**Figure 1-figure supplement 2B**; Mueller et al., 2013). We initially found that *Gria1* mRNA counts were higher than HaloTag mRNA counts (**Figure 1- figure supplement 2C**). We hypothesized that this was the result of tagging a single *Gria1* allele due to low knock-in efficiency. To test this possibility, we quantified the number of active transcription sites (ATS) for HaloTag versus *Gria1*. We find that only 34% of knock-ins have two HaloTag ATS, while 79% of cells express *Gria1* from both alleles (**Figure 1-figure supplement 2D**). When we segregated the cells based on ATS number and quantified mRNA counts, we find that there is no significant difference between HaloTag and *Gria1*, demonstrating that the knock-in does not disrupt *Gria1* expression levels (**Figure 1-figure supplement 2D**).

We next assessed if inserting HaloTag into the NTD disrupts trafficking of GluA1 by examining whether GluA1-HT colocalizes with postsynaptic density markers in a manner similar to untagged GluA1. We performed three-color immunofluorescence by labeling neurons with antibodies for GluA1, HaloTag, and PSD-95 (**Figure 1F**). Although untagged GluA1 and GluA1-HT have weak correlations with PSD-95 in terms of signal intensity (Pearson correlation), both have strong colocalization in terms of their overlapping signal (Manders’ overlap; **Figure 1F**). We detected no significant difference in the colocalization between GluA1 and PSD-95 versus GluA1-HT and PSD-95 (**Figure 1F**). GluA1-HT also colocalizes with Homer1 in a manner similar to untagged GluA1 (**Figure 1-figure supplement 3B**). These results suggest that transport of GluA1 to postsynaptic densities is not significantly disrupted by the insertion of HaloTag.

Because we inserted HaloTag closer to the ligand binding domain than in previously reported tag-GluA1 fusions (which place the tag immediately after the signal peptide; Shi et al., 1999), we tested whether HaloTag disrupts the function of GluA1. We performed whole-cell voltage-clamp recordings on unlabeled and labeled neurons, and measured Na^+^ current density in response to a locally perfused application of 100 µM glutamate (**Figure 1G** and **Figure 1-figure supplement 3C**). In order to exclusively record conductance through AMPAR, current measurements were performed at −70 mV holding potential and in the presence of the voltage gated sodium channel (Nav) blocker tetrodotoxin (TTX) and the NMDAR inhibitor AP5 with 2 mM Mg^2+^. To determine AMPAR current density, recordings were normalized via cell capacitance (pA/pF). We controlled for differences that might arise due to viral transduction and dye labeling by patching unlabeled neurons from coverslips that also contained HaloTag-edited and JF_549_-HTL-labeled neurons. We find no significant difference in peak current density in response to glutamate between untagged GluA1 and GluA1-HT (**Figure 1G**).

After establishing a method to tag GluA1 in cultured rat hippocampal neurons and demonstrating that the tag did not significantly alter the trafficking or function of GluA1, we next developed a protocol to label and image GluA1-HT so that we could track the motion of GluA1-HT vesicles in the dendritic shaft. Labeling GluA1-HT with JF dyes results in saturated signal in dendrites stemming from both intracellular and surface GluA1-HT (**Figure 1B**), making it difficult to detect single GluA1-HT vesicles. To bypass the limitations of previous approaches used to reduce signal saturation (e.g., photobleaching; Hangen et al., 2018), we developed a block-and-chase protocol to achieve sparse labeling of de novo synthesized endogenous GluA1 (**Figure 2A**). To avoid visualizing pre-existing GluA1-HT, GluA1-HT was first labeled with a saturating concentration of JF_646_-HTL in the presence of the translation inhibitor cycloheximide (CHX). Next, JF_646_-HTL and CHX were washed off the neurons and, after an incubation period to allow GluA1-HT translation to recover, de novo synthesized GluA1-HT was labeled with JF_549_-HTL. Because JF_549_-HTL labels newly synthesized GluA1-HT, signal detected by fluorescence microscopy is dramatically reduced in this fluorescence channel, allowing us to identify sparse, punctate GluA1-HT-JF549 conjugates (**Video 1**). We then distinguished GluA1- HT vesicles from surface GluA1-HT based on the fluorescence bleaching characteristics of punctate signal (**Figure 2-figure supplement 1**), its localization in the dendritic shaft, and its motion type. After identifying GluA1-HT vesicles, we used single-particle tracking (SPT) analysis to reconstruct trajectories for their motion (**Figure 2B**, and **Methods** for details; Jaqaman et al., 2008, Liu et al., 2018). We then used hidden Markov modeling with Bayesian model selection (HMM-Bayes) to characterize the motion type for each GluA1-HT vesicle trajectory (i.e., active transport versus diffusion) and determine important motion parameters (i.e., velocity and diffusion coefficient; **Figure 2C**, and **Figure 2-figure supplement 2** for details).

**Figure 2.**
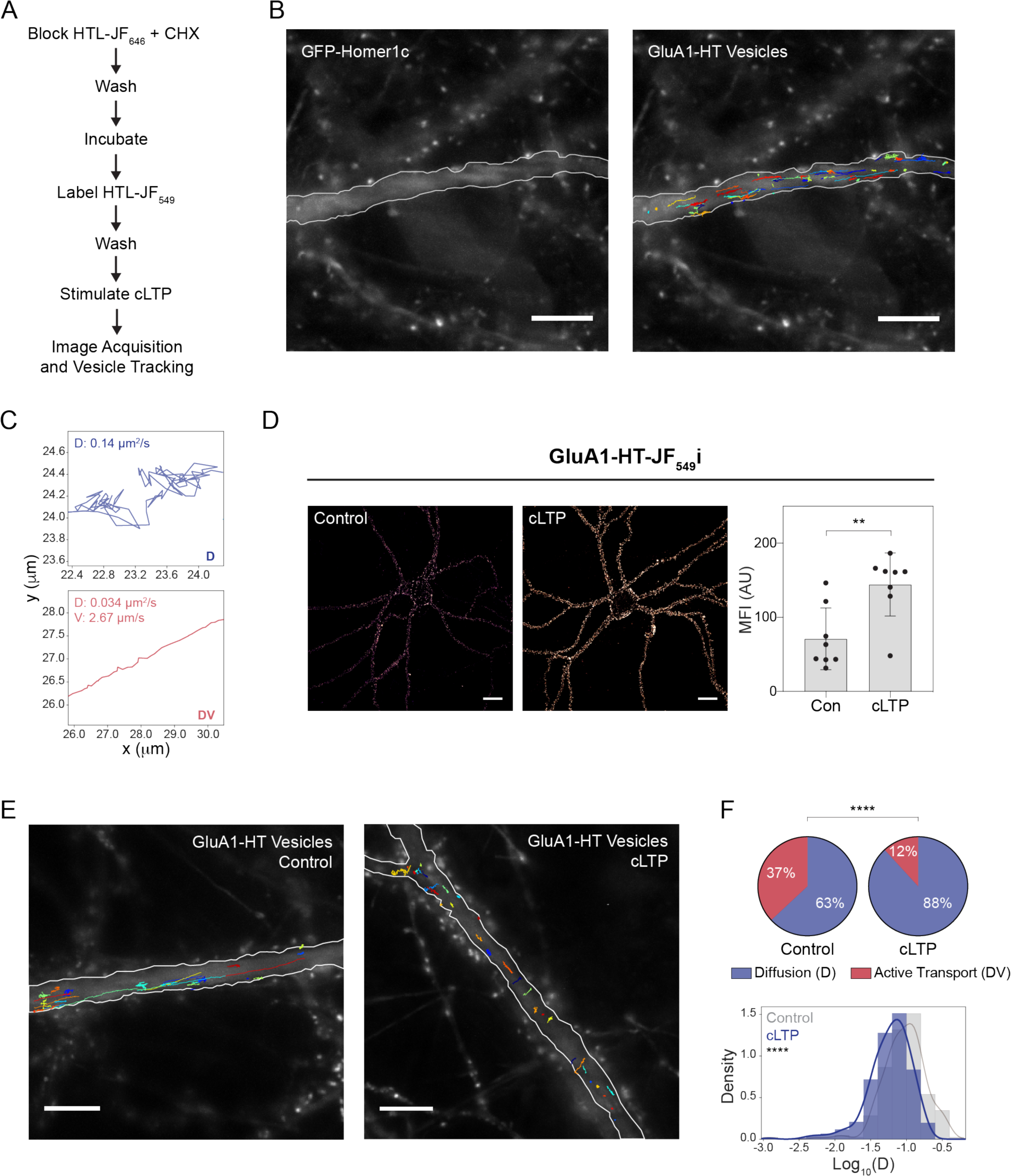
Chemical LTP induction reduces active transport and diffusion of GluA1-HT vesicles in the dendritic shaft. (A) Experimental workflow to achieve sparse GluA1-HT labeling for GluA1-HT vesicle identification and single particle tracking (SPT) analysis. GluA1-HT vesicles were identified post-acquisition based on localization in the dendritic shaft, motion type, and bleaching rate of particles (Figure 2**-figure supplement 1** for details). (B) Representative widefield image of exogenous GFP-Homer1c used to determine dendritic shaft boundaries (GFP-Homer1c) and trajectories of GluA1-HT-JF549 vesicles overlaid on the dendrite (GluA1-HT Vesicles). Scale bar, 10 μm. (C) Exemplar GluA1-HT vesicle trajectories with diffusive motion (D) and active transport (DV) inferred by HMM-Bayes analysis. (D) Images: representative confocal images of the selective labeling of GluA1-HT on neuronal membranes using a membrane impermeable version of the JF_549_-HaloTag ligand (JF_549_i-HTL: “i” for impermeable) without treatment (Control) and after cLTP (cLTP). This experiment demonstrates cLTP stimulates the trafficking of GluA1-HT to the membranes of neurons. Scale bars, 20 μm. Bar graph: mean fluorescence intensities (MFI) of JF_549i_-HTL-labeled GluA1-HT without treatment and after cLTP. Bars represent mean and standard deviation, **p=0.0047 by Mann-Whitney test. Each dot represents a neuron. (E) GluA1-HT vesicle trajectories in a dendritic shaft with no treatment (Control) and during cLTP induction (cLTP). Trajectories were overlaid on widefield images of GFP-Homer1c, which were used to determine the boundaries of the dendritic shaft in each experiment. Scale bar, 10 μm. (F) Pie charts: fractions of vesicles exhibiting active transport (DV) or diffusion (D) in dendritic shafts with no treatment (Control) and during cLTP induction (cLTP). ****p<0.0001 by Mann- Whitney test. n=12-14 experiments for each condition. Histogram: distributions of diffusion coefficients for GluA1-HT vesicles without treatment and during cLTP induction. ****p<0.0001 by Kolmogorov-Smirnov test. n=232-383 trajectories for each condition. Line represents the probability density function of each histogram estimated by kernel density estimation (KDE).

Having established a pipeline to label, track, and analyze GluA1-HT vesicles, we sought to evaluate whether the motion of GluA1-HT vesicles could be modulated by neuronal activity. Glycine-induced chemical LTP (cLTP) is an established method of stimulating neuronal activity and increasing the surface expression of GluA1 in cultured neurons (Molnár, 2011). We hypothesized that cLTP induction could change the motion of GluA1-HT vesicles in the dendritic shaft to support increased GluA1-HT exocytosis during neuronal activity. To test this hypothesis, we first validated that cLTP could stimulate increased expression of GluA1-HT on neuronal surfaces by selectively labeling membrane-bound GluA1-HT with a membrane impermeable variant of JF_549_-HTL termed JF_549i_-HaloTag ligand (JF_549i_-HTL; “i” for impermeant; Xie et al., 2017) after cLTP or in unstimulated neurons (**Figure 2D**). We find increased GluA1-HT-JF549i mean fluorescence intensity (MFI) in neurons stimulated with cLTP versus unstimulated control neurons (**Figure 2D**), demonstrating that cLTP increases trafficking of GluA1-HT to neuronal surfaces. We then imaged and tracked GluA1-HT vesicles in dendrites during cLTP induction and in dendrites with no treatment. GluA1-HT vesicles in the dendrites of cLTP-stimulated neurons exhibit clear qualitative differences in motion versus vesicles in the dendrites of unstimulated control neurons (**Figure 2E**, **Video 2** and **Video 3**). Most strikingly, we observed a loss in long-range motion along the length of the dendritic shaft in cLTP- stimulated neurons (**Figure 2E**). When HMM-Bayes analysis is applied to characterize trajectories collected during cLTP induction, we find a significant decrease in the fraction of GluA1-HT vesicles undergoing active transport and an increase in GluA1-HT vesicles exhibiting diffusion (**Figure 2F**, pie charts). The diffusion coefficients (D) for GluA1-HT vesicles undergoing diffusion are also significantly reduced by cLTP induction (**Figure 2F**, histogram).

In addition, vesicles exhibit increased subdiffusion (i.e., constrained diffusion due to molecular crowding or interactions) in response to cLTP induction (**Figure 2-figure supplement 3**).

These observations demonstrate that the overall motion of GluA1-HT vesicles is inhibited by cLTP, suggesting that vesicle motion is locally confined (defined in this work as the restriction of vesicle motion away from its initial position). We reasoned that if cLTP spatially confines GluA1-HT vesicles then it should, in addition to decreasing the fraction of vesicles undergoing active transport, prevent diffusing vesicles from transitioning to active transport and leaving their local regions. A critical feature of HMM-Bayes analysis is the ability to infer multiple motion states from a single trajectory and determine state-transition probabilities (**Figure 2-figure supplement 4A-B**; Monnier et al., 2015). Using HMM-Bayes analysis, we identified GluA1-HT vesicles that stochastically switch motion states from active transport to diffusion and vice versa under unstimulated control conditions (**Figure 2-figure supplement 4A**). These vesicles switch between active transport and diffusion with approximately the same probability (**Figure 2-figure supplement C**, Control, kDV-D vs kD-DV). By contrast, GluA1-HT vesicles have a greater probability of switching from active transport to diffusion than from diffusion to active transport during cLTP (**Figure 2-figure supplement 4B-C**, cLTP, kDV-D vs kD-DV). Furthermore, GluA1-HT vesicles undergoing diffusion have a high probability to continue diffusing (**Figure 2-figure supplement 4B-C**, cLTP, k_D-D_). Taken together, our observations suggest that cLTP induction results in the local confinement of GluA1-HT vesicles.

### Local induction of neuronal activity concentrates GluA1 vesicles near the site of activity by disrupting GluA1 vesicle motion

We hypothesized that confinement could be a mechanism to increase the intracellular reservoir of GluA1-HT near sites of neuronal activity. However, because cLTP induction results in neuron-wide activity, we were unable to determine if changes to GluA1-HT vesicle motion are limited to sites of activity. Consequently, we tested whether the stimulation of activity at a specific location would alter GluA1-HT vesicle motion and concentration near that location. Single-photon (1P) 4-Methoxy-7-nitroindolinyl-caged-L-glutamate (MNI) uncaging can be used to stimulate neuronal activity in a desired region (Ellis-Davies, 2007). We used 1P MNI uncaging with a 405 nm laser at 2 Hz for 50 seconds to induce structural LTP (sLTP) at single dendritic spines (**Figure 3A**). We selected this LTP protocol because in addition to stimulating calcium transients, previous studies have demonstrated that sLTP changes both the structure of stimulated spines and the concentration of AMPARs in stimulated spines – two important markers for sustained neuronal activity during LTP in hippocampal slices (Matsuzaki et al., 2004; Bosch et al., 2014). After sLTP, we observed a significant increase in the area of the stimulated spine head in dishes containing MNI, but not in dishes without MNI, indicating that this protocol induces sustained local activity (**Figure 3B**).

**Figure 3.**
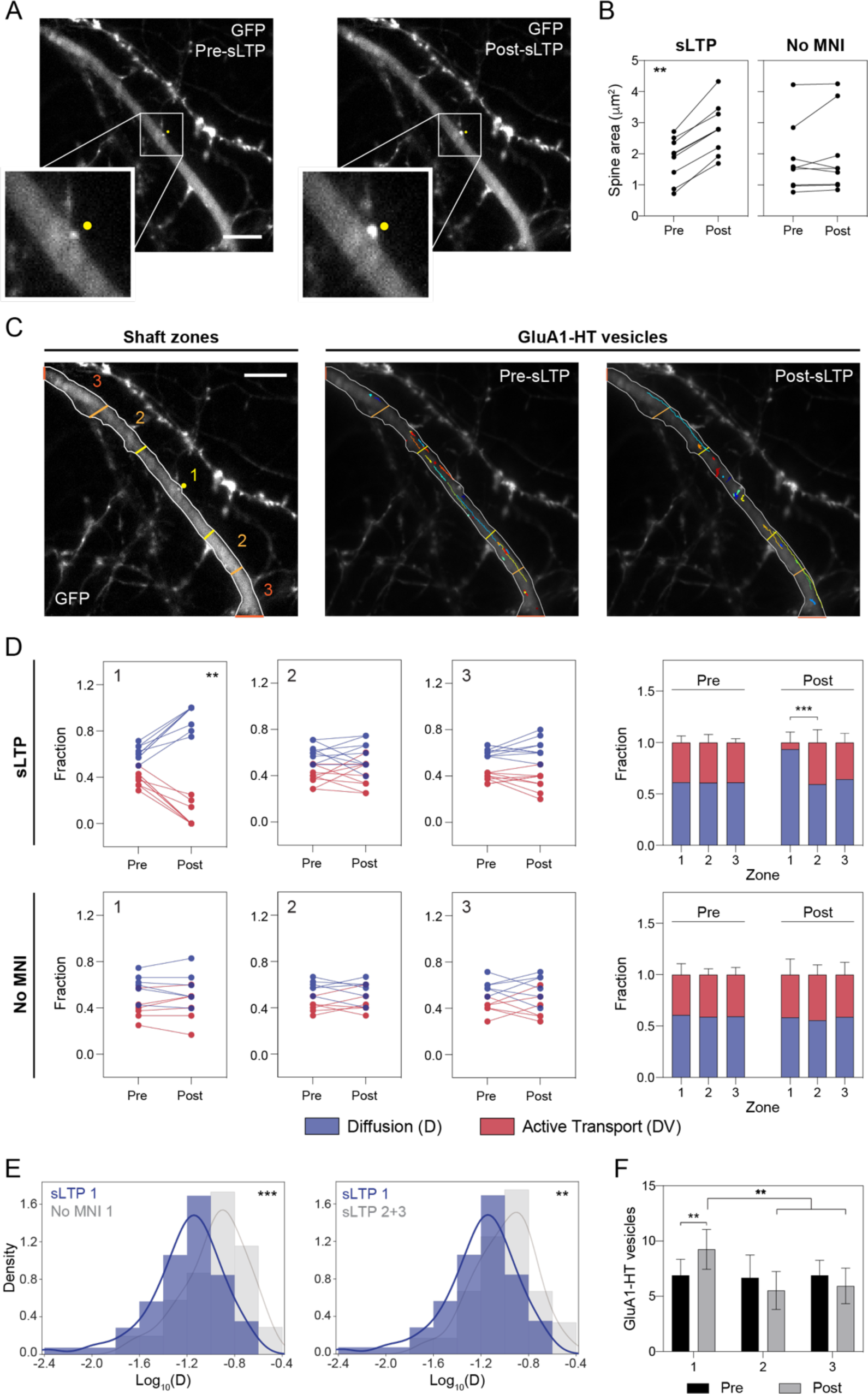
Structural LTP reduces active transport and diffusion of GluA1-HT vesicles proximal to the site of neuronal activity. (A and B) Dendritic spines before and after sLTP. (A) Representative widefield images of a dendrite expressing GFP before sLTP (Pre-sLTP) and after sLTP (Post-sLTP). Yellow dot indicates the site of uncaging. Scale bar, 10 μm. Inset: magnified images of the dendritic spine at the site of uncaging. (B) Line graphs of spine head areas before and after sLTP (sLTP) and stimulation control experiments with no MNI (No MNI). sLTP, **p=0.0039 by Wilcoxon matched pairs test. n=9 for each condition. sLTP was achieved by stimulating a single dendritic spine with a 405 nm laser at 2 Hz for 50 seconds. (C) Shaft zones: representative widefield image of a dendrite transfected with GFP that has been separated into 3 equal zones based on proximity to the uncaged spine (yellow dot) to distinguish proximal and distal areas. The length of the depicted dendrite is 90 μm, so Zone 1 is composed of the 30 μm region around the uncaged spine while Zone 2 is composed of the two 15 μm regions flanking Zone 1, and so forth. Scale bar, 10 μm. GluA1-HT vesicles: GluA1-HT vesicle trajectories before sLTP (Pre-sLTP) and after sLTP (Post-sLTP). (D) Line graphs: fractions of GluA1-HT vesicles exhibiting active transport (DV) or diffusion (D) before or after sLTP in each zone as depicted in Figure 3C. Top row of graphs are of sLTP experiments in dishes containing MNI (sLTP) while the bottom row of graphs are stimulation control experiments in dishes with no MNI (No MNI). sLTP Zone 1, **p=0.0039 by Wilcoxon matched pairs test. Bar graphs: mean fractions of GluA1-HT vesicles exhibiting active transport or diffusion before or after sLTP in each zone. Error bars represent standard deviation. Top graph is of sLTP experiments with MNI while the bottom graph is of stimulation control experiments with no MNI. sLTP: Zone 1 post vs Zone 2 post, ***p=0.0001 by Mann-Whitney test. n=6-9 experiments for each condition. (E) Left: distributions of diffusion coefficients for GluA1-HT vesicles in Zone 1 after sLTP (sLTP 1) versus GluA1-HT vesicles in Zone 1 after stimulation in the absence of MNI (No MNI 1). ***p=0.0001. Right: distribution of diffusion coefficients for GluA1-HT vesicles in Zone 1 (sLTP 1) versus Zone 2+3 (sLTP 2+3) after sLTP. **p=0.0032. Significance was determined by Kolmogorov-Smirnov test. n=52-70 trajectories for each condition. Line represents the probability density function of each histogram estimated by kernel density estimation (KDE). (F) Bar graph of adjusted GluA1-HT vesicle counts before or after sLTP in each zone. Error bars represent standard deviation. Zone 1: pre vs post, **p=0.0050. Zone 1 post vs Zone 2 post, **p=0.0017. Zone 1 post vs Zone 3 post, ***p=0.0005. Significance was determined by Mann- Whitney test. Vesicle count was adjusted for photobleaching.

We then examined whether sLTP changed GluA1-HT vesicle motion proximal to the site of stimulation. To define regions proximal and distal to the site of sLTP, we used GFP to define the borders of the dendritic shaft and separated the shaft longitudinally (i.e., along the length of the dendrite) into 3 equal zones based on proximity to the site of uncaging (**Figure 3C**, Shaft zones). We then assessed the different types of motion that GluA1-HT vesicles exhibit in each zone before and after sLTP (**Figure 3C**, GluA1-HT vesicles, and **Video 4**). We find that sLTP results in reduced active transport and increased diffusion in GluA1-HT vesicles in Zone 1 but not Zone 2 or Zone 3 (**Figure 3D**, sLTP), indicating that stimulation disrupts active transport proximal but not distal to the site of neuronal activity. By contrast, sLTP stimulation in the absence of MNI failed to significantly alter the fraction of vesicles exhibiting active transport both proximal and distal to the site of stimulation (**Figure 3D**, No MNI), demonstrating that changes in vesicle movement after sLTP are not the result of laser exposure. The average diffusion coefficient of GluA1-HT vesicles is also significantly reduced in Zone 1 versus Zone 2 and Zone 3 after sLTP, but only when sLTP stimulation is performed in the presence of MNI (**Figure 3E** and **Figure 3-figure supplement 1A**). Furthermore, GluA1-HT vesicles in Zone 1 imaged up to 10 minutes after the cessation of sLTP stimulation exhibited similar motion behaviors to those imaged immediately after the cessation of sLTP (**Figure 3-figure supplement 1B**), indicating that the vesicles are confined.

To further test whether vesicles are confined to the site of neuronal activity, we used HMM-Bayes analysis to determine the probabilities that vesicles switch between active transport and diffusion in each zone during sLTP. We find that sLTP increases the probability that multistate GluA1-HT vesicles undergoing active transport in Zone 1 switch to and stay in diffusion, but not vice versa (**Figure 3-figure supplement 2**, sLTP Zone 1), in a manner dependent on the presence of MNI (**Figure 3-figure supplement 2**, No MNI). By contrast, multistate vesicles in Zone 2 and Zone 3 have similar probabilities of switching from active transport to diffusion and vice versa after sLTP (**Figure 3-figure supplement 2**, sLTP Zone 2+3). These observations demonstrate that multistate GluA1-HT vesicles proximal to the site of stimulation have low probabilities of being transported away from the site of stimulation.

These results demonstrate that sLTP confines GluA1-HT vesicles near the site of stimulation, but it is unclear whether local confinement actually results in an increased number of vesicles near these sites. After adjusting the number of vesicles for photobleaching (**Methods** for details), we find a significant increase in the number of GluA1-HT vesicles in Zone 1 (but not Zones 2 or 3) after sLTP (**Figure 3F**). Combined, these observations demonstrate that neurons use confinement as a mechanism to increase the number of vesicles near the sites of neuronal activity.

### Confinement of GluA1 vesicles during neuronal activity is mediated by F-actin-induced molecular crowding in the dendritic shaft

Having demonstrated that sLTP concentrates GluA1-HT vesicles in the dendritic shaft near the sites of neuronal activity by reducing the motion of GluA1-HT vesicles, we sought to determine the molecular mechanisms involved in disrupting vesicle motion. We hypothesized that actin polymerization in the dendritic shaft might be involved because actin polymerization is induced by calcium signaling (Okamato et al., 2009) and plays a critical role in the expansion of dendritic spines during sustained neuronal activity (Fukazawa et al., 2003). Moreover, recent studies have observed the rearrangement of F-actin networks in the dendritic shaft during neuronal activity (Schätzle et al., 2018; Lavoie-Cardinal et al., 2020), and found that F-actin networks can reposition lysosomes in neurites (van Bommel et al., 2019; Katrukha et al., 2017). Importantly, F-actin networks formed in response to sLTP are persistent (Okamoto et al., 2004), and therefore could be a mechanism to confine GluA1-HT vesicles even after the cessation of sLTP stimulation.

To determine whether actin polymerization occurs in the dendritic shaft of cultured rat hippocampal neurons during neuronal activity, we tested whether cLTP would lead to the redistribution of F-tractin, an actin binding peptide derived from rat inositol 1,4,5-triphosphate 3- kinase (IPTKA) that is used as a marker for F-actin (Schell et al., 2001). We transduced neurons with a plasmid containing F-tractin-mNeongreen (tractin) and imaged before and during cLTP induction (**Figure 4A**). Prior to cLTP, tractin is diffusely distributed in the dendritic shaft and concentrated in dendritic spines (**Figure 4A**, Control). During cLTP induction, tractin in the dendritic shaft redistributes into a network of filaments (**Figure 4A**, cLTP). To eliminate the possibility that qualitative changes in the distribution of tractin are due to gross morphological changes in the dendrite, neurons were also transduced with a plasmid expressing tdTomato (**Figure 4-figure supplement 1A**). tdTomato did not dramatically redistribute during cLTP, indicating that the change in tractin localization is due to increased F-actin. To quantify actin polymerization, we determined the length of the filamentous tractin network by thresholding and skeletonizing tractin signal in the dendritic shaft, and then measuring the length of the skeleton (**Figure 4-figure supplement 2**). When the same thresholding and measurement parameters are applied to a dendritic shaft before and during cLTP, we find a significant increase in the length of the tractin skeleton during treatment (**Figure 4A**, Skeleton length). Moreover, we find the mean fluorescence intensity (MFI) of the tractin skeleton is significantly increased after cLTP (**Figure 4A**, MFI).

**Figure 4.**
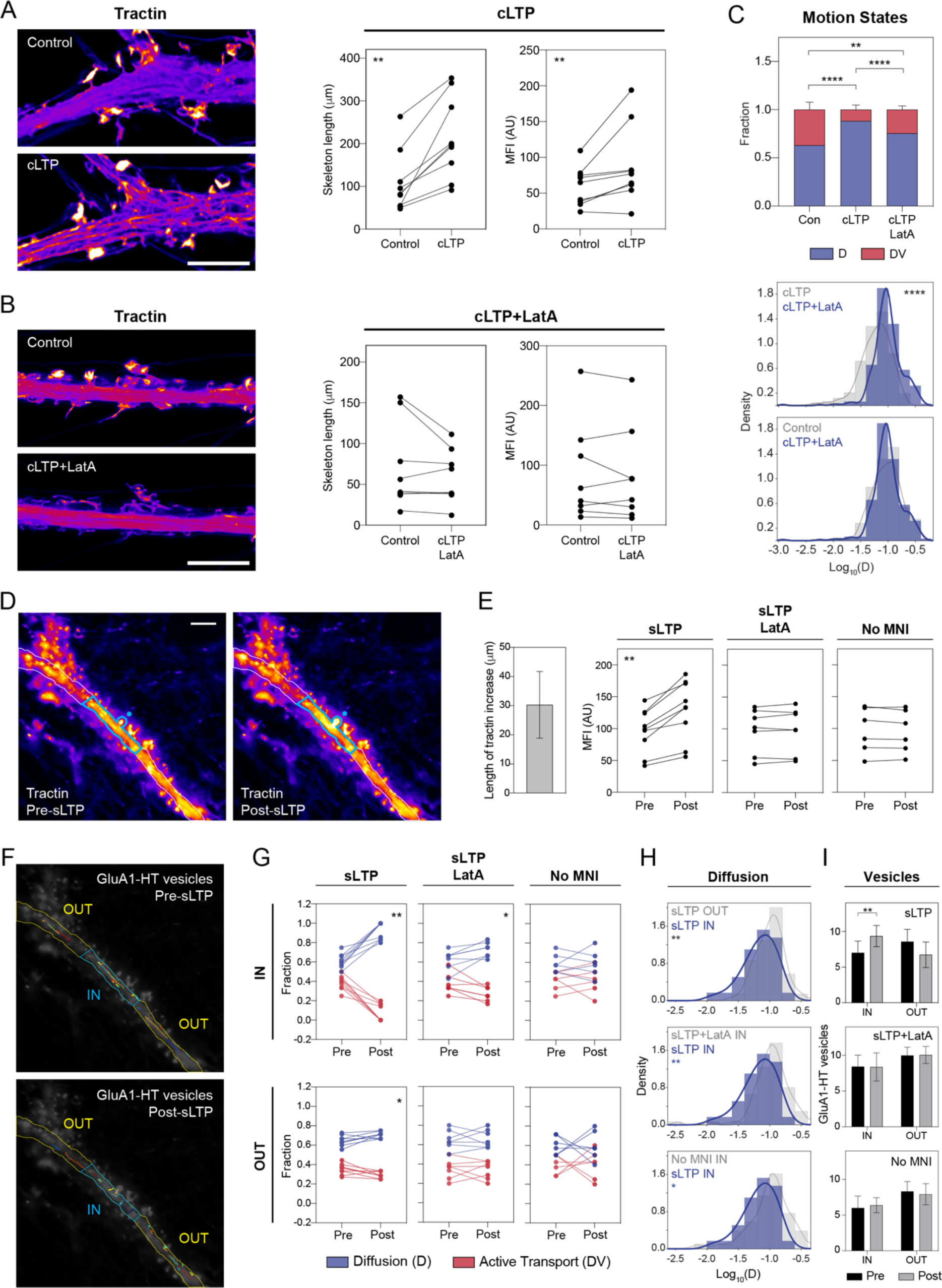
Reduced motion of GluA1-HT vesicles during neuronal activity is mediated by actin polymerization in the dendritic shaft. (A and B) Actin polymerization in the dendritic shaft in response to cLTP induction. (A) Images: representative Airyscan images of F-tractin-mNeongreen (tractin) in a dendrite before treatment (Control) and during cLTP (cLTP). Scale bar, 5 μm. Graphs: combined length of tractin filaments (Skeleton length) and tractin mean fluorescence intensity (MFI) before treatment and during cLTP in the same dendritic shafts. Skeleton length, **p=0.0038; MFI, **p=0.0078. (B) Images: representative Airyscan images of tractin in a dendrite before treatment (Control) and during cLTP in the presence of Latrunculin A (LatA; cLTP+LatA). Scale bar, 5 μm. Graphs: Skeleton length and MFI before treatment and during cLTP+LatA in the same dendritic shaft. Significance was determined Wilcoxon matched pairs test, n=8-9 for each condition. Skeleton length was determined as described in Figure 4**-figure supplement 2**. (C) Bar graph (Motion States): fraction of GluA1-HT vesicles exhibiting active transport (DV) or diffusion (D) in dendritic shafts without treatment (Con), during cLTP (cLTP), and during cLTP in the presence of LatA (cLTP+LatA). Con vs cLTP, ****p<0.0001; Con vs cLTP+LatA, **p=0.0012; cLTP vs cLTP+LatA, ****p<0.0001. Significance was determined by Mann- Whitney test. n=9-14 experiments for each condition. Histograms: distributions of diffusion coefficients for GluA1-HT vesicles in dendritic shafts during cLTP+LatA versus cLTP (top) or Control (bottom). cLTP vs cLTP+LatA, ****p<0.0001 by Kolmogorov-Smirnov test. n=232-383 trajectories for each condition. Line represents the probability density function of each histogram estimated by kernel density estimation (KDE). (D and E) Actin polymerization in the dendritic shaft in response to sLTP. (D) Representative widefield images of tractin before sLTP (pre-sLTP) and after sLTP (post-sLTP). Blue dot represents the site of uncaging. Blue outline denotes area with increased tractin signal after stimulation. Scale bar, 10 μm. (E) Bar graph: mean length (along the length of the dendrite) of the regions proximal to uncaged spines where increased tractin MFI was observed after sLTP. Error bar represents standard deviation. Line graphs: mean fluorescence intensity (MFI) of tractin in dendrites before and after sLTP (sLTP), sLTP in the presence of LatA (sLTP+LatA), and sLTP stimulation in the absence of MNI (No MNI). In sLTP experiments, the MFI for tractin was measured only in the area where there was an increase in tractin intensity. In sLTP+LatA and No MNI experiments, tractin intensity was measured across the entire length of the dendrite because there were no specific regions where an increase in tractin intensity could be observed. sLTP, **p=0.0039 by Wilcoxon matched pairs test. n=6-9 experiments for each condition. (F-H) sLTP-mediated changes in GluA1-HT vesicle motion in regions of dendritic shafts with increased F-actin. (F) GluA1-HT vesicle trajectories before sLTP (Pre-sLTP) and after sLTP (Post-sLTP) inside (IN) and outside (OUT) the region where there was increased actin polymerization post-sLTP. The region in which actin polymerization occurred was determined by separating the region where tractin intensity increased from the regions where it did not. (G) Line graphs of fractions of GluA1-HT vesicles exhibiting active transport (DV) or diffusion (D) before and after sLTP (sLTP), sLTP in the presence LatA (sLTP+LatA), and sLTP stimulation in the absence of MNI (No MNI) inside (IN) and outside (OUT) regions of actin polymerization. Because we did not detect significant increases in actin polymerization for sLTP+LatA and No MNI, IN regions for these conditions were determined by isolating the 30 μm region flanking the uncaging site, which was the mean length of the region where actin polymerization occurred after sLTP (Figure 4E). IN: sLTP, **p=0.0039; sLTP+LatA, *p=0.0312. OUT: sLTP, *p=0.0234. Significance was determined by Wilcoxon matched pairs test. n=6-9 experiments for each condition. (H) Top: distributions of diffusion coefficients for GluA1-HT vesicles in regions with actin polymerization (sLTP IN) versus regions without actin polymerization (sLTP OUT) after sLTP. **p=0.0078. Middle: distributions of diffusion coefficients for GluA1-HT vesicles in regions with actin polymerization after sLTP versus in equivalent regions after sLTP in the presence of LatA (sLTP+LatA IN). **p=0.0026. Bottom: distributions of diffusion coefficients for GluA1-HT vesicles in regions with actin polymerization after sLTP versus in equivalent regions after stimulation in the absence of MNI (No MNI IN). *p=0.0116. Significance was determined by Kolmogorov-Smirnov test. n=53-70 trajectories for each condition. Lines represent the probability density function of each histogram estimated by kernel density estimation (KDE). (I) Bar graphs of adjusted vesicle counts inside (IN) or outside (OUT) regions with actin polymerization after sLTP (sLTP), sLTP in the presence of LatA (sLTP+LatA), or stimulation in the absence of MNI (No MNI). Error bars represent standard deviation. sLTP: IN, **p=0.0081 by Mann-Whitney test. n=6-9 experiments for each condition.

To confirm that the change in tractin distribution during cLTP induction is the result of actin polymerization, we imaged tractin before and during cLTP in dendrites that were also treated with latrunculin A (LatA), an inhibitor of actin polymerization (**Figure 4B** and **Figure 4- figure supplement 1B**). Treating neurons with LatA not only prevents redistribution of tractin during cLTP, but also reduces the intensity of tractin signal in dendritic spines, demonstrating that actin polymerization is inhibited (**Figure 4B**, images). We find no significant increase in the length of the tractin skeleton (**Figure 4B**, Skeleton length) or tractin MFI (**Figure 4B**, MFI) after quantification, and in some dendrites find that the tractin skeleton length decreases after cLTP stimulation in the presence of LatA. These findings demonstrate that cLTP treatment results in actin polymerization and thus the redistribution of tractin in the dendritic shaft.

We then examined whether cLTP-induced actin polymerization results in the confinement of GluA1-HT vesicles by testing whether blocking actin polymerization with LatA prevented cLTP-mediated changes in GluA1-HT vesicle mobility. The loss of active transport exhibited by GluA1-HT vesicles during cLTP is strongly, but not completely, inhibited by LatA treatment (**Figure 4C**, bar graph). This observation suggests that while F-actin plays a role in disrupting active transport in the dendritic shaft during neuronal activity, other mechanisms also contribute to the loss of motion. In addition, treatment with LatA during cLTP induction abrogates the effect of cLTP on GluA1-HT vesicle diffusion (**Figure 4C**, top histogram), resulting in diffusion coefficients that are similar to those for GluA1-HT vesicles under control conditions (**Figure 4C**, bottom histogram, and **Figure 2F**, histogram, to compare cLTP to control conditions). Lastly, multistate GluA1-HT vesicles have roughly equal probabilities of switching between active transport and diffusion and vice versa when treated with LatA during cLTP, similar to the state-transition probabilities exhibited by GluA1-HT vesicles under unstimulated conditions (**Figure 2-figure supplement 4C**, cLTP+LatA). These findings are consistent with the hypothesis that actin polymerization in the dendritic shaft confines GluA1- HT vesicles.

We next sought to determine whether sLTP-mediated changes in local actin networks play a role in positioning vesicles near sites of stimulation. First, we tested if sLTP leads to increased F-actin in the dendritic shaft near the spine targeted by glutamate uncaging. Although the widefield microscope used for 1P MNI uncaging lacked sufficient resolution for us to quantify actin skeletons, we observed a significant increase in tractin MFI in spines stimulated with sLTP and in the dendritic shaft proximal to these spines, reflecting increased actin polymerization at these locations (**Figure 4D**, blue outline and **Figure 4E**, sLTP). Tractin signal increases in an approximately 30 μm longitudinal section of dendritic shaft (i.e., 30 μm along the length of the dendrite) surrounding the sLTP-stimulated spine (**Figure 4E**, bar graph). The increase in tractin MFI during sLTP is dependent on actin polymerization (**Figure 4E**, sLTP+LatA) and is not an artifact of photostimulation (**Figure 4E**, No MNI).

Having found that sLTP increases actin polymerization in the dendritic shaft proximal to the uncaging site, we tested whether sLTP-mediated actin polymerization confined GluA1-HT vesicles by tracking GluA1-HT vesicles before and after sLTP and characterizing their motion inside and outside the area of actin polymerization (**Figure 4F** and **Video 5**). We find that sLTP significantly reduces the fraction of vesicles undergoing active transport in regions of dendritic shafts with actin polymerization (**Figure 4G**, sLTP IN). Furthermore, the reduction of active transport is more substantial inside the region of actin polymerization than outside the region of actin polymerization (**Figure 4G**, sLTP OUT). The effect of sLTP on GluA1-HT vesicle motion is strongly, but not completely, disrupted by LatA treatment (**Figure 4G**, sLTP+LatA) and is not an artifact of repeated laser illumination (**Figure 4G**, No MNI). sLTP also decreased GluA1-HT vesicle diffusion inside, but not outside, regions of actin polymerization in a manner dependent on actin polymerization (**Figure 4H**). In addition, multistate GluA1-HT vesicles undergoing active transport inside, but not outside, regions of actin polymerization during sLTP have an increased probability of switching to diffusion (**Figure 4-figure supplement 3**, IN vs OUT).

LatA treatment during sLTP, by contrast, results in roughly equal state-transition probabilities between active transport and diffusion inside regions of actin polymerization, similar to the state-transition probabilities of multistate GluA1-HT vesicles during sLTP stimulation in the absence MNI (**Figure 4-figure supplement 3**, LatA). Importantly, sLTP results in increased numbers of GluA1-HT vesicles in regions in the dendritic shaft where actin polymerization occurs in a manner that is dependent on actin polymerization (**Figure 4I**). These observations demonstrate that the local confinement and concentration of GluA1-HT vesicles in the dendritic shaft near the site of sLTP is the result of sLTP-induced actin polymerization in the same region.

Having established actin polymerization as the mechanism that mediates stimulation-dependent GluA1-HT vesicle confinement, we sought to determine whether F-actin perturbed vesicle motion through direct or indirect interactions. AMPARs interact with myosins Va, Vb, and VI (Correia et al., 2008, Wang et al., 2008, Nash et al., 2010, Esteves da Silva et al., 2015), which are involved in the calcium-dependent, short-range recruitment of AMPARs in endosomes to and from dendritic spines. We hypothesized that interactions between the intracellular domain of GluA1 and myosin V and/or VI may enable myosin to anchor GluA1-HT vesicles to actin during neuronal activity, disrupting active transport and limiting diffusion. We tested this possibility by inhibiting myosin V and VI using a myosin inhibitor cocktail composed of MyoVin-1, Pentabromopseudilin (PBP), and 2,4,6-Triiodophenol (TIP). We find no significant difference in the fractions of GluA1-HT vesicles exhibiting active transport during cLTP and sLTP versus cLTP and sLTP in the presence of myosin inhibitors (**Figure 5A**). Likewise, inhibition of myosin did not alter the diffusion coefficients of GluA1-HT vesicles during cLTP or sLTP (**Figure 5B**). Based on these observations, we conclude that F-actin disrupts GluA1-HT vesicle motion independent of myosin activity.

**Figure 5.**
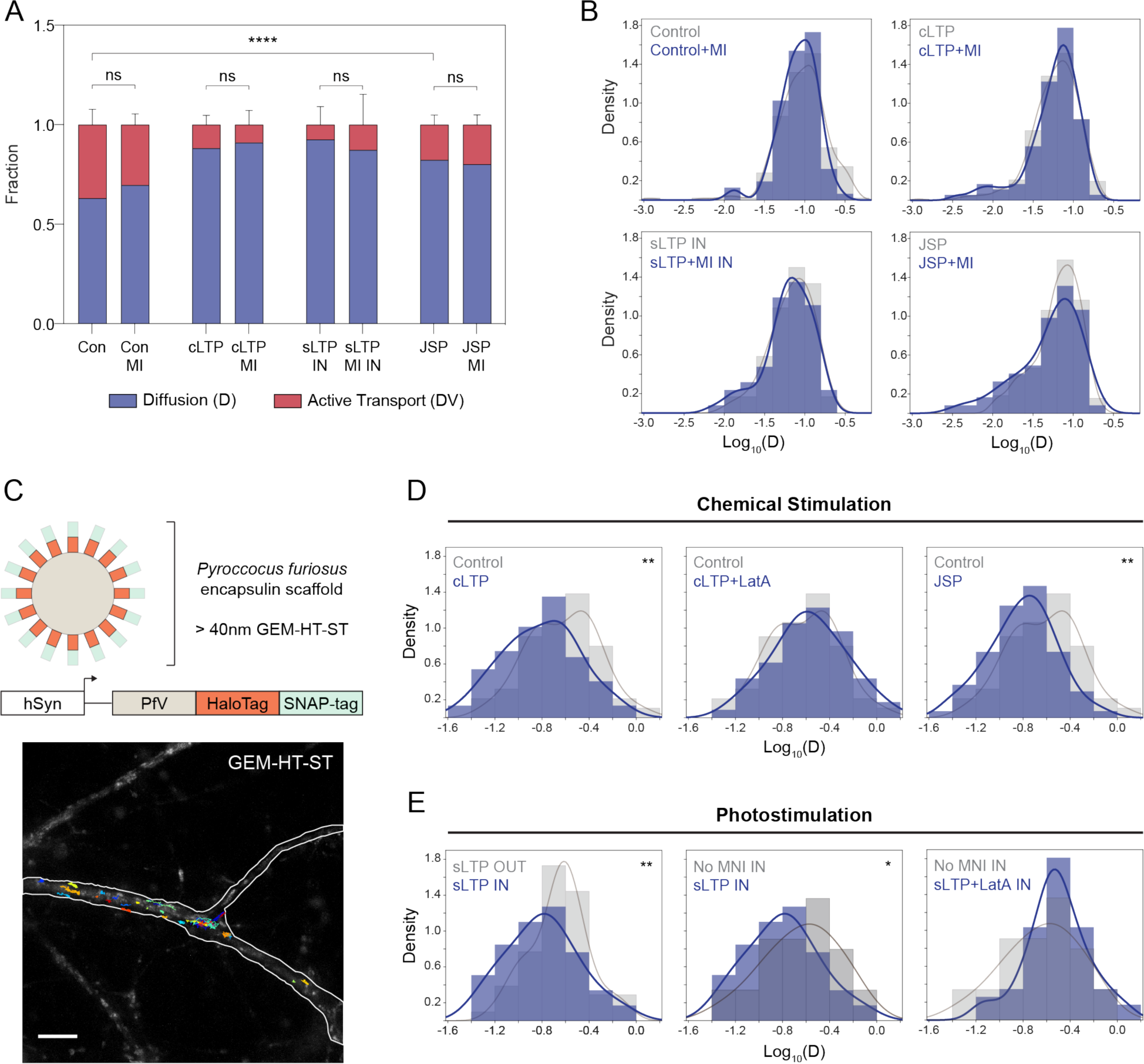
Reduced motion of GluA1-HT vesicles in the dendritic shaft during neuronal activity is mediated by actin-induced molecular crowding and is independent of myosin V and VI activity. (A) Bar graph of the fractions of GluA1-HT vesicles exhibiting active transport (DV) or diffusion (D) in dendritic shafts without stimulation (Con), during cLTP (cLTP), after sLTP (sLTP IN, in regions where actin polymerization occurred), or during treatment with Jasplakinolide (JSP) in the absence or presence of a myosin V and VI inhibitor cocktail (MI). Con vs JSP, ****p<0.0001. Significance was determined by Mann-Whitney test. n=6-14 experiments for each condition. Bars represent mean and standard deviation. Figure 5**-figure supplement 1A** for Con vs JSP side by side. (B) Distributions of diffusion coefficients of GluA1-HT vesicles in dendritic shafts without stimulation (Control), during cLTP (cLTP), after sLTP (sLTP IN), or during treatment with Jasplakinolide (JSP) in the absence or presence of a myosin V and VI inhibitor cocktail (MI). Significance was determined by Kolmogorov-Smirnov test. n=60-383 trajectories for each condition. Lines represent the probability density function of each histogram estimated by kernel density estimation (KDE). Figure 5**-figure supplement 1B** for Con vs JSP distributions. (C-E) Diffusion of a rheological probe in the dendritic shaft in response to stimulation-mediated actin polymerization. (C) Schematic: GEM-HT-ST rheological probe (top). GEM-HT-ST was created by fusing HaloTag and SNAP-tag to the PfV protein from *Pyroccocus furiosus* (bottom). When expressed, PfV-HT-ST fusion proteins self-assemble into a 40 nm encapsulin scaffold that can be labeled with either JF_549_-HTL or JF_549_-SNAP-tag ligand (JF_549_-STL). Image: trajectories of GEM-HT-ST labeled with JF_549_-HTL. Scale bar, 10 μm. (D) Distributions of diffusion coefficients of GEM-HT-ST after chemical stimulation versus no treatment. Control vs cLTP, **p=0.0047; Control vs JSP, **p=0.0016. (E) Distributions of diffusion coefficients of GEM- HT-ST after sLTP in regions where actin polymerization occurred versus controls. IN region was determined by isolating a 30 μm region flanking the uncaging site (the average length of dendrite where actin polymerization occurred near the site of uncaging during sLTP; Figure 4E). sLTP IN vs sLTP OUT, **p=0.0083; sLTP IN vs No MNI IN, *p=0.0446 Significance was determined by Kolmogorov-Smirnov test. n=44-94 trajectories for each condition. Lines represent the probability density function of each histogram estimated by kernel density estimation (KDE).

Previous studies have demonstrated that F-actin can constrain the motion of lysosomes (van Bommel et al., 2019), leading us to speculate that actin polymerization itself could block the motion of GluA1-HT vesicles. Treatment of neurons with Jasplakinolide (JSP), an F-actin stabilizer that promotes actin polymerization, is sufficient to partially inhibit active transport and reduce the rate of diffusion of GluA1-HT vesicles (**Figure 5A-B**, JSP, and **Figure 5-figure supplement 1**). Furthermore, the JSP-mediated reduction in GluA1-HT vesicle motion is not altered by myosin inhibition, indicating that F-actin itself plays a role in the altered motion of GluA1-HT vesicles (**Figure 5A-B**, JSP).

Previous studies have found that actin can induce molecular crowding in biological systems such as neuronal axons and prevent active transport of vesicles and organelles (Sood et al., 2018). We hypothesized that actin polymerization could disrupt vesicle motion by changing the rheological properties of the dendritic cytoplasm. To examine this possibility, we tested whether cLTP and sLTP would alter the diffusion of a rheological probe consisting of a 40 nm genetically encoded multimeric nanoparticle (GEM) (Delarue et al., 2018) whose size is more similar to GluA1-HT vesicles than are fluorescent proteins (**Figure 5C-E**). GEM is based on the encapsulin protein PfV of *Pyroccocus furiosus* (**Figure 5C**, schematic), which self-assembles into an icosahedral scaffold of 120 monomers when expressed. When fused to both HaloTag and the self-labeling SNAP-tag (ST), the particle arising from expression of GEM-HT-ST can be labeled with Janelia Fluor dye ligands and tracked in dendrites (**Figure 5C**, image, and **Video 6**).

We find that GEM-HT-ST diffusion is significantly reduced during cLTP induction (**Figure 5D**, cLTP), and this reduction is dependent on actin polymerization as LatA prevents the decrease in diffusion coefficient (**Figure 5D**, cLTP+LatA). Moreover, actin polymerization stimulated by JSP is sufficient to reduce the motion of GEM (**Figure 5D**, JSP). sLTP also results in a significant reduction in the rate of GEM diffusion in regions of dendritic shafts where actin polymerization occurs (**Figure 5E**, sLTP IN vs sLTP OUT) in a manner that is dependent on the presence of MNI (**Figure 5E**, No MNI In) and actin polymerization (**Figure 5E**, sLTP+LatA).

These experiments demonstrate that neuronal activity changes the rheological properties of the dendritic shaft by stimulating actin polymerization. Combined with our findings that triggering actin polymerization is sufficient to disrupt GluA1-HT vesicle motion (via JSP treatment) and that myosin inhibition did not alter activity-mediated changes in motion, these observations are consistent with the hypothesis that actin polymerization confines GluA1-HT vesicles by altering the properties of the dendric cytoplasm independent of direct interactions between GluA1 and myosin.

### Local increases in GluA1 exocytosis during neuronal activity are dependent on actin-mediated GluA1 vesicle confinement and myosin activity

Although we defined a mechanism by which actin polymerization can concentrate GluA1 vesicles near the sites of neuronal activity, it is unclear whether this mechanism contributes to increased trafficking of GluA1 to synapses during activity – whether positioning vesicles near the sites of activity also results in increased exocytosis of GluA1-HT at those sites. To study the exocytic rates of endogenous GluA1, we fused super ecliptic pHluorin (SEP) to HaloTag, and knocked this tandem reporter into the NTD of GluA1 (**Figure 6A** and **Figure 6-figure supplement 1**). SEP is a pH-sensitive variant of GFP that has low fluorescence at low pH and high fluorescence at neutral pH. We integrated HT-SEP into the extracellular NTD of GluA1 such that the tag is exposed to the low pH lumen of vesicles during transport. Intracellular GluA1 has low fluorescence until exocytosis, at which point it is exposed to neutral pH extracellular medium and exhibits strong fluorescence (**Figure 6A**, diagram). During exocytosis, GluA1-HT- SEP released from vesicles is temporarily confined at the sites of exocytosis and appears as bright puncta under fluorescence microscopy (**Figure 6A**, images, and **Video 7**). Consequently, single GluA1-HT-SEP exocytic events can be observed and quantified.

**Figure 6.**
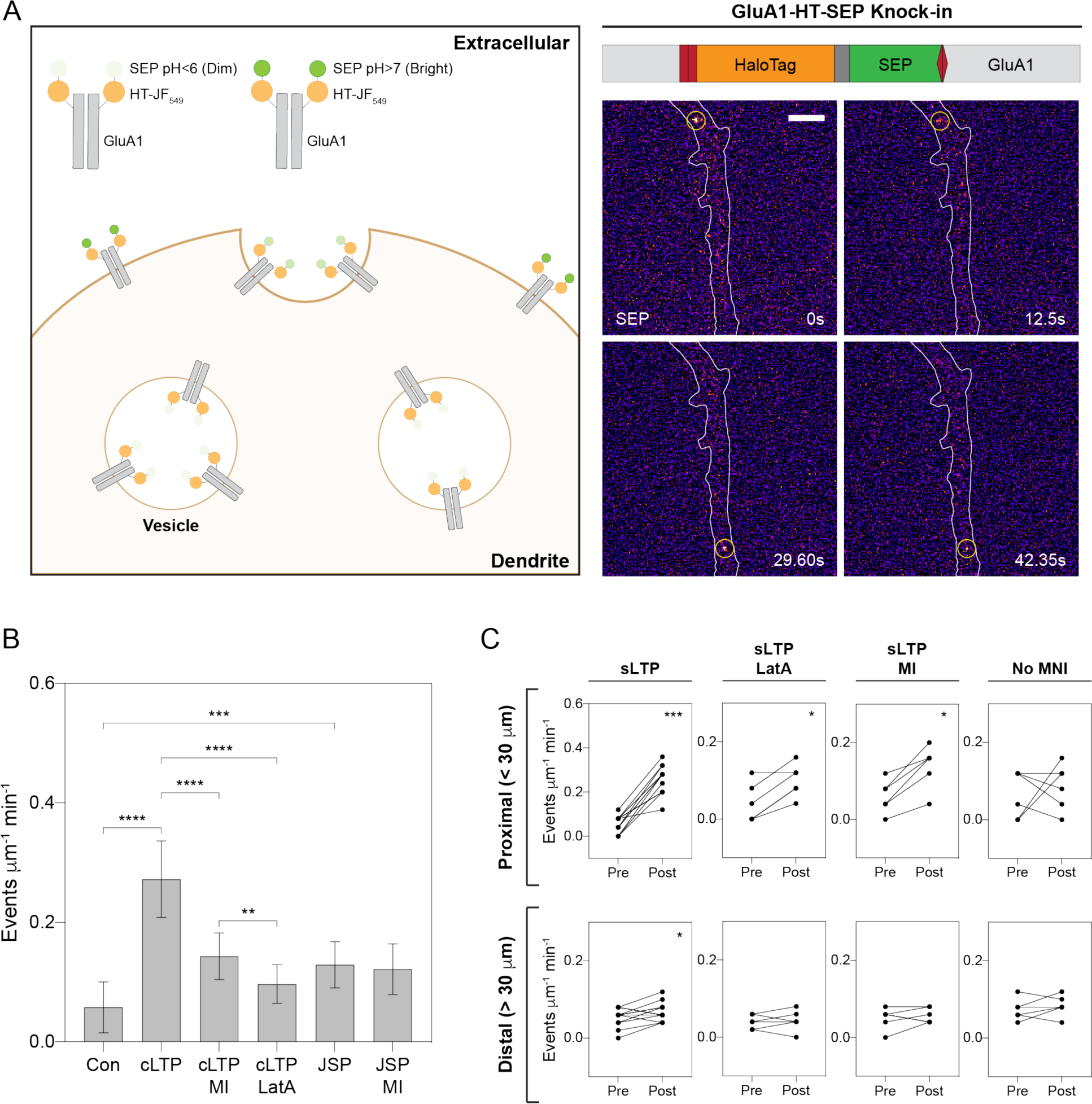
Increased GluA1-HT-SEP exocytosis during neuronal activity is dependent on actin polymerization and myosin activity. (A) Tagging endogenous GluA1 with a HaloTag-pHluorin tandem fusion reporter (HT-SEP) to track GluA1 exocytosis events. Diagram: endogenous GluA1 tagged with HT-SEP (GluA1-HT- SEP, schematic of fusion on right) has low SEP fluorescence inside vesicles due to the low pH of the vesicle lumen. When a GluA1 exocytosis event occurs, SEP will be exposed to the neutral pH of the extracellular medium, resulting in increased fluorescence. Because receptors are temporarily spatially restricted during exocytosis, a spot with fluorescence can be observed during the event. Images: time lapse of two GluA1-HT-SEP exocytosis events (yellow circles). Scale bar, 5 μm. (B) Bar graph of GluA1-HT-SEP exocytosis in response to chemical stimulation in the absence or presence of a myosin inhibitor cocktail (MI) or LatA. Bars represent mean events per μm per min and standard deviation. Con vs cLTP, ****p<0.0001; Con vs JSP, ***p=0.0005; cLTP vs cLTP+MI, ****p<0.0001, cLTP vs cLTP+LatA, ****p<0.0001; cLTP+MI vs cLTP+LatA, **p=0.0063. Significance was determined by Mann-Whitney test. n=9-12 experiments for each condition. (C) Line graphs of GluA1-HT-SEP exocytic events before and after sLTP (sLTP), sLTP in the presence of LatA (sLTP+LatA), sLTP in the presence of myosin inhibitors (sLTP+MI), and sLTP stimulation in the absence of MNI (No MNI). Proximal: GluA1-HT-SEP exocytic events within the 30 μm region surrounding the site of uncaging (30 μm was the average length of dendrite where actin polymerization occurred near the site of uncaging during sLTP; see Figure 4E). sLTP, ***p=0.0005; sLTP+LatA, *p=0.0312; sLTP+MI, *p=0.0312. Distal: GluA1-HT-SEP exocytic events outside of the 30 μm region surrounding the site of uncaging. sLTP, *p=0.0430. Significance was determined by Wilcoxon matched paired test. n=6-12 experiments for each condition.

Using the GluA1-HT-SEP reporter, we sought to confirm previous findings that cLTP increases GluA1 exocytosis (Kopec et al., 2006), and then establish whether stimulation-mediated actin polymerization plays a role in increased exocytosis. GluA1-HT-SEP exocytic events occur at a low rate prior to stimulation, but increase dramatically during cLTP induction (**Figure 6B**). The increase in GluA1-HT-SEP exocytosis is dependent on actin polymerization, as LatA reduces the rate of exocytosis during cLTP induction. Myosin inhibition also disrupts cLTP-stimulated GluA1-HT-SEP exocytosis (**Figure 6B**), suggesting that while myosin does not play a role in disrupting transport of GluA1-HT vesicles, it plays a role in regulating exocytosis of GluA1-HT-SEP. Interestingly, neither LatA treatment nor myosin inhibition completely blocks the cLTP-mediated increase in GluA1-HT-SEP exocytosis (**Figure 6B**), indicating that not all exocytic events require F-actin or myosin (e.g., already docked GluA1-HT-SEP vesicles). Surprisingly, actin polymerization stimulated by JSP also significantly increases GluA1-HT-SEP exocytosis, even in the presence of myosin inhibitors (**Figure 6B**), demonstrating that F-actin is itself sufficient to promote exocytosis. However, it is unclear whether F-actin exerts this effect on GluA1-HT-SEP exocytosis simply by concentrating GluA1-HT vesicles or by directly interacting with vesicles to increase exocytosis. Overall, these observations demonstrate that actin polymerization mediates GluA1-HT-SEP exocytosis during cLTP.

We next tested whether sLTP results in local increases in GluA1-HT-SEP exocytosis, and if the increase in exocytosis is spatially correlated with and dependent on actin polymerization (**Figure 6C**). In the absence of a marker for F-actin (due to overlapping fluorescence signals between tractin and SEP), we defined the area proximal to the site of stimulation – the 30 µm region surrounding the uncaging site – as the region of actin polymerization based on our previous observations (**Figure 4D-E**). sLTP stimulation increases GluA1-HT-SEP exocytosis events both proximal and distal to the site of uncaging (**Figure 6C**, sLTP). However, the number of events is much greater proximal to the uncaging site. Similar to their effect on cLTP-mediated GluA1-HT-SEP exocytosis, LatA treatment and myosin inhibition both partially block sLTP- mediated increases in GluA1-HT-SEP exocytosis (**Figure 6C**, sLTP+LatA and sLTP+MI).

When MNI is removed from the media, there is no increase in exocytic events after sLTP (**Figure 6C**, No MNI).

Based on our observations, we conclude that the accumulation of GluA1-HT vesicles near sites of sLTP is spatially correlated with increased exocytosis of GluA1-HT-SEP, and that local disruption of GluA1-HT vesicle motion and the increase in GluA1-HT-SEP exocytosis are both dependent on actin polymerization in the dendritic shaft. However, this correlation does not demonstrate that concentration of GluA1-HT-SEP vesicles near the site of neuronal activity directly contributes to increased trafficking of GluA1-HT-SEP to the cell surface. In other words, it is unclear whether GluA1-HT-SEP vesicles destined for exocytosis (i.e., pre-exocytosis vesicles) are drawn from a local source or from distal loci via long-range active transport immediately prior to exocytosis. To examine these two possibilities, we sparsely labeled GluA1- HT-SEP vesicles with JF dye ligands and simultaneously tracked vesicle motion and exocytosis (**Figure 7A** and **Video 8**).

**Figure 7.**
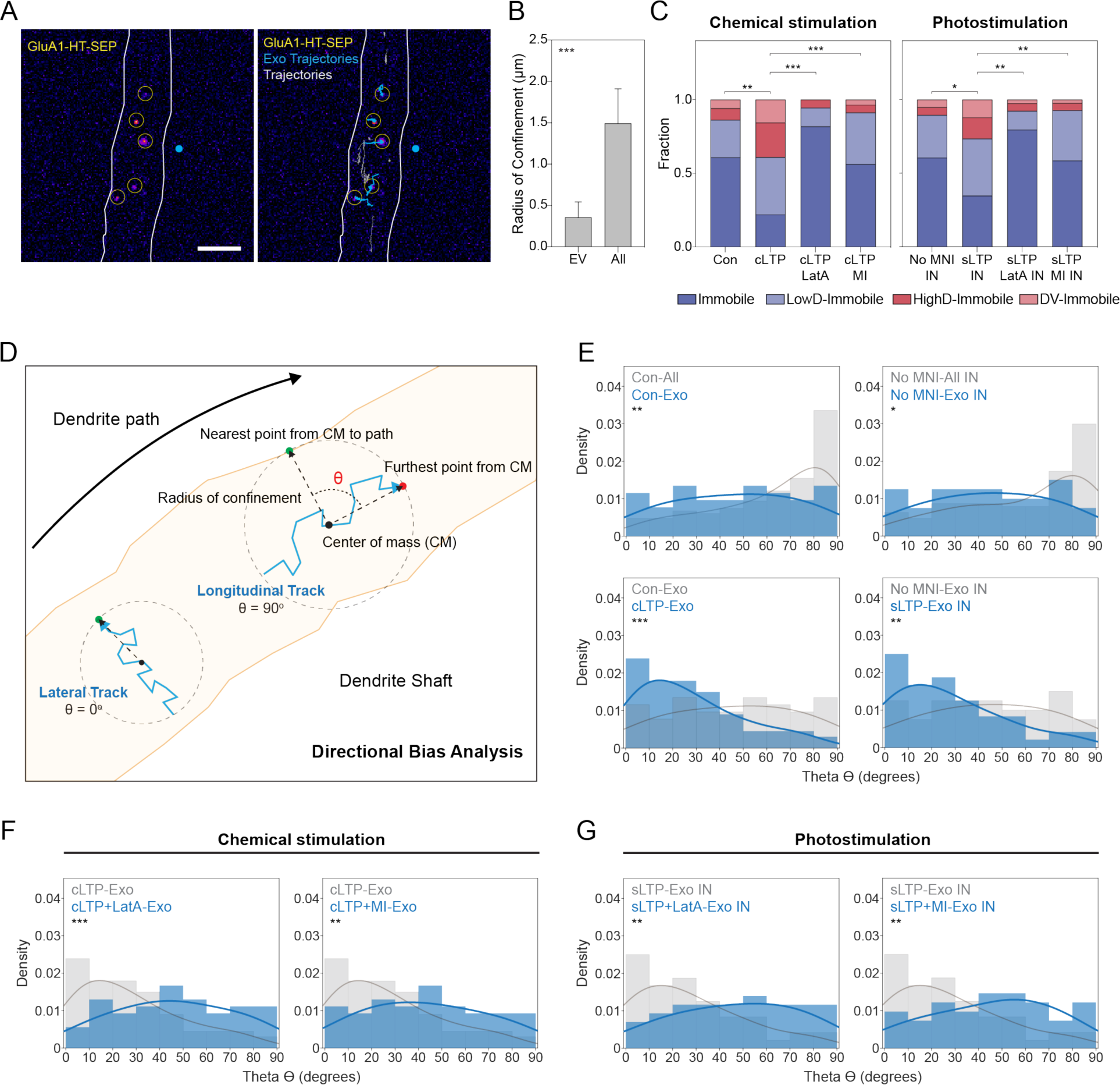
GluA1-HT-SEP vesicles are drawn from a local pool in the dendritic shaft prior to exocytosis and exhibit lateral, not longitudinal, motion that is dependent on actin polymerization in the shaft during neuronal activity. (A) Left: representative projection of GluA1-HT-SEP exocytosis over time after sLTP (i.e., the brightest pixels from each image in a timelapse compressed into a single image). Yellow circles indicate exocytic events. Blue dot indicates site of uncaging. Right: GluA1-HT-SEP vesicle trajectories overlaid on the projection of exocytosis over time. Blue trajectories are GluA1-HT- SEP vesicles that dock and undergo exocytosis. Gray trajectories are GluA1-HT-SEP vesicles with no detected exocytosis during the span of imaging. Scale bar, 5 μm. (B) Mean radius of confinement for GluA1-HT-SEP vesicle trajectories prior to exocytosis (Exo) and for all GluA1-HT-SEP vesicle trajectories (All). Error bars represent 95% confidence intervals. ***p=0.0001 by Kolmogorov-Smirnov test. n=52-149 trajectories. (C) Bar graphs of GluA1-HT-SEP motion types (Figure 7**-figure supplement 1**) prior to exocytosis under either chemical stimulation or photostimulation. Motion states were determined by HMM-Bayes analysis. For photostimulation experiments, only trajectories in regions where actin polymerization occurred were used (see Figure 4E). Con vs cLTP, **p=0.0094; cLTP vs cLTP+LatA, ***p=0.0009; cLTP vs cLTP+MI, ***p=0.0002; No MNI IN vs sLTP IN, *p=0.0267; sLTP IN vs sLTP+LatA IN, **p=0.0082; sLTP IN vs sLTP+MI IN, **p=0.0010. Significance was determined by Mann-Whitney test. n=9-15 experiments for each condition. (D and E) Directional biases of GluA1-HT-SEP vesicles in the dendritic shaft. (D) Diagram of directional bias analysis. The directional bias of a trajectory can be determined by calculating the angle, theta (0), created between the line from the center of mass of a trajectory (CM) to the nearest point on the dendrite path and the line from the CM to the furthest point from the CM. Theta of 90° indicates longitudinal movement while theta of 0° indicates lateral movement. (E) Top left: distributions of theta for pre-exocytosis vesicles under unstimulated conditions (Con- Exo) versus all vesicles under unstimulated conditions (Con-All). **p=0.0043. Bottom left: distributions of theta for pre-exocytosis vesicles during cLTP (cLTP-Exo) versus unstimulated conditions. ***p=0.0009. Top right: distributions of theta for pre-exocytosis vesicles after sLTP stimulation in the absence of MNI (No MNI-Exo IN) versus all vesicles after sLTP stimulation in the absence of MNI (No MNI-All IN). *p=0.0175. Bottom right: distributions of theta for pre- exocytosis vesicles during sLTP (sLTP-Exo IN) versus vesicles during sLTP stimulation in the absence of MNI. **p=0.0084. Significance was determined by Kolmogorov-Smirnov test. n=40- 149 trajectories for each condition. Lines represent the probability density function of each histogram estimated by kernel density estimation (KDE). (F and G) Directional biases of pre-exocytosis GluA1-HT-SEP vesicles in the dendritic shaft during neuronal activity in the absence or presence of LatA or a myosin inhibitor cocktail (MI). (F) Left: distributions of theta for pre-exocytosis GluA1-HT-SEP vesicles during cLTP (cLTP- Exo) versus pre-exocytosis GluA1-HT-SEP vesicles during cLTP in the presence of LatA (cLTP+LatA-Exo). ***p=0.0010. Right: distributions of theta for pre-exocytosis GluA1-HT-SEP vesicles during cLTP versus pre-exocytosis vesicles during cLTP in the presence of myosin inhibitors (cLTP+MI-Exo). **p=0.0043. (G) Left: distributions of theta for pre-exocytosis vesicles after sLTP (sLTP-Exo IN) versus pre-exocytosis vesicles after sLTP in presence of LatA (sLTP+LatA-Exo IN). **p=0.0053. Right: distributions of theta for pre-exocytosis vesicles during sLTP versus pre-exocytosis vesicles during sLTP in the presence of myosin inhibitors (sLTP+MI-Exo IN). **p=0.0033. Significance was determined by Kolmogorov-Smirnov test. n=41-54 trajectories for each condition. Lines represent the probability density function of each histogram estimated by kernel density estimation (KDE).

We hypothesized that if GluA1-HT-SEP vesicles are drawn from local sources then the trajectories of GluA1-HT-SEP vesicles prior to exocytosis should travel relatively short net distances. Consequently, we measured the radius of confinement for the trajectories of pre- exocytosis GluA1-HT-SEP vesicles (**Figure 7B**). The radius of confinement estimates the area in which a trajectory is confined by estimating the size of the circle that best encompasses the trajectory. When compared to all GluA1-HT-SEP vesicle trajectories (including vesicles that did not exocytose during the duration of a time-lapse sequence), pre-exocytosis vesicles have significantly smaller radii of confinement (**Figure 7B**), demonstrating that GluA1-HT-SEP vesicles are not imported from distal loci immediately prior to exocytosis.

We then reasoned that if pre-exocytosis GluA1-HT-SEP vesicles travel relatively short distances, few would require active transport. We used HMM-Bayes analysis to infer the motion states of pre-exocytosis GluA1-HT-SEP vesicles and find that they often exhibit at least two motion states: an initial state of active transport or diffusion followed by a second state of immobility (**Figure 7-figure supplement 1**). Because the immobile state always precedes exocytosis, we believe this state is representative of vesicle docking. We separated pre- exocytosis motion into four categories: 1) immobility (i.e., docked prior to imaging); 2) low rates of diffusion prior to immobility; 3) high rates of diffusion prior to immobility; and 4) active transport prior to immobility (**Figure 7C** and **Figure 7-figure supplement 1**). We separated vesicles with diffusive motion into two categories because these trajectories were qualitatively different. For example, trajectories with high diffusion coefficients often exhibited a linear path with a directional bias, similar to vesicles exhibiting active transport (**Figure 7-figure supplement 1C-D**).

Given that pre-exocytosis GluA1-HT-SEP vesicles had relatively small search spaces, we anticipated most vesicles would exhibit immobility or low rates of diffusion prior to exocytosis. Instead, we find that the fraction of pre-exocytosis GluA1-HT-SEP vesicles exhibiting active transport and high rates of diffusion increases significantly in response to both cLTP and sLTP (**Figure 7C**). cLTP- and sLTP-induced active transport and high rates of diffusion prior to exocytosis are dependent on both actin polymerization and myosin activity, as inhibiting either results in near complete losses of these vesicle populations. Inhibition of actin polymerization, but not myosin, also reduces the fraction of pre-exocytosis GluA1-HT-SEP vesicles with low rates of diffusion, suggesting that F-actin may also promote the exocytosis of GluA1-HT-SEP by confining GluA1-HT-SEP vesicles (**Figure 7C**). Most pre-exocytosis GluA1-HT-SEP vesicles under LatA treatment or myosin inhibition are immobile (i.e., already docked). Overall, these findings suggest that stimulation specifically increases the transport and motion of *pre-exocytosis* GluA1-HT-SEP vesicles in a manner dependent on actin polymerization and myosin activity, even while stimulation generally inhibits the motion of *all* GluA1-HT-SEP vesicles.

Although surprising, the observation that pre-exocytosis GluA1-HT-SEP vesicles exhibit limited radii of confinement does not preclude the possibility that vesicles with short-range motion proximal to the site of neuronal activity could also be driven by active transport or high diffusion rates. Although we find that long-range motion along the length of the dendritic shaft is reduced during neuronal activity, it is possible that transport of GluA1-HT-SEP vesicles perpendicular to the dendritic shaft towards the cell membrane might increase in a manner dependent on myosin and F-actin as a mechanism to promote exocytosis. To test this possibility, we analyzed the directional bias, described by the angle theta, of each pre-exocytosis GluA1-HT- SEP vesicle trajectory (**Figure 7D** and **Methods** for details). Theta of ∼90° indicates that the vesicle is moving parallel to the dendritic shaft (i.e., longitudinal motion) whereas theta of ∼0° indicates the vesicle is moving perpendicular to the dendritic shaft (i.e., lateral motion).

When we examine theta for all GluA1-HT-SEP trajectories (regardless of whether they undergo exocytosis), we find strong biases for longitudinal motion under unstimulated conditions and sLTP in the absence of MNI (**Figure 7E**, top panels, Con-All and No MNI-All IN), consistent with our observation that a large fraction of GluA1-HT vesicles undergo long-range active transport along the path of the dendritic shaft under unstimulated conditions (**Figure 2E- F**). By contrast, pre-exocytosis GluA1-HT-SEP vesicles did not exhibit a strong bias for longitudinal or lateral motion under unstimulated conditions, consistent with the observation that long-range longitudinal motion is lost prior to exocytosis (**Figure 7E**, top panels, Con-Exo and No MNI-Exo IN). When stimulated with cLTP or sLTP, pre-exocytosis GluA1-HT-SEP vesicles exhibit a bias for lateral motion (**Figure 7E**, bottom panels, cLTP-Exo and sLTP-Exo IN). In addition, pre-exocytosis vesicles with active transport or high rates of diffusion also exhibit a bias for lateral motion (**Figure 7-figure supplement 2**). When combined, these results suggest that stimulation induces the lateral transport of GluA1-HT-SEP vesicles to sites of exocytosis.

Importantly, the change in directional bias in pre-exocytosis GluA1-HT-SEP vesicles during activity is dependent on actin polymerization and myosin activity as either LatA treatment or myosin inhibition prevented the increase in lateral motion during cLTP and sLTP (**Figure 7F- G**).

In summary, pre-exocytosis GluA1-HT-SEP vesicles under unstimulated conditions exhibit short-range diffusive motion that does not have a strong bias for either longitudinal or lateral motion. Neuronal activity increases the fraction of pre-exocytosis GluA1-HT-SEP vesicles exhibiting either high rates of diffusion or active transport, but lateral to the path of the dendrite. Importantly, both increased transport and the change in directional bias are dependent on actin polymerization and myosin activity. When coupled with the finding that actin polymerization and myosin activity are required for activity-dependent increases in GluA1-HT- SEP exocytosis, these results strongly indicate that F-actin networks in the dendritic shaft are a substrate for myosin-dependent trafficking of GluA1-HT-SEP vesicles to the dendrite periphery.

## Discussion

In this study, we have developed a novel method to identify, track and characterize the motion of vesicles containing GluA1, enabling us to better understand how AMPARs are delivered specifically to sites undergoing neuronal activity. We use homology-independent targeted integration (HITI) to tag endogenous GluA1 with HaloTag (GluA1-HT) and then a block-and-chase labeling protocol with Janelia Fluor (JF) dye ligands to achieve a sparse labeling density suitable for the detection of GluA1-HT vesicles. After identifying GluA1-HT vesicles based on their photobleaching characteristics, we utilize single-particle tracking (SPT) followed by hidden Markov modeling with Bayesian model selection (HMM-Bayes) to describe their motion. Using this technique, we find that GluA1-HT vesicles experience a loss of long-range motion along the path of the dendrite (i.e., longitudinal motion) during chemical and structural LTP (cLTP and sLTP). GluA1-HT vesicles become locally confined by actin polymerization that is triggered in the dendritic shaft near the site of sLTP stimulation, resulting in an increase in the intracellular reservoir of GluA1-HT near that site. We then utilized a pHluorin-HaloTag fusion inserted into GluA1 (GluA1-HT-SEP) to examine how local vesicular reservoirs of GluA1 contribute to GluA1 exocytosis. We find that a large fraction of pre- exocytosis GluA1-HT-SEP vesicles undergo short-range transport perpendicular to the length of the dendritic shaft (i.e., lateral motion) near the site of neuronal stimulation in a manner dependent on both actin polymerization and myosin activity.

Based on these findings, we propose a new model in which neurons utilize actin polymerization in the dendritic shaft to specify the location to which AMPARs are delivered during neuronal activity (**Figure 8**). First, actin polymerization occurs in the dendritic shaft proximal to the site of neuronal activity, resulting in molecular crowding that changes the rheological properties of the dendritic cytoplasm at this location (**Figure 8A-B**). The increased crowding inhibits the longitudinal motion of GluA1 vesicles, concentrating intracellular GluA1 near the site of neuronal activity (**Figure 8A-B**). F-actin then acts as a substrate for myosin V and/or VI – activated by the influx of calcium during neuronal activity (Correia et al., 2008; Wang et al., 2008) – to translocate into the dendritic shaft and recruit GluA1 vesicles to the periphery of the dendrite (**Figure 8C**). At the periphery, GluA1 vesicles undergo exocytosis, increasing the amount of surface bound GluA1 that can then diffuse into synapses (**Figure 8D**). Critically, actin polymerization by itself can disrupt the longitudinal motion of vesicles to create a larger reservoir of intracellular GluA1, but both F-actin and myosin are required for short-range transport of GluA1 to the periphery of the cell.

**Figure 8.**
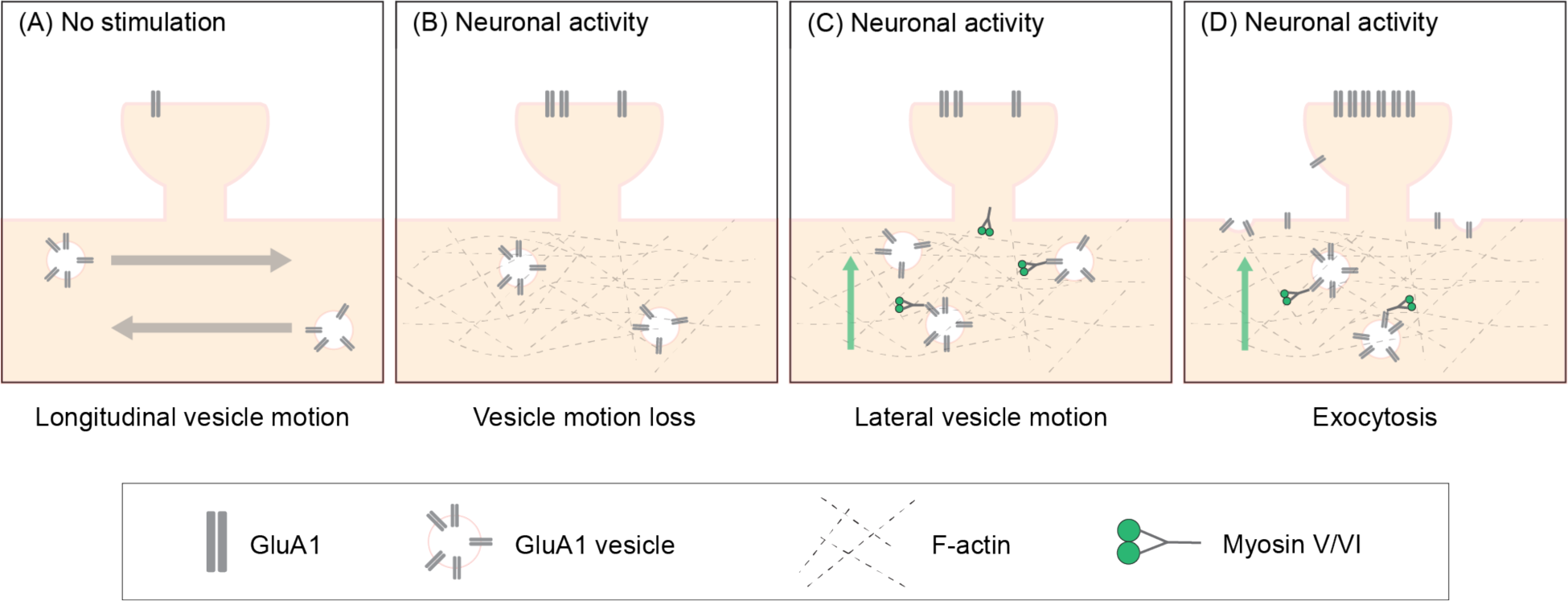
Actin polymerization in the dendritic shaft proximal to the site of neuronal activity promotes GluA1 exocytosis by increasing the local pool of GluA1 vesicles and facilitating myosin-dependent transport to the dendrite periphery. (A) Under unstimulated conditions, GluA1 vesicles are transported longitudinally along the dendritic shaft. (B) During the induction of neuronal activity, actin polymerizes in the dendritic shaft near the site of activity. Actin polymerization disrupts the longitudinal motion of vesicles resulting in an increased pool of GluA1 vesicles near the site of activity. (C) Myosin V and/or VI, which is inactive and sequestered in dendritic spines in unstimulated conditions, is activated by calcium influx during neuronal activity. Myosin subsequently translocates to the base of the dendritic spines. (D) Myosin V and/or VI recruits GluA1 vesicles to the periphery resulting in increased exocytosis of GluA1.

Our observations of GluA1 vesicle trafficking differ somewhat from previous reports (Esteves da Silva et al., 2015; Bowen et al., 2017; Hangen et al., 2018). Hangen et al. (2018) found that GluA1 vesicles undergoing active transport during cLTP induction had decreased velocity and paused (i.e., vesicles temporarily lost active transport) more frequently. By contrast, we find that only a small fraction of GluA1-HT vesicles undergo active transport during cLTP induction, and vesicles that exhibit active transport often switch to diffusion and rarely switch back to active transport (i.e., vesicles stably lose active transport). Our observations lead us to conclude that GluA1-HT vesicles are locally confined during cLTP induction rather than paused.

Differences in methodological approaches likely contribute to discrepancies between our observations and those of Hangen et al. (2018). Hangen et al. (2018) used chemically induced dimerization (Inoue et al., 2005) to control the release of exogenous GluA1 from the ER and tracked GluA1 vesicles as they crossed photobleached regions of dendrite, enabling them to unambiguously identify GluA1 vesicles and characterize active transport using kymograph analysis. However, since photobleaching removes the signal from all vesicles (both those undergoing active transport and diffusion) within the bleached region of the dendrite, only vesicles originating from outside of the bleached region being transported through the bleached region can be characterized. As we did not need to bleach obstructive signal prior to imaging, we find that many GluA1-HT vesicles exhibit diffusion, particularly after stimulation, in addition to active transport.

Our approaches to photostimulation may have also contributed to differences in observations. Hangen et al. (2018) simultaneously uncaged glutamate at 10 adjacent spines with a series of 10 laser pulses and found that the transient influx of calcium *during* stimulation was correlated with a decrease in GluA1 vesicle velocity and increased vesicle pausing. By contrast, we apply glutamate-uncaging laser pulses at 2 Hz for 50 seconds to a single spine (i.e., sLTP), and find that GluA1-HT vesicles are confined near the site of uncaging *after* stimulation. In contrast to pausing, GluA1-HT vesicle confinement is relatively stable, leading us to hypothesize that a lasting change to the dendritic cytoplasm, rather than transient calcium influx, results in vesicle confinement. We identified actin polymerization near spines undergoing sLTP stimulation as the mechanism mediating GluA1-HT vesicle confinement. Importantly, stimulating actin polymerization with Jasplakinolide (JSP), which ostensibly does not increase intracellular calcium, is sufficient to confine GluA1-HT vesicles. Nevertheless, our findings do not eliminate the possibility that calcium is involved in confinement. Calcium signaling plays a significant role in the reorganization of actin networks, especially in dendritic spines after photostimulation (Bosch et al., 2015). Thus, while F-actin mediates the changes in motion exhibited by GluA1 vesicles during neuronal activity, actin polymerization in the dendritic shaft may be triggered by calcium signaling (e.g., through CaMKII activation).

The site from which actin polymerization in the dendritic shaft originates is unclear. Previous studies have identified extensive actin remodeling at the bases of dendritic spines during neuronal activity (Schatzle et al., 2018), leading us to speculate that the expansion of F-actin in dendritic spines during stimulation extends into the dendritic shaft. This hypothesis is attractive also because F-actin would physically connect the vesicular reservoir of GluA1 to the site of stimulation. Others have identified F-actin networks at shaft synapses (synapses located on the dendritic shaft rather than on dendritic spines; van Bommel et al., 2019), as well as the remodeling of periodic submembrane actin rings into F-actin fibers in the dendritic shaft (Lavoie-Cardinal et al., 2020), raising the possibility that actin polymerization originates from a location in the dendritic shaft proximal to the site of stimulation. Further investigation could reveal interactions between post-stimulation signaling and actin polymerization.

Interestingly, actin polymerization in the dendritic shaft is sufficient to disrupt the longitudinal transport of GluA1-HT independent of myosin activity. The loss of longitudinal transport is the result of F-actin-induced molecular crowding, which creates obstructions to transport near the site of stimulation, preventing long-range motion and decreasing the rate of diffusion. Critically, the effect of F-actin-induced molecular crowding is likely not specific to GluA1 vesicles, but rather affects particle motion based on the size of the particle. This raises the possibility that actin polymerization in the dendritic shaft is a general mechanism to coordinate the localization of important proteins and organelles at the site of neuronal activity. For example, F-actin patches in the dendritic shaft position lysosomes in shaft synapses via passive trapping and myosin-mediated anchoring to support AMPAR turnover (van Bommel et al., 2019; Goo et al., 2017). F-actin may play a similar role in positioning lysosomes, and other organelles, near the sites of stimulation.

We find that F-actin in the dendritic shaft also promotes GluA1-HT-SEP exocytosis by acting as a substrate for short-range transport to the dendrite periphery. Myosin V and/or VI are required for the lateral transport of pre-exocytosis GluA1-HT-SEP vesicles. Given this role, we hypothesize that myosin-mediated transport may also be a mechanism to regulate which cargoes are ultimately exocytosed or brought to the spines undergoing LTP because myosin will interact with select cargoes (Hammer and Sellers, 2011) whereas F-actin has a general effect on motion. Other mechanisms may also add specificity to which cargoes are destined for exocytosis. For example, myosin cargo adaptors that interact with GluA1 may further enhance the specificity of transport (Correia et al., 2008).

Because AMPARs are primarily transported along the length of dendrites on microtubules (Esteves da Silva et al., 2015; Setou et al., 2002; Hoerndli et al., 2013), our finding that F-actin mediates the lateral transport of pre-exocytosis GluA1-HT-SEP vesicles suggests that vesicles must be transferred from microtubule-based transport to actin-based transport prior to exocytosis. The precise mechanism for this exchange is not known but our finding that F-actin disrupts long-range active transport along the length of the dendrite independent of myosin – creating a larger pool of diffusing vesicles – suggests that the transfer is not a continuous hand- off between kinesin/dynein and myosin. Rather, we hypothesize that actin polymerization changes the rheological properties in the cytoplasm in a manner that is generally obstructive to motion, resulting in the stalling of vesicular cargo or detachment from microtubule tracks (Nettesheim et al., 2020). However, because F-actin is a substrate for myosin, actin polymerization selectively enhances myosin-based transport in this environment. In other words, neuronal activity alters the dendritic cytoplasm such that it favors myosin-mediated transport over kinesin/dynein-mediated transport. Such a mechanism could have important implications for how cargo in general is transferred as changes to the subcellular environment could alter the mode of transport indirectly rather than through the direct interactions between motors, cargo, and cytoskeletal elements.

F-actin and myosin also play direct roles in facilitating exocytosis by enhancing vesicle anchoring and fusion (Rudolf et al., 2010; Meunier and Gutierrez, 2016). Consequently, it is difficult to determine whether F-actin and myosin contribute more to the surface expression of GluA1 by affecting trafficking or exocytosis. We find that many GluA1-HT-SEP vesicles undergo myosin-dependent lateral transport immediately prior to exocytosis, suggesting that myosin-dependent transport increases GluA1 in the cell membrane. Strikingly, some pre- exocytosis GluA1-HT-SEP vesicles exhibit slow rates of diffusion prior to docking and exocytosis. Critically, these vesicles increase in number in response to stimulation in a manner dependent on actin polymerization but not myosin activity. This observation raises the possibility that the confinement of GluA1 vesicles by F-actin, in the absence of transport, is sufficient to induce increased GluA1 docking and exocytosis, though the precise mechanism is unclear. One possibility may be that confinement in response to actin polymerization simply situates GluA1 vesicles near sites of exocytosis, and consequently only very limited diffusion is necessary before a vesicle is docked. Alternatively, the polymerization of actin may also directly influence GluA1 vesicles towards docking. Previous studies have demonstrated that rearrangements in subcortical F-actin networks can promote vesicle fusion by bringing vesicles closer to the cell membrane (Papadopulos et al., 2015). Such a mechanism raises interesting questions about how F-actin interacts with objects to influence motion.

The precise control of synaptic protein trafficking is vital to synaptic transmission, and thus learning and memory. Nevertheless, many mechanisms regulating synaptic protein trafficking remain to be fully understood. Here we identify a novel mechanism through which actin polymerization in the dendritic shaft can regulate the surface expression of GluA1 specifically at sites with stimulating inputs. Because F-actin can exert direct and indirect effects on a variety of particles in the dendritic cytoplasm, our findings raise interesting questions regarding whether actin polymerization is a general mechanism to coordinate the delivery of proteins during synaptic plasticity. Elucidating whether and how actin regulates the motion of proteins in the dendritic shaft during neuronal activity could help us better understand the cellular bases for learning and memory.

## Materials and methods

Animal work was conducted according to the Institutional Animal Care and Use Committee guidelines of Janelia Research Campus.

### Key resources table

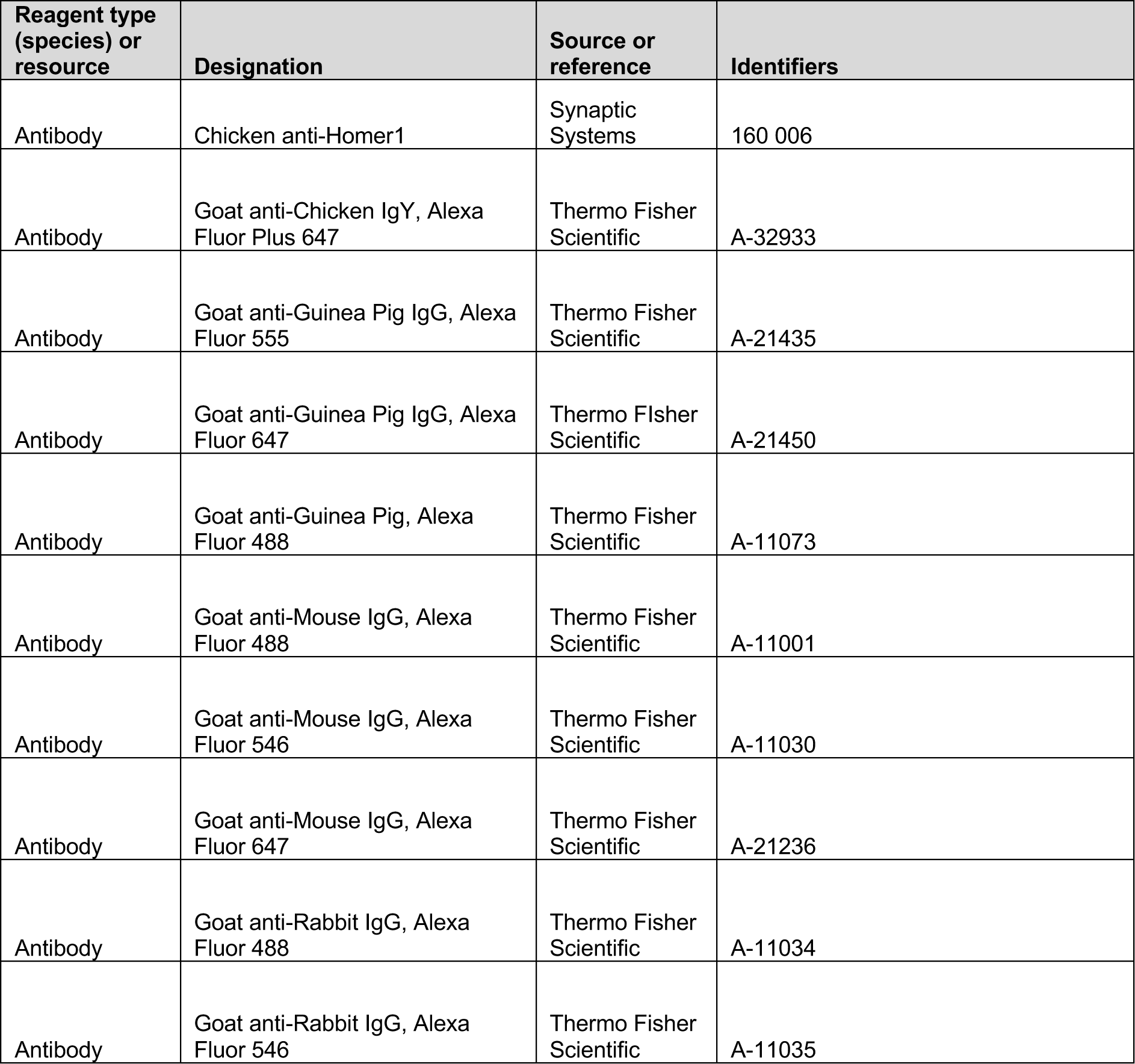
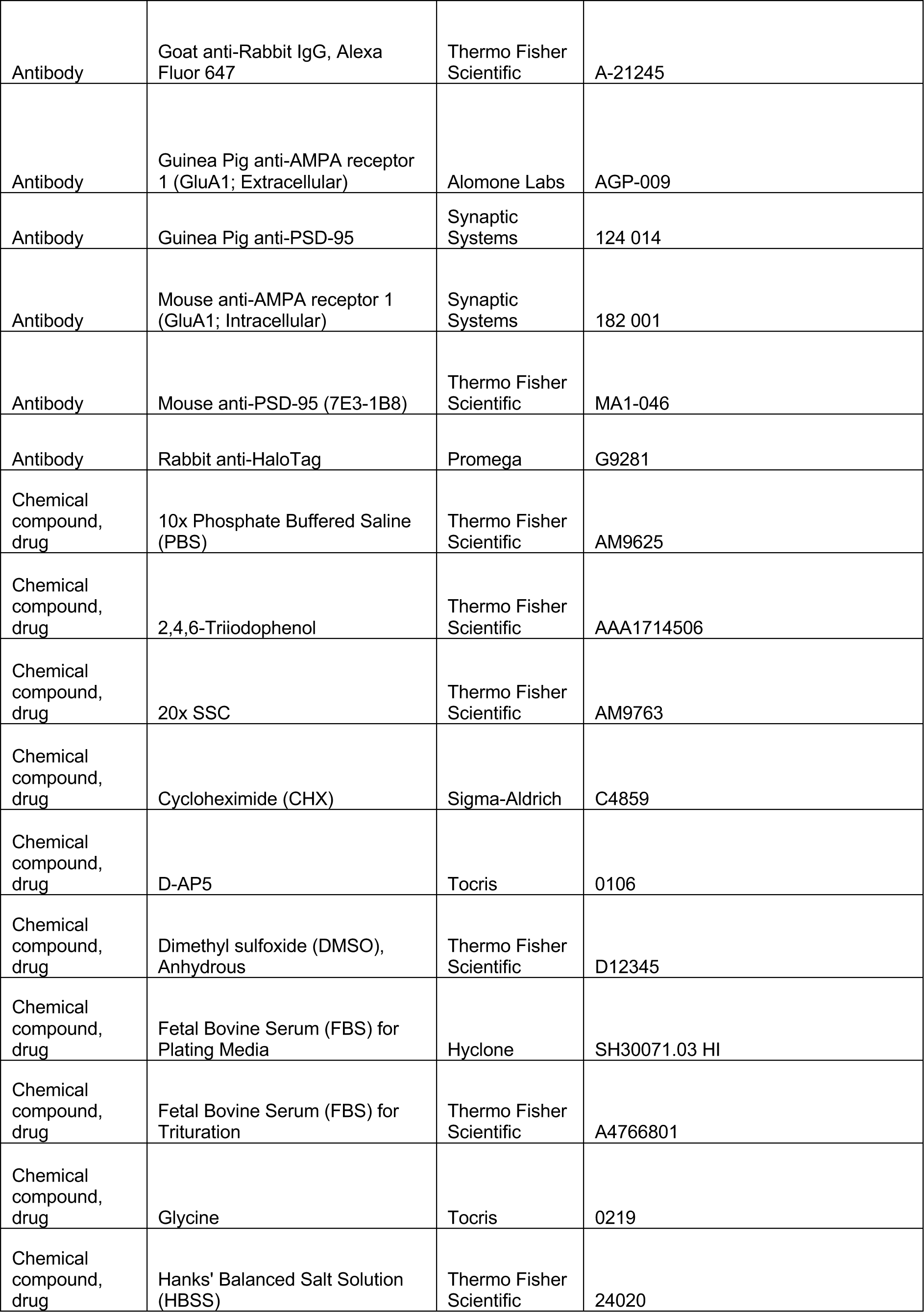
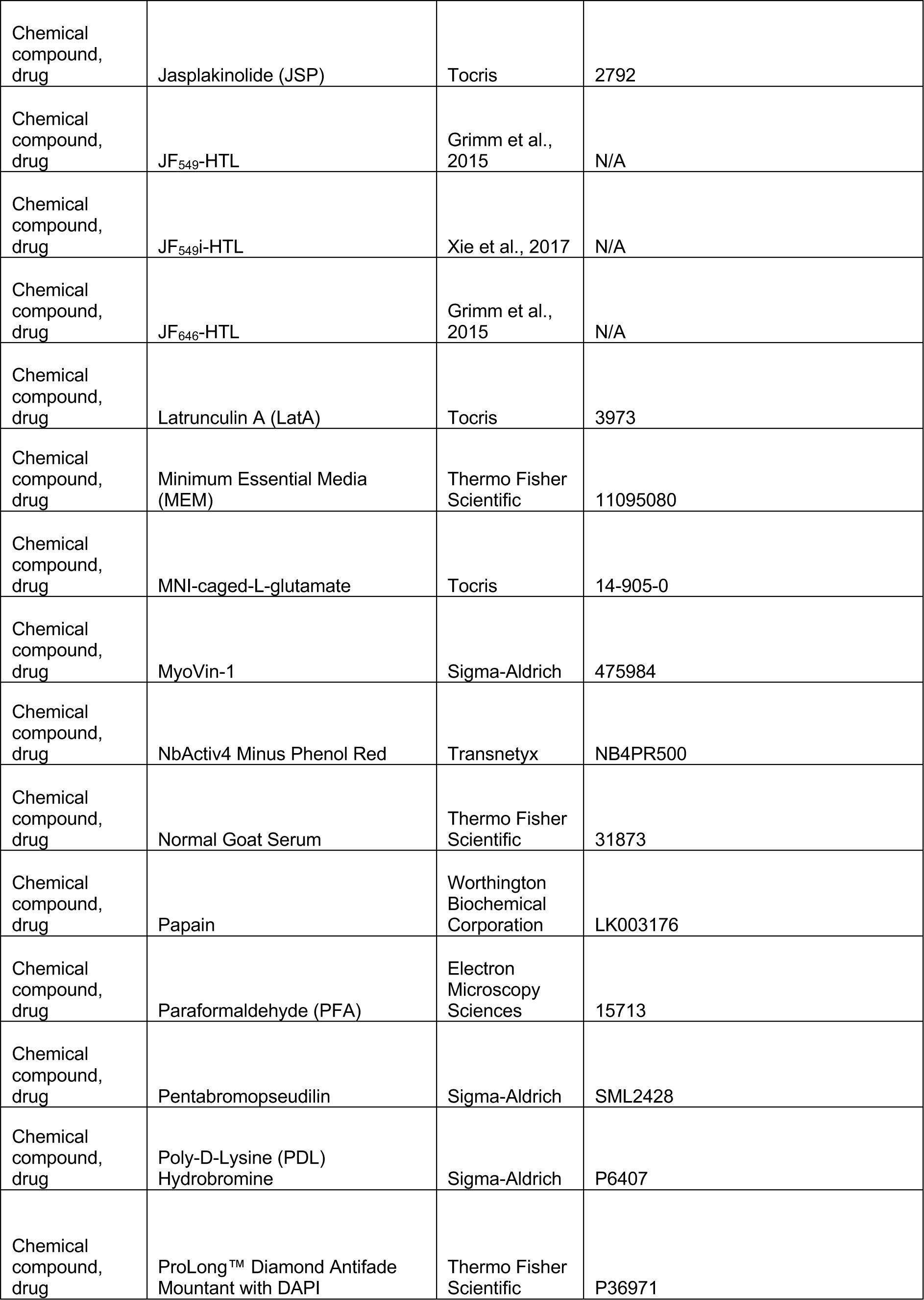
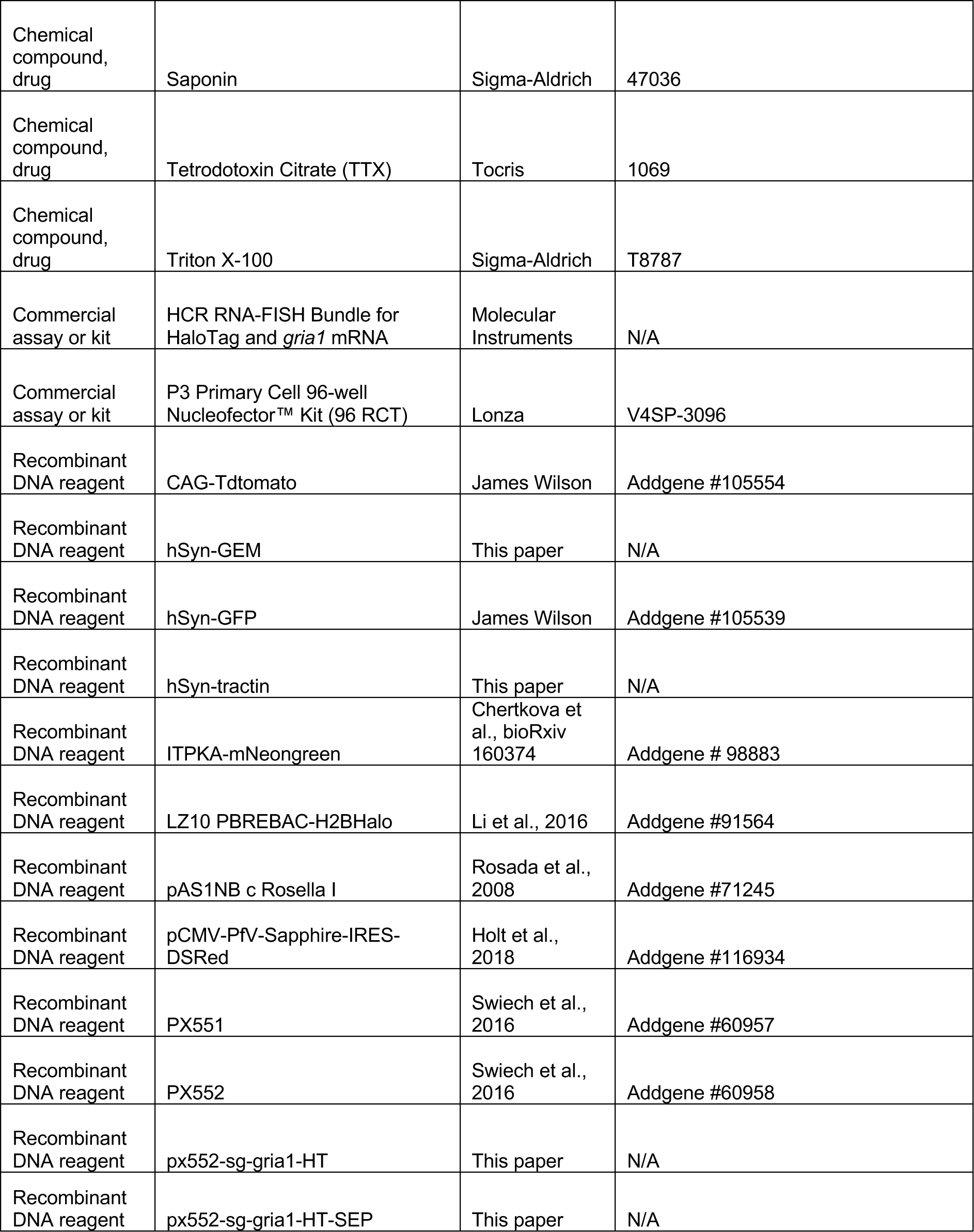
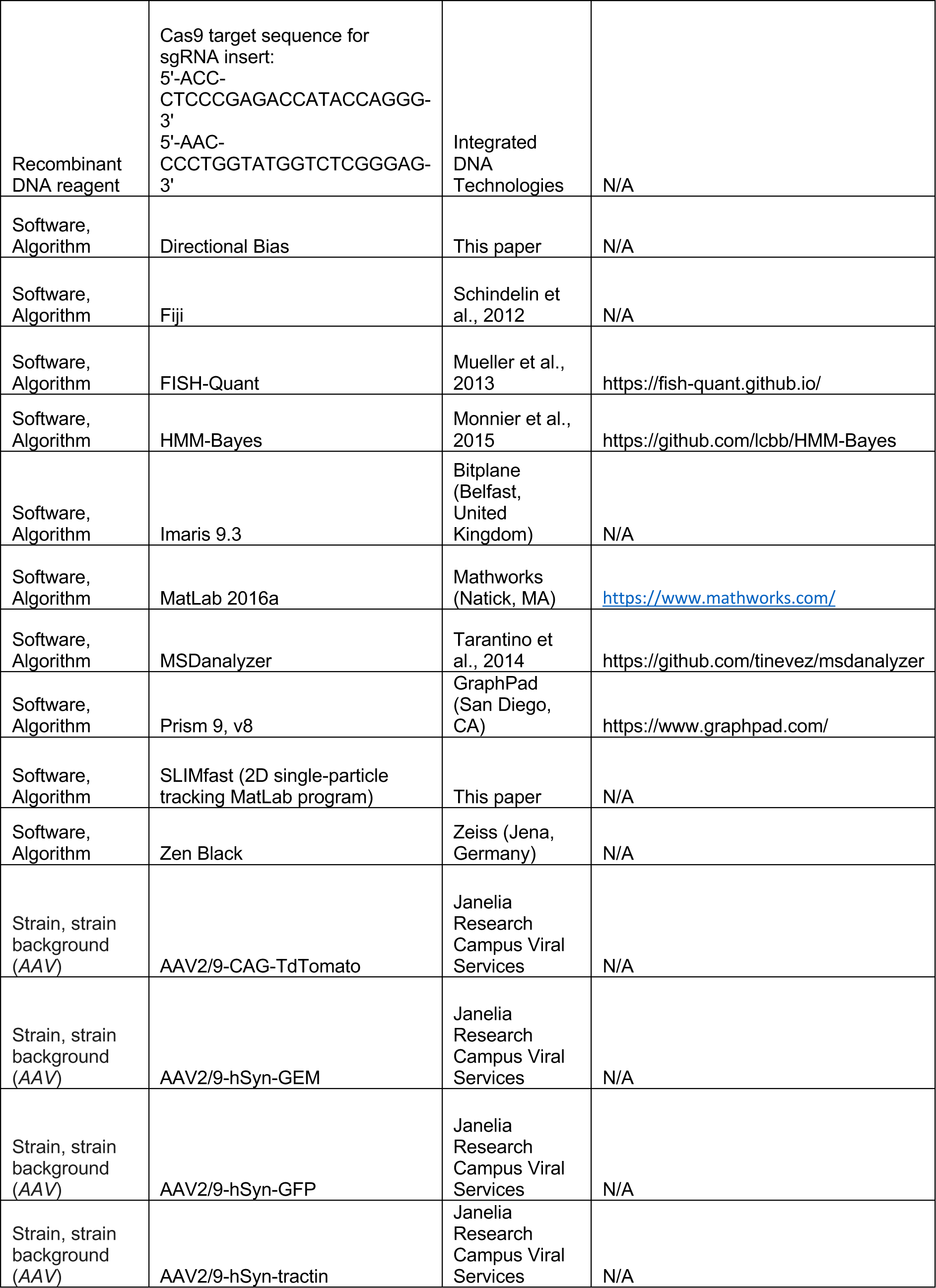
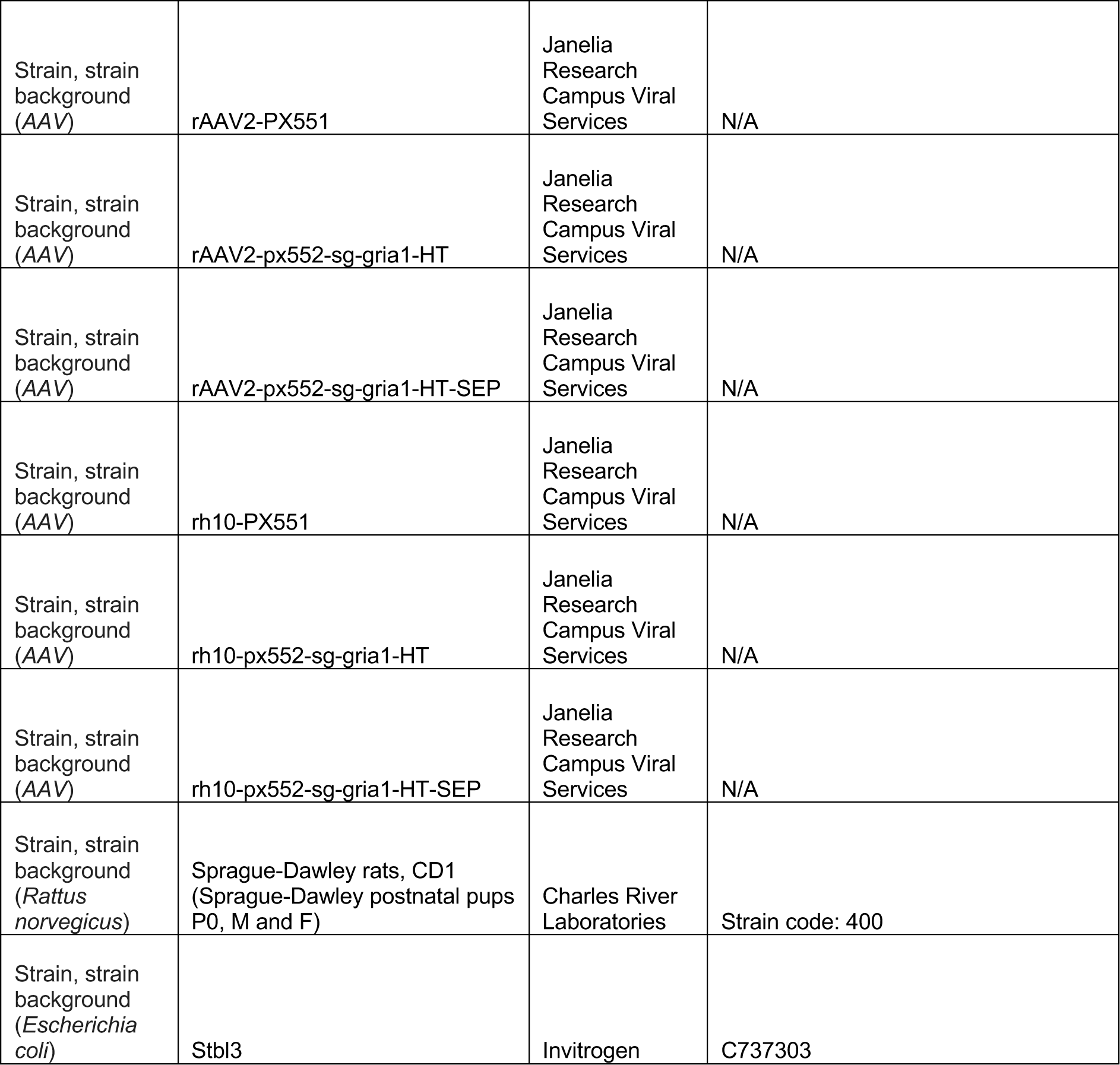

### Plasmid construction

To generate the HaloTag donor construct (px552-sg-Gria1-HT), miRFP670 was first amplified from pBAD/His-miRFP670 (Shcherbakova et al., 2016) and cloned into the KpnI and EcoRI sites of PX552 (Swiech et al., 2014). HaloTag was then amplified from LZ10 PBREBAC-H2BHalo (Liu et al., 2016) using primers that add glycine-serine linkers and *Gria1* sequence to be targeted by Cas9. This HaloTag amplicon was cloned into the XbaI site of PX552 to generate px552-Gria1-HT. To introduce a single guide RNA (sgRNA) insert in px552-Gria1-HT for Cas9 to target *Gria1*, we ordered the 20 bp Cas9 target sequence with 5’ overhangs (ACC and AAC) from Integrated DNA Technologies (IDT, Newark, NJ), and cloned the Cas9 target sequence into the SapI sites of px552-Gria1-HT (as previously described in Sweitch et al., 2014), creating px552-sg-Gria1-HT. For the HaloTag-SEP donor construct (px552-sg-Gria1-HT-SEP), HaloTag was amplified from LZ10 PBREBAC-H2BHalo and SEP was amplified from pAS1NB c Rosella I (Rosada et al., 2008). We used primers that add glycine-serine linkers and *Gria1* sequence to be targeted by Cas9 on the 5’-end of HaloTag and the 3’-end of SEP, and primers that introduce a short glycine-serine linker between HaloTag and SEP. The PCR products for HaloTag and SEP were fused using overlap extension PCR, and cloned into the MfeI and NheI sites of px552-sg- Gria1-HT to generate px552-sg-Gria1-HT-SEP. The Cas9 expression construct (PX551) was previously described (Swiech et al., 2014). To generate the tractin construct (hSyn-tractin), tractin-mNeongreen (also known as ITPKA-mNeongreen) was amplified from the ITPKA- mNeongreen construct (Cherkova et al., bioRxiv 160374) and cloned into the KpnI and BsrGI sites of px552-Gria1-HT. Additional sequences between the ITRs that are not related to the expression of tractin-mNeongreen (i.e., the sgRNA insert and the HaloTag donor sequence) were removed by digesting the construct with MluI and NheI and inserting a short double-stranded DNA oligo with 5’ overhangs that are complementary to MluI and NheI, creating hSyn-tractin.

The GEM construct (hSyn-GEM) was created by amplifying PfV from pCMV-PfV-Sapphire- IRES-DSRed (Holt et al., 2018), fusing PfV, HaloTag, and SNAP-tag by overlap extension PCR, and cloning the product into the KpnI and BsrGI sites of hSyn-tractin, creating hSyn-GEM. All plasmids were propagated in Stbl3 *e. coli* cells.

### Neuronal primary culture and electroporation

Dissociated hippocampal neurons were prepared from P0 Sprague-Dawley rat pups (Charles River). Hippocampi were dissected out and digested with papain in dissection solution (10 mM HEPES in Hanks’ balanced salt solution; HBSS). After digestion, the tissues were gently triturated in minimum essential media (MEM) with 10% fetal bovine serum and filtered with a 40 μm cell strainer. The cell density and viability were determined by labeling neurons with trypan blue and counting cells with a Countess 3 cell counter. To transfect neurons via electroporation, 500,000 neurons were resuspended in 20 μL complete P3 (P3 solution with supplement) and moved to a cuvette containing 0.5 ug of each plasmid to be transfected.

Samples were then electroporated using an Amaxa 4D-Nucleofector with settings CU-110. 80 μL of plating media (MEM with 10% fetal bovine serum, 28 mM glucose, 2.4 mM NaHCO3, 100 μg/mL transferrin, 25 μg/mL insulin, 2 mM L-glutamine) was added to the cuvette immediately after electroporation and samples were allowed to recover for 5 minutes at 37°c and 5% CO2. The electroporated sample was then removed from the cuvette and added to an Eppendorf tube with 500 μL plating media. Approximately, 50,000-75,000 cells were spread onto the coverslip of a poly-D-lysine (PDL) coated 10 mm MatTek dish. 6 hours later, cells were fed with 2 mL NbActiv4 neuronal culture media. Half of the neuronal culture media was removed and replaced with fresh NbActiv4 every week until the neurons were used.

### Adeno-associated virus (AAV) packaging and transduction

px552-sg-Gria1-HT, px552-sg-Gria1-HT-SEP, PX551, hSyn-tractin, hSyn-GEM, CAG-tdtomato, and hSyn-GFP were packaged into adeno-associated virus (AAV) by the Janelia Research Campus Viral Services Shared Resource. Briefly, HEK293T cells were transiently transfected with 84 µg of DNA at a ratio of pHelper plasmid:capsid plasmid:AAV construct = 3:2:5. Transfected cells were replenished with fresh serum- and phenol-free Dulbecco’s Modified Eagle Medium (DMEM) at 6-8 hours post-transfection and incubated for three days at 37°c and 5% CO2. AAVs were collected from both cells and supernatant and purified by two rounds of continuous cesium chloride density gradient. AAV preparations were dialyzed, concentrated to 100 µL, and sterilized by filtration. The final viral titers were measured by quantitative PCR (qPCR) on the inverted terminal repeats (ITRs). AAVs were pseudotyped with AAV2/9, SL1 (retro), or rh10 capsids. For AAV2/9-hSyn-tractin, AAV2/9-hSyn-GEM, AAV2/9-CAG-tdtomato, or AAV2/9-hSyn-GFP, 1x10^8^ genomic copies of virus was mixed into 50 uL of NbActiv4 and then added directly into neuronal cultures between 3-7 days in vitro (DIV3-7). Fluorescence signals from these reporters were detectable 3 days after transduction.

### Homology-independent targeted integration (HITI)

We used homology-independent targeted integration (HITI) to insert HaloTag or HaloTag-SEP into the endogenous loci of *Gria1* (see **Figure 1A** and **Figure 1-figure supplement 1**). px552- sg-Gria1-HT, px552-sg-Gria1-HT-SEP, and PX551 was delivered to cultured rat hippocampal neurons by either electroporation or AAV-mediated transduction. For electroporation, 0.5 ug of both px552-sg-Gria1-HT/HT-SEP and PX551 were mixed with neurons suspended in complete P3 buffer and electroporated as described above (see **Neuronal primary culture and electroporation**). For AAV-mediated transduction, 1x10^10^ genomic copies of rh10-px552-sg- Gria1-HT/HT-SEP virus and 1x10^9^ genomic copies of rh10-PX551 virus were mixed into 50 uL of NbActiv4 and then added directly into neuronal cultures at DIV3. HaloTag and HaloTag-SEP positive cells could be observed 7 days after transduction. To determine the HaloTag knock-in efficiency (HaloTag KI), we first counted the number of neurons expressing HaloTag and the number of neurons expressing the miRFP670 transfection marker (also expressed from px552- sg-Gria1-HT/HT-SEP) in a dish. We then calculated the knock-in efficiency by dividing the number of HaloTag^+^ neurons with the number of neurons that have been transfected with both px552-sg-Gria1-HT and PX551:

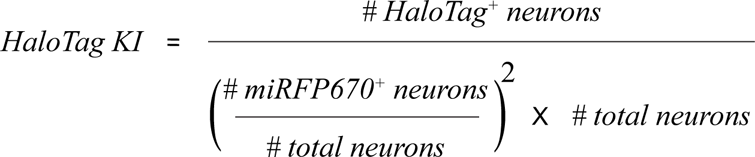

Because PX551 has no transfection marker, we assumed that the rate of PX551 transfection is equal to the rate of px552-sg-Gria1-HT transfection. Consequently, we extrapolate that the rate of co-transfection is equal to the rate of px552-sg-Gria1-HT transfection squared.

### Janelia Fluor (JF) dye labeling

Janelia Fluor 549 Halotag ligand (JF_549_-HTL), Janelia Fluor 646 Halotag ligand (JF_646_-HTL), and cell membrane impermeable Janelia Fluor 549 Halotag ligand (JF_549i_-HTL) were kind gifts from Dr. Luke Lavis. Dyes were reconstituted in DMSO and diluted to working concentrations of 10 nM (JF_549_-HTL and JF_549_i-HTL) or 20 nM (JF_646_-HTL) in imaging buffer (150 mM NaCl, 2 mM CaCl2, 2 mM MgCl2, 5 mM KCl, 10 mM HEPES, 30 mM D-glucose, pH 7.4). To label HaloTag expressed in rat hippocampal neurons, NbActiv4 was removed and replaced with imaging buffer containing working concentrations of JF dye-HTL. Neurons were incubated with JF dye-HTL for 30 minutes at 37°c and 5% CO2. Cells were then washed three times with imaging buffer and allowed to incubate for another 30 minutes at 37°c and 5% CO2. After the second incubation, cells were washed an additional three times with imaging buffer and then used for stimulation or imaging. For block-and-chase labeling experiments, neurons expressing HaloTag were first labeled with 20 nM JF_646_-HTL under 50 nM cycloheximide (CHX) in NbActiv4 for 1 hour at 37°c and 5% CO2. JF_646_-HTL and CHX were removed by rinsing neurons with NbActiv4 three times. Neurons were then incubated in NbActiv4 for 3-5 hours at 37°c and 5% CO2 to allow protein synthesis to recover. After the recovery period, neurons were labeled with 10 nM JF_549_- HTL in imaging buffer for 30 minutes at 37°c and 5% CO2. Finally, neurons were rinsed an additional three times with imaging buffer and stimulated or imaged.

### Immunofluorescence

Cultured rat hippocampal neurons were fixed between DIV 12 and 21. Neurons were fixed with fixation buffer (1x PBS, 4% PFA, 4% sucrose, 1 mM MgCl2, 0.1 mM CaCl2) for 20 minutes.

Fixation solution was quenched with 0.1 M glycine in PBS-MC (1xPBS, 1 mM MgCl2, 0.1 mM CaCl2) for 10 minutes. Neurons were then washed three times for 5 minutes each wash with PBS-MC. To label epitopes on the neuronal surface (e.g., the extracellular domain of GluA1), neurons were blocked for 1 hour with blocking buffer (5% normal goat serum in PBS-MC) and then labeled with primary antibodies in blocking buffer for 1 hour at room temperature. After labeling, neurons were washed three times for 5 minutes each wash with PBS-MC. Extracellular epitopes were labeled with secondary antibody in blocking buffer for 1 hour at room temperature. Lastly, neurons were washed three times for 5 minutes each with PBS. To label intracellular epitopes (e.g., Homer1 and PSD-95), neurons were simultaneously permeabilized and blocked with 0.1% saponin blocking buffer (0.1% saponin in blocking buffer), and then labeled with primary antibodies in 0.02% saponin blocking buffer overnight at 4°C. Neurons were washed three times for 10 minutes each wash with 0.02% saponin blocking buffer.

Intracellular epitopes were labeled with secondary antibodies in 0.02% saponin blocking buffer for 1 hour at room temperature. Neurons were then washed three times for 10 minutes each wash with 0.02% saponin blocking buffer and an additional three times for 5 minutes each wash with PBS. After labeling, neurons were mounted in Prolong Diamond Antifade Mountant with DAPI for 24 hours. Labeled samples were imaged with an inverted Carl Zeiss LSM 880 microscope equipped with 405/488/561/594/633 nm laser lines for illumination, and 2 multi-Alkali photomultiplier tubes (PMTs) and a 32-channel spectral gallium arsenide phosphide (GaAsP) PMT. QUASAR detection windows were adjusted optimally for each fluorophores. A 40x Plan Apochromat oil-immersion objective (NA=1.3) was used for all immunofluorescence imaging. The pinhole size was set to 1 airy unit (AU) based on the longest emission wavelength detected.

### Colocalization Analysis

To examine the colocalization of HaloTag, GluA1, and post-synaptic density markers, we performed intensity-based colocalization on antibody-labeled immunofluorescence images using Imaris. Images were first deconvolved using Zen Black and imported to Imaris. To perform colocalization analysis in a specific neuron of interest (e.g., a neuron expressing HaloTag), we used Imaris to create a surface object of the cell of interest and then masked signal outside of the surface object. We then applied an automatic intensity threshold using Otsu’s method – an established thresholding method that enables us to minimize subjective analysis – to each individual fluorescence channel to be used for colocalization analysis. Using these threshold levels, we created a new volume channel corresponding to the colocalizing voxels between the two fluorescence channels. We report both the Pearson’s coefficient (which indicate whether the intensity of two signals co-vary) and the Manders’ coefficients (which indicate whether two signals overlap).

### Hybridization chain reaction RNA fluorescence in situ hybridization (HCR RNA-FISH)

Probes targeting *Rattus norvegicus Gria1* and HaloTag mRNA were designed by Molecular Instruments. Cultured rat hippocampal neurons were fixed as described (see **Immunofluorescence**). Neurons were permeabilized with 0.1% Triton X-100 in PBS-MC for 15 minutes. Neurons were washed three times with PBS-MC for 5 minutes each wash and then two times with 2x SSC (0.3 M NaCl and 0.03 M sodium citrate) for 5 minutes each wash. Neurons were next incubated for 30 minutes at 37°C in 30% probe hybridization buffer (30% formamide, 5x SSC, 9 mM citric acid pH 6, 0.1% Tween-20, 50 μg/mL heparin, 1x Denhardt’s solution, 10% low molecular weight dextran sulfate). *Gria1* and HaloTag mRNA were then labeled with *Gria1-* and HaloTag-targeting hybridization probes in 30% probe hybridization buffer for 12 hours at 37°C. After hybridization, neurons were washed four times with 30% probe wash buffer (30% formamide, 5x SSC, 9 mM citric acid pH 6, 0.1% Tween-20, 50 μg/mL heparin) and three times with 5x SSCT (5x SSC, 0.1% Tween 20) for 5 minutes each wash at room temperature.

Neurons were then incubated with amplification buffer (5x SSC, 0.1% Tween-20, 10% low molecular weight dextran sulfate) for 30 minutes at room temperature. Hybridization probes were then amplified with amplification probes conjugated with Alexa 488 or Alexa 546 in amplification buffer for 45 minutes at room temperature. After amplification, neurons were wash two times with 5x SSCT for 5 minutes each, two times with 5x SSCT for 30 minutes each, and once with 5x SSCT for 5 minutes at room temperature. After HCR RNA-FISH labeling, cells were mounted in Prolong Diamond Antifade Mountant for 24 hours. Samples were imaged using an inverted Carl Zeiss LSM 880 microscope (see **Immunofluorescence**) with a 63x Plan Apochromat oil-immersion objective (NA=1.4).

### mRNA quantification and correlated localization analysis

Fluorescent puncta representative of individual mRNA were identified and counted using the FISH-Quant program for MATLAB (Mueller et al., 2013). First, neurons were identified using JF_646_-HTL labeled GluA1-HT and manually outlined. Overlapping neurons were removed from analysis. A Gaussian kernel filter was applied to HCR RNA-FISH images to enhance spot-like features. A pre-detection threshold was determined based on the dimmest and brightest pixel on each image and applied to the image. Additional false-positives were then removed based on their low quality score. Pre-detected spots were then fit with a 3D Gaussian function. Detected spots were then visually inspected for one neuronal cell body to determine if the detection parameters were adequate, after which the detection parameters were applied to the remaining cells in the image to quantify the number of spots in each cell. We used object-based colocalization to determine whether spots from two channels (i.e., HaloTag mRNA labeling and *Gria1* labeling) have correlated localization (**Figure 1D**). Two-channel images were first deconvolved with Zen Black and then imported to Imaris. mRNA in each channel were detected using the Imaris Spot Detection algorithm with a default quality filter for thresholding. Detected spots were visually inspected. We then examined the colocalization of thresholded spots from the two channels. A distance cutoff (i.e., the maximum distance within which two spots can be considered correlated in their localization) of 0.5 μm was used, as this corresponds to a physical distance of 1470 linearized basepairs (half the maximum distance in which probes from the two channels can be separated if they are both on the same mRNA – i.e., if *Gria1-*HaloTag is synthesized as a single mRNA). We report the percentage of spots in one channel that are within 0.5 mm of spots from the second channel and vice versa as a metric to determine whether the two labels have hybridized to a single mRNA (i.e., *Gria1-*HaloTag).

### Vesicle imaging and single-particle tracking

GluA1-HT and GluA1-HT-SEP vesicles were imaged with a Nikon Eclipse TiE inverted microscope equipped with 405/488/561/642 nm laser lines, three iXon Ultra 897 electron multiplying charge-coupled device (EMCCD) cameras connected via a tri-cam splitter for simultaneous multicolor acquisition, an automatic total internal reflection fluorescence (TIRF) illuminator, and a perfect focusing system. A Tokai Hit Stage Top Incubator was used to maintain constant environmental conditions of 5% CO2 and 37°C during imaging. A 100x TIRF Apochromat oil-immersion objective (NA=1.49) was used for imaging. To image GluA1-HT and GluA1-HT-SEP vesicles, we employed highly inclined and laminated sheet (HILO) illumination. Specifically, the TIRF illuminator was adjusted to deliver the laser beam with an incident angle smaller than the total internal reflection angle, which generated a highly inclined light sheet centered on the focal plane. Compared with standard epifluorescence illumination, HILO illumination reduces background from out-of-focus excitation. Images were acquired at 20 Hz.

The *xy* pixel size for all GluA1-HT and GluA1-HT-SEP vesicle tracking experiments was 160 nm. 2D single-particle tracking was performed with a custom MATLAB program. Single molecule localization (x,y) was obtained through 2D Gaussian fitting and tracking was based on the multiple-target tracing (MTT) algorithm. The localization and tracking parameters in SPT experiments are listed in the **Table 1**. The resulting tracks were manually curated, and GluA1-HT and GluA1-HT-SEP vesicle tracks were identified based on the position of the vesicle in the dendrite, its motion type, and its bleaching characteristics (**Figure 2-figure supplement 1**).

**Table 1.** Localization and tracking parameters for the MTT program.

|  |  |  |
| --- | --- | --- |
| Localization error | 1E-6 |  |
| Deflation loops | 3 |  |
| Maximum number competitors | 3 |  |
| Maximum diffusion coefficient ($\mu\text{m}^2/\text{s}$) | 1 | For GluA1-HT |
| Maximum diffusion coefficient ($\mu\text{m}^2/\text{s}$) | 3 | For GEM |

### Hidden Markov modeling with Bayesian model selection (HMM-Bayes) analysis

The diffusion and transport states of individual GluA1-HT and GluA1-HT-SEP trajectories were analyzed by HMM-Bayes MATLAB program using default parameters. HMM-Bayes classifies each jumping step in a trajectory as either diffusion or active transport and also calculates the diffusion coefficient and velocity of each step. Active transport (DV) is modeled as directed motion (V) with Brownian diffusion (D) according to the equation:

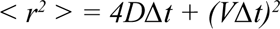

We defined the maximum number of unique motion states that can be inferred for a trajectory to be 3 (i.e., a trajectory can be assigned 1, 2, or 3 unique states). Diffusion coefficients determined by HMM-Bayes analysis were validated by comparing these values to diffusion coefficients for the same trajectories determined with the standard MSD curve fitting by MSDanalyzer in MATLAB (minimal fitting of R^2^ = 0.8; see **Figure 2-figure supplement 2** for more detail).

### Chemical stimulation

To determine whether chemical LTP (cLTP) increased the concentration of GluA1-HT on neuronal surfaces, we washed cultured neurons three times with stimulation buffer (150 mM NaCl, 2 mM CaCl2, 5 mM KCl, 10 mM HEPES, 30 mM D-glucose, 20 mM bicuculline, 1 μM strychnine, pH 7.4) and then stimulated neurons with 0.2 mM glycine in stimulation buffer for 15 minutes at 37°c and 5% CO2. Neurons were rinsed twice with wash buffer (150 mM NaCl, 2 mM CaCl2, 2 mM MgCl2, 5 mM KCl, 10 mM HEPES, 30 mM D-glucose, 20 mM bicuculline, 1 μM strychnine, pH 7.4) and then incubated with 25 nM cell impermeable JF_549_-HTL (JF_549i_-HTL in wash buffer for 30 minutes. Cells were then washed three times with wash buffer and fixed as described (see **Immunofluorescence)**. For live-cell GluA1-HT vesicle tracking experiments, the dish was placed onto the stage of a Nikon Eclipse TiE microscope after three washes with stimulation buffer (see **Vesicle imaging conditions and single-particle tracking** for imaging setup). After a desired field of view was identified, stimulation buffer was exchanged for 0.2 mM glycine in stimulation buffer. A desired Z section was identified and imaging commenced immediately. For Jasplakinolide (JSP) stimulation experiments, neurons were first washed with imaging buffer and placed onto the Nikon Eclipse TiE microscope. After a desired field of view was identified, JSP in imaging buffer was added to the dish to a final concentration of 0.25 μM. After 10 minutes of incubation with JSP, we commenced imaging. For experiments with Latrunculin A (LatA), neurons were pretreated with 1 μM LatA. In addition, LatA was added to every buffer for stimulation and imaging. For myosin V and VI inhibition, a cocktail of 4 μM Pentabromopseudilin (PBP), 10 μM MyoVin-1, and 4 μM of 2,4,6-Triiodophenol (TIP) was added to stimulation buffer during chemical stimulation and photostimulation.

### Photostimulation

Neurons were first washed three times with stimulation buffer. Stimulation buffer was replaced with 2 mM MNI-caged glutamate in stimulation buffer. We placed the dish onto the Nikon Eclipse TiE microscope and identified neurons expressing GFP and GluA1-HT (or just GluA1- HT-SEP for exocytosis experiments; see **Vesicle imaging conditions and single-particle tracking** for imaging setup). We selected dendritic spines to be targeted for glutamate uncaging-evoked sLTP based on their morphology in the FITC channel. We then positioned the 405 nm uncaging laser 1 μm from the tip of the dendritic spine head. sLTP was stimulated by firing the 405 nm uncaging laser at 2 Hz for 50 seconds with a pulse width of 1 ms. After photostimulation, we checked that the dendritic spine head of interest had expanded in area in the FITC channel. In order to examine the effect of sLTP on GluA1-HT vesicle motion, we performed timelapse imaging on GluA1-HT vesicles in the TRITC channel (see **Vesicle imaging and single-particle tracking**) immediately before and after sLTP stimulation. To account for photobleaching and determine the adjusted number of vesicles after photostimulation, we determined the vesicle counts before and after stimulation in no MNI controls. We then used the fold loss in vesicles after photostimulation in no MNI controls to correct the number of vesicles after photostimulation across all conditions.

### F-actin imaging

To examine the effect of cLTP on the length of F-actin fibers in the dendritic shaft, cultured rat hippocampal neurons expressing tractin were imaged using an inverted Carl Zeiss LSM 880 microscope (see **Immunofluorescence**) with a 63x Plan Apochromat oil-immersion objective (NA=1.4). To achieve higher resolution, emitted photons were collected with a 32-channel spectral GaAsP PMT for Airyscan processing. As these experiments were performed on live cells, a PeCon Incubator XL enclosure and Lab-TekTM S1 heating insert and CO2 lid were used to maintain constant environmental conditions of 5% CO2 and 37°C during imaging. Prior to imaging, neurons expressing tractin were washed three times with stimulation buffer (see **Chemical stimulation**) and placed onto the Zeiss LSM 880 stage. A dendrite of interest expressing tractin was identified and a z-stack of the dendrite acquired. After acquisition of the dendrite before cLTP (pre-cLTP), stimulation buffer was exchanged for 0.2 mM glycine in stimulation buffer. After 15 minutes, a z-stack of the dendrite (post-cLTP) was acquired with identical imaging parameters. Airyscan processing was applied to both the pre- and post-cLTP images. The F-actin skeletons were then quantified as described in **Figure 4-figure supplement 2**.

### Radius of confinement and directional bias analysis

To determine the directional bias of a trajectory using the angle analysis, we implemented a custom MATLAB program. Specifically, for a given trajectory, we defined the radius of confinement as the distance from the centroid of a trajectory to the furthest point in the trajectory to the centroid. Next, we created a mask around the dendrite in which the trajectory is localized and set the dendritic path along the length of the dendrite. We then determined the point on the dendritic path that is closest to the centroid of the trajectory. The angle, theta 0, created by these three points (the point on the dendritic path closest to the centroid, the centroid itself, and the point on the trajectory furthest from the centroid) indicates whether the motion of the trajectory is biased *towards* the length of the dendritic path or *along* the length of the dendritic path. Theta ∼ 0° indicates that the motion of the trajectory is biased perpendicular to the dendritic path (i.e., lateral motion) while theta ∼ 90° indicates that the motion of the trajectory is biased parallel to the dendritic path (i.e., longitudinal motion).

### Electrophysiology

Whole-cell voltage-clamp recordings were performed to determine the preservation of GluA1- HT and GluA1-HT-SEP functionality. Cultured hippocampal neurons were assessed at DIV7-8. Labeling GluA1-HT/HT-SEP with JF_549_-HTL was performed 12 hours prior to patch clamp experiments. GluA1-HT/ HT-SEP positive and negative neurons were determined via epifluorescence. Whole-cell configuration was achieved using pipettes pulled and polished to a resistance of 2-5 MΩ. SylGard 184 coating was applied to reduce pipette capacitance. Pipette solution composition included 120 mM Cs-MES; 5 mM NaCl, 10 mM TEA-Cl, 5 mM Lidocaine, 1.1 mM EGTA, 10 mM HEPES, 0.3 mM Na-GTP, 4 mM Mg-ATP. External solution contained 135 mM NaCl, 3 mM KCl, 2 mM MgCl2, 1.5 mM CaCl2, 10 mM TEA-Cl, 10 mM HEPES, 10 mM Glucose. The external solution was supplemented with 200 nM TTX and 10 µM AP5. Cell capacitance was estimated and corrected to 80% with a 10 µsec lag. Gap free voltage clamp recordings were performed at 100 kHz acquisition frequency with a 1.0 kHz Bessel filter. Voltage was clamped to -70 mV, correcting for liquid junction potential. Following baseline measurements, 100 µM glutamate in pH adjusted external solution was locally perfused for 5 seconds using a fused silicate pipe positioned within 100-200 µm of the neuron recorded. The neuron then recovered with continued bath perfusion and local perfusion of glutamate-free external solution. For analysis, the gap-free epochs were reduced 50 fold. The data was normalized to current density via cell capacitance estimations. Peak current density for each response was determined.

### Statistical analysis

All statistical tests and correlation analysis were performed in GraphPad Prism, version 8. Trajectories for each condition were pooled from at least 6 experiments (one dish of neurons per experiment). 2-3 dishes of neurons for each condition were extracted from a single animal, and at least 3 animals were used for a condition. We found no significant difference in the fraction of vesicles exhibiting active transport and in the average diffusion coefficients of vesicles exhibiting diffusion between dissections under control conditions and under cLTP treatment (**Figure 2-figure supplement 2**), demonstrating that variation stemming from technical and biological replicates is less than variation due to experimental conditions.

## Acknowledgements

We thank members of the O’Shea lab and our colleagues Dr. Heejun Choi, Dr. Kyle Harrington, Dr. Dong-woo Hwang, Dr. Xi Long, and Dr. Young J. Yoon for useful discussions; Dr. David E. Clapham and Dr. Pietro De Camilli for their helpful suggestions and feedback on this manuscript; Dr. Luke D. Lavis and Jonathan B. Grimm for providing the Janelia Fluor dyes and guidance on their usage; Dr. Damien Alcor from the Janelia Research Campus (JRC) Light Microscopy Center and Dr. Andrian Gutu for their assistance with imaging platforms; Dr. Hyun Ah Yi and Dr. Alina D. Gutu from the JRC Viral Tools facility for packaging AAVs; and Michelle Quiambao for administrative assistance. This work was supported by the Howard Hughes Medical Institute.

## References

1. Abbott, L. F., & Nelson, S. B. (2000). Synaptic plasticity: taming the beast. Nature neuroscience, 3 *Suppl*, 1178–1183.

2. Bliss, T. V., & Collingridge, G. L. (2013). Expression of NMDA receptor-dependent LTP in the hippocampus: bridging the divide. Molecular brain, 6, 5.

3. Bliss, T. V., & Cooke, S. F. (2011). Long-term potentiation and long-term depression: a clinical perspective. *Clinics (Sao Paulo*, Brazil), 66 Suppl 1(Suppl 1), 3–17.

4. Bosch, M., Castro, J., Saneyoshi, T., Matsuno, H., Sur, M., & Hayashi, Y. (2014). Structural and molecular remodeling of dendritic spine substructures during long-term potentiation. Neuron, 82(2), 444–459.

5. Bowen, A. B., Bourke, A. M., Hiester, B. G., Hanus, C., & Kennedy, M. J. (2017). Golgi-independent secretory trafficking through recycling endosomes in neuronal dendrites and spines. eLife, 6, e27362.

6. Chater, T. E., & Goda, Y. (2014). The role of AMPA receptors in postsynaptic mechanisms of synaptic plasticity. Frontiers in cellular neuroscience, 8, 401.

7. Choi, H., Schwarzkopf, M., Fornace, M. E., Acharya, A., Artavanis, G., Stegmaier, J., Cunha, A., & Pierce, N. A. (2018). Third-generation *in situ* hybridization chain reaction: multiplexed, quantitative, sensitive, versatile, robust. *Development (Cambridge*, England), 145(12), dev165753.

8. Choquet, D., & Opazo, P. (2022). The role of AMPAR lateral diffusion in memory. Seminars in cell & developmental biology, 125, 76–83.

9. Citri, A., & Malenka, R. C. (2008). Synaptic plasticity: multiple forms, functions, and mechanisms. Neuropsychopharmacology : official publication of the American College of Neuropsychopharmacology, 33(1), 18–41.

10. Correia, S. S., Bassani, S., Brown, T. C., Lisé, M. F., Backos, D. S., El-Husseini, A., Passafaro, M., & Esteban, J. A. (2008). Motor protein-dependent transport of AMPA receptors into spines during long-term potentiation. Nature neuroscience, 11(4), 457–466.

11. Craig, A. M., Blackstone, C. D., Huganir, R. L., & Banker, G. (1993). The distribution of glutamate receptors in cultured rat hippocampal neurons: postsynaptic clustering of AMPA-selective subunits. Neuron, 10(6), 1055–1068.

12. Delarue, M., Brittingham, G. P., Pfeffer, S., Surovtsev, I. V., Pinglay, S., Kennedy, K. J., Schaffer, M., Gutierrez, J. I., Sang, D., Poterewicz, G., Chung, J. K., Plitzko, J. M., Groves, J. T., Jacobs-Wagner, C., Engel, B. D., & Holt, L. J. (2018). mTORC1 Controls Phase Separation and the Biophysical Properties of the Cytoplasm by Tuning Crowding. Cell, 174(2), 338–349.e20.

13. Diering, G. H., & Huganir, R. L. (2018). The AMPA Receptor Code of Synaptic Plasticity. Neuron, 100(2), 314–329.

14. Ellis-Davies G. C. (2007). Caged compounds: photorelease technology for control of cellular chemistry and physiology. Nature methods, 4(8), 619–628.

15. Esteves da Silva, M., Adrian, M., Schätzle, P., Lipka, J., Watanabe, T., Cho, S., Futai, K., Wierenga, C. J., Kapitein, L. C., & Hoogenraad, C. C. (2015). Positioning of AMPA Receptor- Containing Endosomes Regulates Synapse Architecture. Cell reports, 13(5), 933–943.

16. Fukazawa, Y., Saitoh, Y., Ozawa, F., Ohta, Y., Mizuno, K., & Inokuchi, K. (2003). Hippocampal LTP is accompanied by enhanced F-actin content within the dendritic spine that is essential for late LTP maintenance in vivo. Neuron, 38(3), 447–460.

17. Goo, M. S., Sancho, L., Slepak, N., Boassa, D., Deerinck, T. J., Ellisman, M. H., Bloodgood, B. L., & Patrick, G. N. (2017). Activity-dependent trafficking of lysosomes in dendrites and dendritic spines. The Journal of cell biology, 216(8), 2499–2513.

18. Grimm, J. B., Muthusamy, A. K., Liang, Y., Brown, T. A., Lemon, W. C., Patel, R., Lu, R., Macklin, J. J., Keller, P. J., Ji, N., & Lavis, L. D. (2017). A general method to fine-tune fluorophores for live-cell and in vivo imaging. Nature methods, 14(10), 987–994.

19. Groc, L., & Choquet, D. (2020). Linking glutamate receptor movements and synapse function. *Science (New York*, N.Y.), 368(6496), eaay4631.

20. Hammer, J. A., 3rd, & Sellers, J. R. (2011). Walking to work: roles for class V myosins as cargo transporters. Nature reviews. Molecular cell biology, 13(1), 13–26.

21. Hangen, E., Cordelières, F. P., Petersen, J. D., Choquet, D., & Coussen, F. (2018). Neuronal Activity and Intracellular Calcium Levels Regulate Intracellular Transport of Newly Synthesized AMPAR. Cell reports, 24(4), 1001–1012.e3.

22. Herring, B. E., & Nicoll, R. A. (2016). Long-Term Potentiation: From CaMKII to AMPA Receptor Trafficking. Annual review of physiology, 78, 351–365.

23. Hoerndli, F. J., Maxfield, D. A., Brockie, P. J., Mellem, J. E., Jensen, E., Wang, R., Madsen, D. M., & Maricq, A. V. (2013). Kinesin-1 regulates synaptic strength by mediating the delivery, removal, and redistribution of AMPA receptors. Neuron, 80(6), 1421–1437.

24. Hoogenraad, C. C., Milstein, A. D., Ethell, I. M., Henkemeyer, M., & Sheng, M. (2005). GRIP1 controls dendrite morphogenesis by regulating EphB receptor trafficking. Nature neuroscience, 8(7), 906–915.

25. Jaqaman, K., Loerke, D., Mettlen, M., Kuwata, H., Grinstein, S., Schmid, S. L., & Danuser, G. (2008). Robust single-particle tracking in live-cell time-lapse sequences. Nature methods, 5(8), 695–702.

26. Kandel E. R., Schwartz J. H., Jessell T. M., Siegelbaum S. A., Hudspeth A. J. (2013). Principles of Neural Science, 5th Edn New York, NY: McGraw-Hill Medical.

27. Katrukha, E. A., Mikhaylova, M., van Brakel, H. X., van Bergen En Henegouwen, P. M., Akhmanova, A., Hoogenraad, C. C., & Kapitein, L. C. (2017). Probing cytoskeletal modulation of passive and active intracellular dynamics using nanobody-functionalized quantum dots. Nature communications, 8, 14772.

28. Kopec, C. D., Li, B., Wei, W., Boehm, J., & Malinow, R. (2006). Glutamate receptor exocytosis and spine enlargement during chemically induced long-term potentiation. The Journal of neuroscience : the official journal of the Society for Neuroscience, 26(7), 2000–2009.

29. Lavoie-Cardinal, F., Bilodeau, A., Lemieux, M., Gardner, M. A., Wiesner, T., Laramée, G., Gagné, C., & De Koninck, P. (2020). Neuronal activity remodels the F-actin based submembrane lattice in dendrites but not axons of hippocampal neurons. Scientific reports, 10(1), 11960.

30. Lin, D. T., Makino, Y., Sharma, K., Hayashi, T., Neve, R., Takamiya, K., & Huganir, R. L. (2009). Regulation of AMPA receptor extrasynaptic insertion by 4.1N, phosphorylation and palmitoylation. Nature neuroscience, 12(7), 879–887.

31. Lisman, J., Yasuda, R., & Raghavachari, S. (2012). Mechanisms of CaMKII action in long-term potentiation. Nature reviews. Neuroscience, 13(3), 169–182.

32. Liu, H., Dong, P., Ioannou, M. S., Li, L., Shea, J., Pasolli, H. A., Grimm, J. B., Rivlin, P. K., Lavis, L. D., Koyama, M., & Liu, Z. (2018). Visualizing long-term single-molecule dynamics in vivo by stochastic protein labeling. Proceedings of the National Academy of Sciences of the United States of America, 115(2), 343–348.

33. Magee, J. C., & Grienberger, C. (2020). Synaptic Plasticity Forms and Functions. Annual review of neuroscience, 43, 95–117.

34. Makino, H., & Malinow, R. (2009). AMPA receptor incorporation into synapses during LTP: the role of lateral movement and exocytosis. Neuron, 64(3), 381–390.

35. Matsuzaki, M., Honkura, N., Ellis-Davies, G. C., & Kasai, H. (2004). Structural basis of long-term potentiation in single dendritic spines. Nature, 429(6993), 761–766.

36. Meunier, F. A., & Gutiérrez, L. M. (2016). Captivating New Roles of F-Actin Cortex in Exocytosis and Bulk Endocytosis in Neurosecretory Cells. Trends in neurosciences, 39(9), 605– 613.

37. Molnár E. (2011). Long-term potentiation in cultured hippocampal neurons. Seminars in cell & developmental biology, 22(5), 506–513.

38. Monnier, N., Barry, Z., Park, H. Y., Su, K. C., Katz, Z., English, B. P., Dey, A., Pan, K., Cheeseman, I. M., Singer, R. H., & Bathe, M. (2015). Inferring transient particle transport dynamics in live cells. Nature methods, 12(9), 838–840.

39. Mueller, F., Senecal, A., Tantale, K., Marie-Nelly, H., Ly, N., Collin, O., Basyuk, E., Bertrand, E., Darzacq, X., & Zimmer, C. (2013). FISH-quant: automatic counting of transcripts in 3D FISH images. Nature methods, 10(4), 277–278.

40. Nettesheim, G., Nabti, I., Murade, C.U. et al. Macromolecular crowding acts as a physical regulator of intracellular transport. Nat. Phys. 16, 1144–1151 (2020).

41. Okamoto, K., Nagai, T., Miyawaki, A., & Hayashi, Y. (2004). Rapid and persistent modulation of actin dynamics regulates postsynaptic reorganization underlying bidirectional plasticity. Nature neuroscience, 7(10), 1104–1112.

42. Opazo, P., Labrecque, S., Tigaret, C. M., Frouin, A., Wiseman, P. W., De Koninck, P., & Choquet, D. (2010). CaMKII triggers the diffusional trapping of surface AMPARs through phosphorylation of stargazin. Neuron, 67(2), 239–252.

43. Opazo, P., Sainlos, M., & Choquet, D. (2012). Regulation of AMPA receptor surface diffusion by PSD-95 slots. Current opinion in neurobiology, 22(3), 453–460.

44. Papadopulos, A., Gomez, G. A., Martin, S., Jackson, J., Gormal, R. S., Keating, D. J., Yap, A. S., & Meunier, F. A. (2015). Activity-driven relaxation of the cortical actomyosin II network synchronizes Munc18-1-dependent neurosecretory vesicle docking. Nature communications, 6, 6297.

45. Park M. (2018). AMPA Receptor Trafficking for Postsynaptic Potentiation. Frontiers in cellular neuroscience, 12, 361.

46. Patterson, M. A., Szatmari, E. M., & Yasuda, R. (2010). AMPA receptors are exocytosed in stimulated spines and adjacent dendrites in a Ras-ERK-dependent manner during long-term potentiation. Proceedings of the National Academy of Sciences of the United States of America, 107(36), 15951–15956.

47. Penn, A. C., Zhang, C. L., Georges, F., Royer, L., Breillat, C., Hosy, E., Petersen, J. D., Humeau, Y., & Choquet, D. (2017). Hippocampal LTP and contextual learning require surface diffusion of AMPA receptors. Nature, 549(7672), 384–388.

48. Rudolf, R., Bittins, C. M., & Gerdes, H. H. (2011). The role of myosin V in exocytosis and synaptic plasticity. Journal of neurochemistry, 116(2), 177–191.

49. Schätzle, P., Esteves da Silva, M., Tas, R. P., Katrukha, E. A., Hu, H. Y., Wierenga, C. J., Kapitein, L. C., & Hoogenraad, C. C. (2018). Activity-Dependent Actin Remodeling at the Base of Dendritic Spines Promotes Microtubule Entry. Current biology : CB, 28(13), 2081–2093.e6.

50. Schell, M. J., Erneux, C., & Irvine, R. F. (2001). Inositol 1,4,5-trisphosphate 3-kinase A associates with F-actin and dendritic spines via its N terminus. The Journal of biological chemistry, 276(40), 37537–37546.

51. Setou, M., Seog, D. H., Tanaka, Y., Kanai, Y., Takei, Y., Kawagishi, M., & Hirokawa, N. (2002). Glutamate-receptor-interacting protein GRIP1 directly steers kinesin to dendrites. Nature, 417(6884), 83–87.

52. Shepherd, J. D., & Huganir, R. L. (2007). The cell biology of synaptic plasticity: AMPA receptor trafficking. Annual review of cell and developmental biology, 23, 613–643.

53. Shi, S. H., Hayashi, Y., Petralia, R. S., Zaman, S. H., Wenthold, R. J., Svoboda, K., & Malinow, R. (1999). Rapid spine delivery and redistribution of AMPA receptors after synaptic NMDA receptor activation. *Science (New York*, N.Y.), 284(5421), 1811–1816.

54. Sood, P., Murthy, K., Kumar, V., Nonet, M. L., Menon, G. I., & Koushika, S. P. (2018). Cargo crowding at actin-rich regions along axons causes local traffic jams. *Traffic (Copenhagen*, Denmark), 19(3), 166–181.

55. Suzuki, K., Tsunekawa, Y., Hernandez-Benitez, R., Wu, J., Zhu, J., Kim, E. J., Hatanaka, F., Yamamoto, M., Araoka, T., Li, Z., Kurita, M., Hishida, T., Li, M., Aizawa, E., Guo, S., Chen, S., Goebl, A., Soligalla, R. D., Qu, J., Jiang, T., … Belmonte, J. C. (2016). In vivo genome editing via CRISPR/Cas9 mediated homology-independent targeted integration. Nature, 540(7631), 144–149.

56. Tao-Cheng, J. H., Crocker, V. T., Winters, C. A., Azzam, R., Chludzinski, J., & Reese, T. S. (2011). Trafficking of AMPA receptors at plasma membranes of hippocampal neurons. The Journal of neuroscience : the official journal of the Society for Neuroscience, 31(13), 4834– 4843.

57. Tardin, C., Cognet, L., Bats, C., Lounis, B., & Choquet, D. (2003). Direct imaging of lateral movements of AMPA receptors inside synapses. The EMBO journal, 22(18), 4656–4665.

58. van Bommel, B., Konietzny, A., Kobler, O., Bär, J., & Mikhaylova, M. (2019). F-actin patches associated with glutamatergic synapses control positioning of dendritic lysosomes. The EMBO journal, 38(15), e101183.

59. Wagner, W., Lippmann, K., Heisler, F. F., Gromova, K. V., Lombino, F. L., Roesler, M. K., Pechmann, Y., Hornig, S., Schweizer, M., Polo, S., Schwarz, J. R., Eilers, J., & Kneussel, M. (2019). Myosin VI Drives Clathrin-Mediated AMPA Receptor Endocytosis to Facilitate Cerebellar Long-Term Depression. Cell reports, 28(1), 11–20.e9.

60. Wang, Z., Edwards, J. G., Riley, N., Provance, D. W., Jr, Karcher, R., Li, X. D., Davison, I. G., Ikebe, M., Mercer, J. A., Kauer, J. A., & Ehlers, M. D. (2008). Myosin Vb mobilizes recycling endosomes and AMPA receptors for postsynaptic plasticity. Cell, 135(3), 535–548.

