## Supplemental Information for "Plasticity-induced actin polymerization in the dendritic shaft regulates intracellular AMPA receptor trafficking"

Figure 1-figure supplement 1

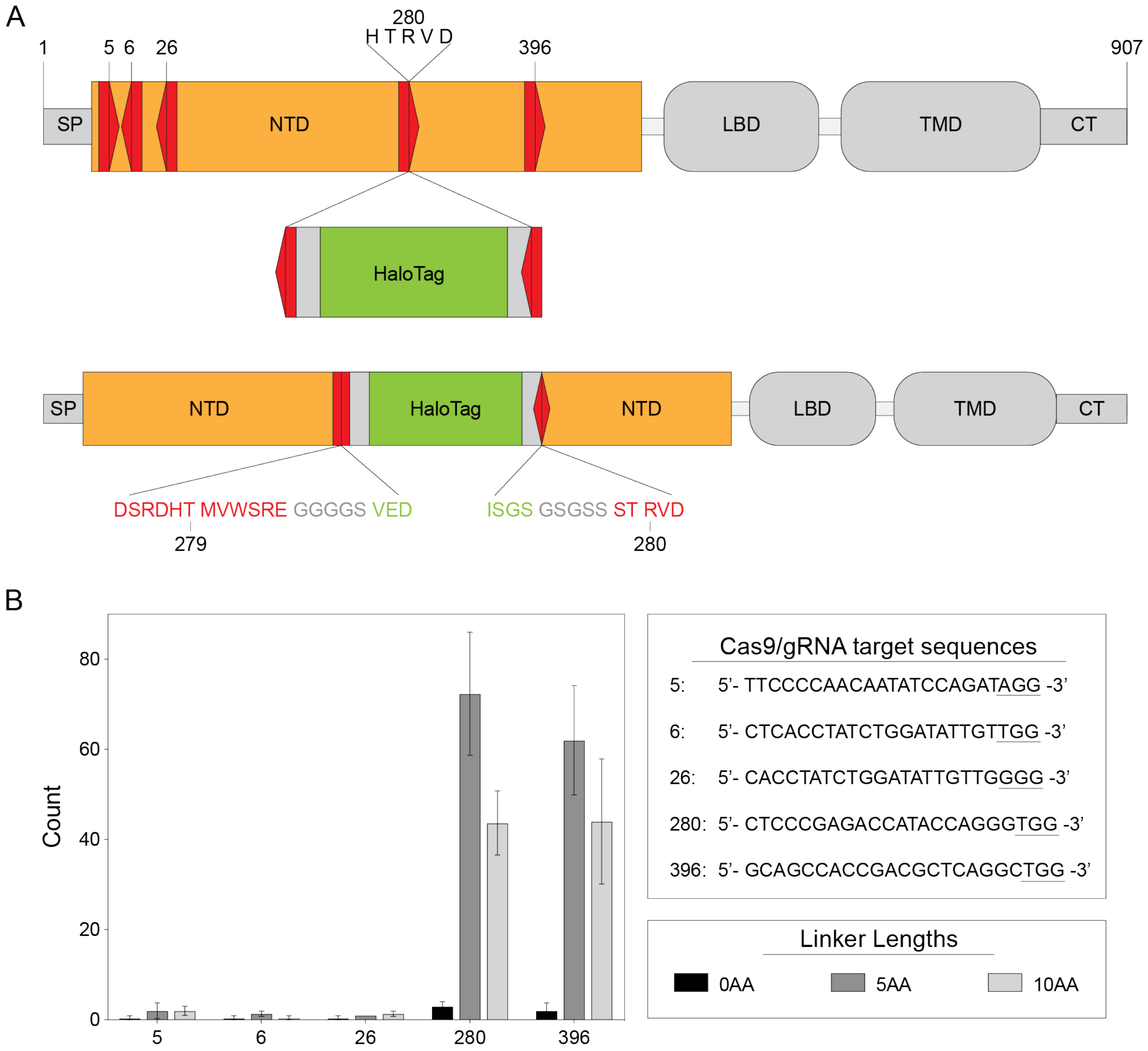

**Figure 1-figure supplement 1. Successful insertion of HaloTag into *Gria1* depends on target site and linker length.**

(A) Schematic of GluA1 protein domains. SP, signal peptide; NTD, amino terminal domain; LBD, ligand binding domain; TMD, transmembrane domain; CT, C-terminus. The NTD and LBD are extracellular. Red pentagons represent the relative locations of attempted Cas9/guide RNA (gRNA) target sequences. Direction indicates whether the upper or lower DNA strand of *Gria1* was targeted (pointing right or left, respectively). Number indicates amino acid position. Copies of Cas9/gRNA target sequence flanking HaloTag sequence face the opposite direction as the genomic *Gria1* Cas9/gRNA target sequence. When HaloTag is inserted in the correct orientation, the Cas9/gRNA target sequence will not be restored, preventing Cas9 from cutting the same sites after repair. Gray rectangles flanking HaloTag represent peptide linkers.

(B) Bar graph: mean number of cells with HaloTag knock-ins for different insertion sites and with different peptide linker lengths. n=3 transfections for each Cas9/gRNA target sequence and linker length. Error bars represent standard deviation. Top right panel: Cas9/gRNA target sequences with the protospacer adjacent motif (PAM) sequence underlined. Lower right panel: different linker lengths tested. Insertion at 280R with a linker length of 5 amino acids was used for all GluA1 tagging experiments because it has the highest knock-in efficiency.

Figure 1-figure supplement 2

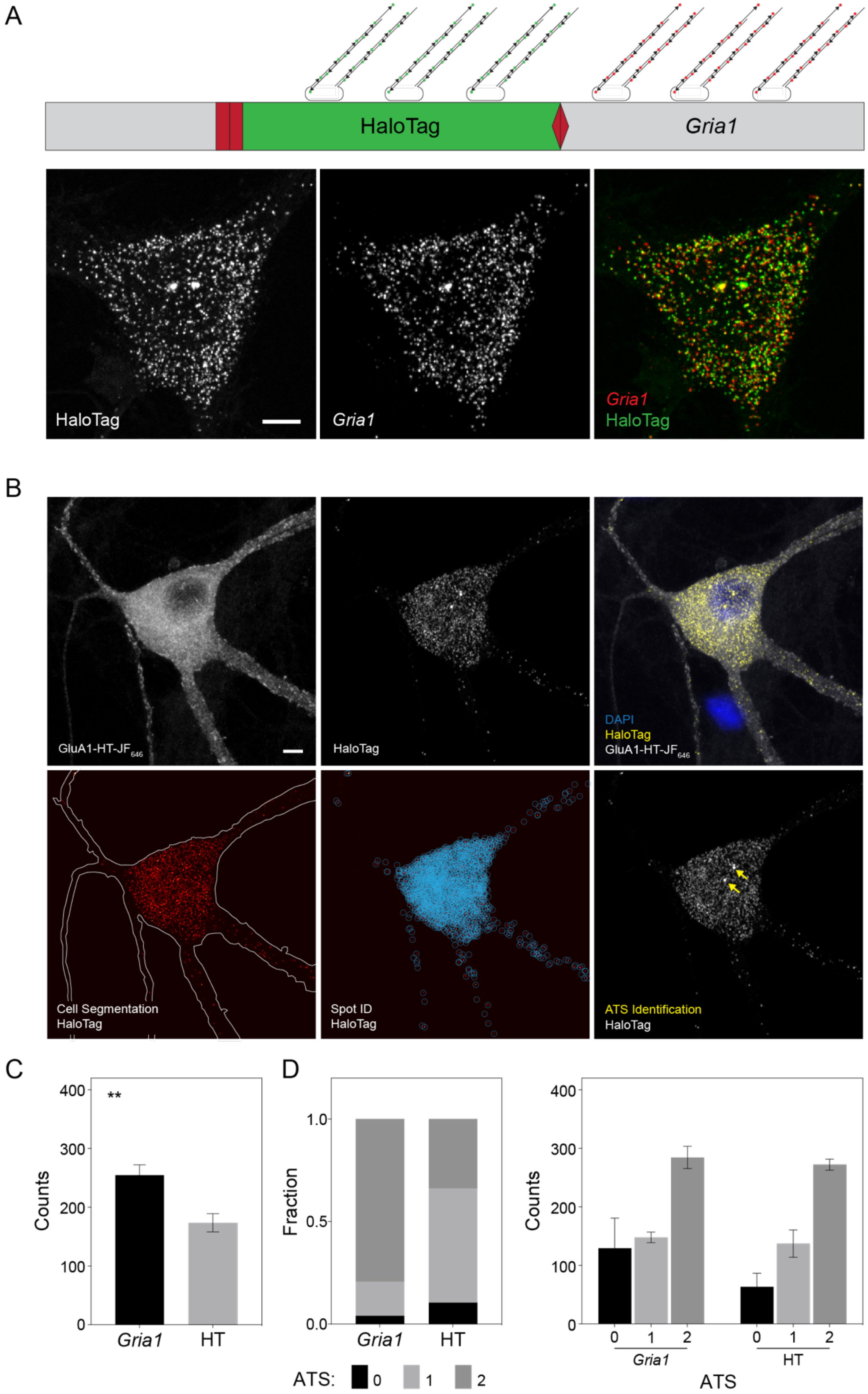

**Figure 1-figure supplement 2. Identification and quantification of HaloTag knock-in cells with HCR.**

(A) Schematic: two-color HCR RNA-FISH labeling targeting *Gria1* and HaloTag mRNA to validate HaloTag knock-in. Probes conjugated to Alexa Fluor 488 (green) were used to label HaloTag mRNA while probes conjugated to Alexa Fluor 546 (red) were used to label *Gria1* mRNA. Images: representative confocal images of a neuron labeled with HaloTag and *Gria1* probes. Scale bar, 5 μm.

(B) Top row: representative confocal images of a cell expressing GluA1-HT labeled with $JF_{646}$-HTL and also labeled with probes targeting HaloTag mRNA. Scale bar, 5 μm. Bottom row: to quantify HaloTag mRNA, neurons were segmented based on GluA1-HT-$JF_{646}$ labeling, then mRNA spots and ATS were identified using FISH-Quant.

(C) Average number of *Gria1* and HaloTag mRNA spots per cell. **p=0.0044 by t-test with Welch's correction. n=3 experiments.

(D) Left: fraction of cells that contain no ATS, one ATS, or two ATS when labeled with *Gria1* or HaloTag probes. The majority of cells labeled with HaloTag probes have only one ATS, likely because only one *Gria1* allele is tagged, accounting for lower mRNA spot count for HaloTag. Right: average number of mRNA spots per cell when cells are separated based on the number of ATS. There is no significant difference in the average number of mRNA spots per cell. Significance determined by t-test with Welch's correction.

Figure 1-figure supplement 3

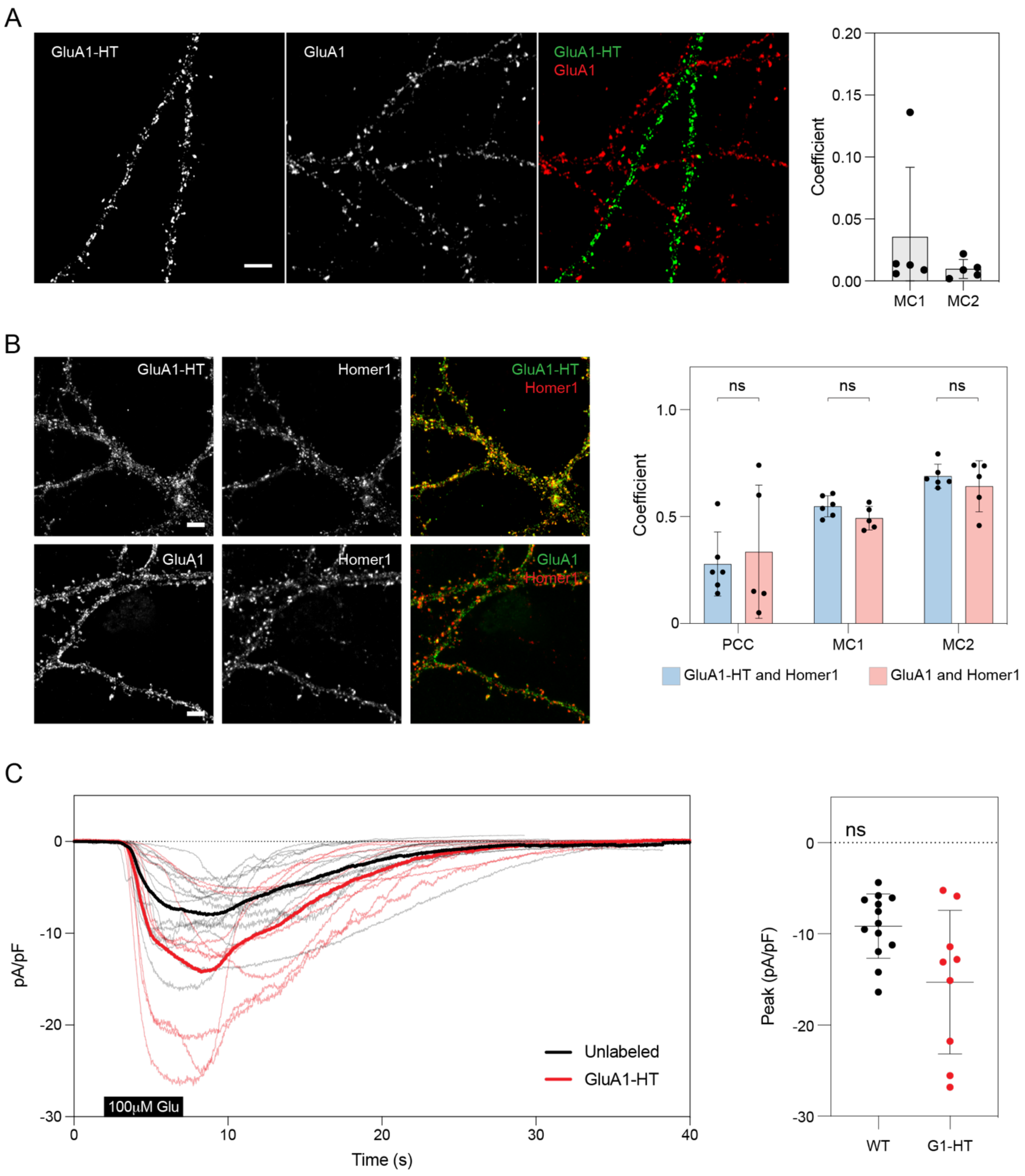

**Figure 1-figure supplement 3. Validation of HaloTag knock-in and GluA1-HT trafficking.**

(A) Images: representative confocal images of cells labeled with anti-HaloTag (GluA1-HT) and anti-GluA1 (GluA1) antibodies. For cells that are labeled with anti-HaloTag antibody, there is little or no labeling with anti-GluA1 antibody because HaloTag insertion disrupts the extracellular anti-GluA1 antibody epitope. Scale bar, 5 μm. Graph: Manders' coefficients (MC) of cells labeled with anti-HaloTag and anti-GluA1 antibodies. Bars represent mean and standard deviation, each dot represents a cell.

(B) Images: representative confocal images of dendrites labeled with anti-HaloTag antibody, anti-GluA1 antibody, and anti-Homer1 antibody from the same dish. Scale bars, 5 μm. Neurons that express GluA1-HT label weakly for GluA1. Graph: PCCs and MCs between GluA1-HT and Homer1 labeling vs GluA1 and Homer1 labeling are not significantly different. Bars represent mean and standard deviation, each dot represents a cell. Significance was determined by Mann-Whitney test.

(C) Currents elicited by GluA1-HT and unlabeled neurons in response to locally perfused glutamate. Each thin line represents the current elicited in an individual cell, while the thick lines represent the average currents. Peak current densities presented in **Figure 1G** (shown here on right) are derived from these trajectories. Current densities for GluA1-HT neurons and unlabeled neurons were recorded following a 5 sec 100 μM glutamate stimulation at a -70 mV holding potential.

Figure 2-figure supplement 1

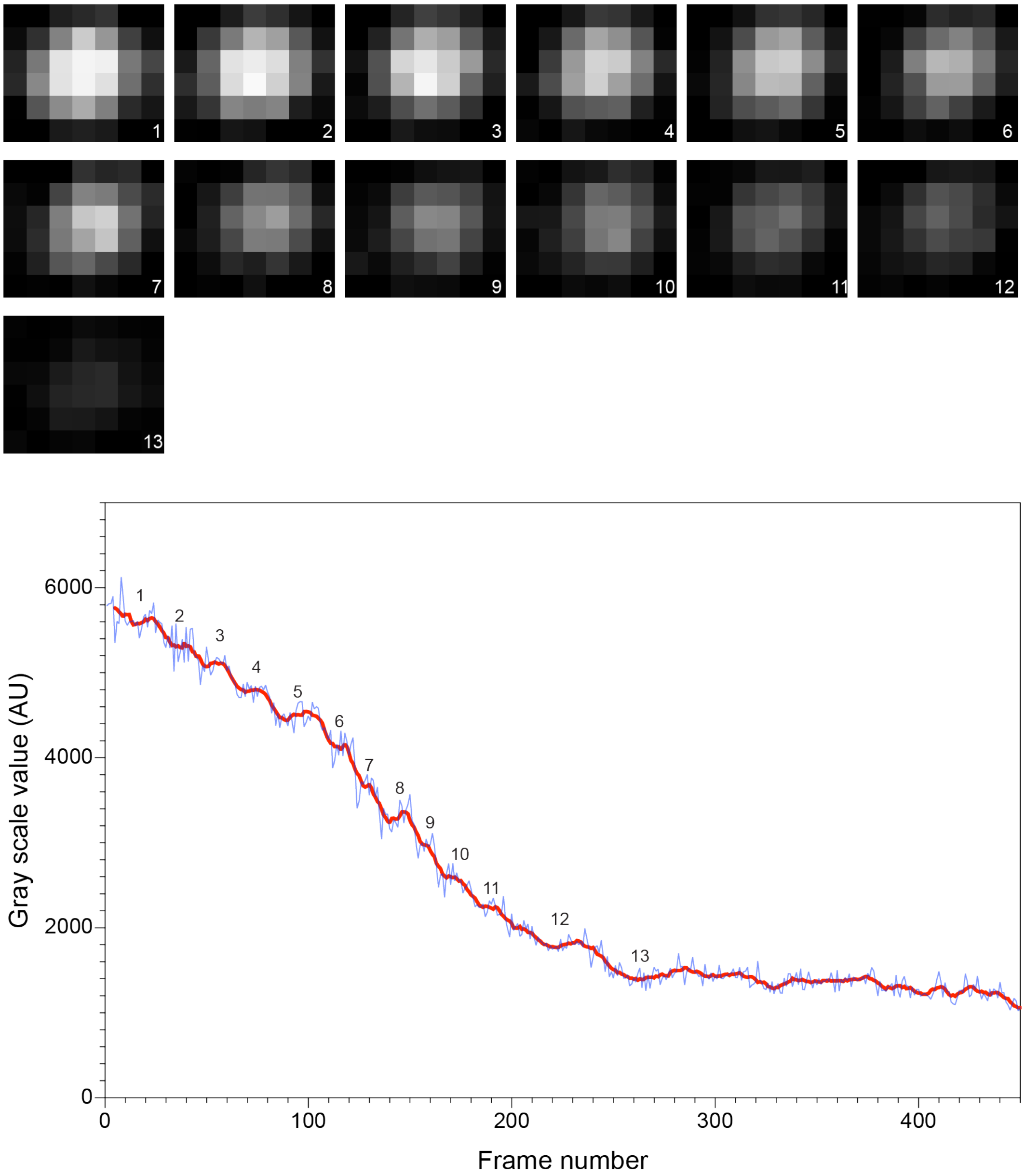

**Figure 2-figure supplement 1. Fluorescence signal from GluA1-HT-$JF_{549}$ vesicles bleaches in a stepwise fashion.**

Images: representative epifluorescence images of a GluA1-HT-$JF_{549}$ vesicle as it is bleached due to repeated light exposure. Graph: mean fluorescence intensity of vesicle plotted against frame number. Red line represents a running average of 9 frames. Number indicates step that corresponds to representative image. Vesicles contain multiple GluA1-HT particles and therefore can bleach over many steps. An individual GluA1 homomeric receptor will contain up to 4 labeled subunits, which will bleach in 4 steps. The most abundant AMPAR receptor type is the GluA1-GluA2 heterotetramer which will bleach in 1-2 steps.

Figure 2-figure supplement 2

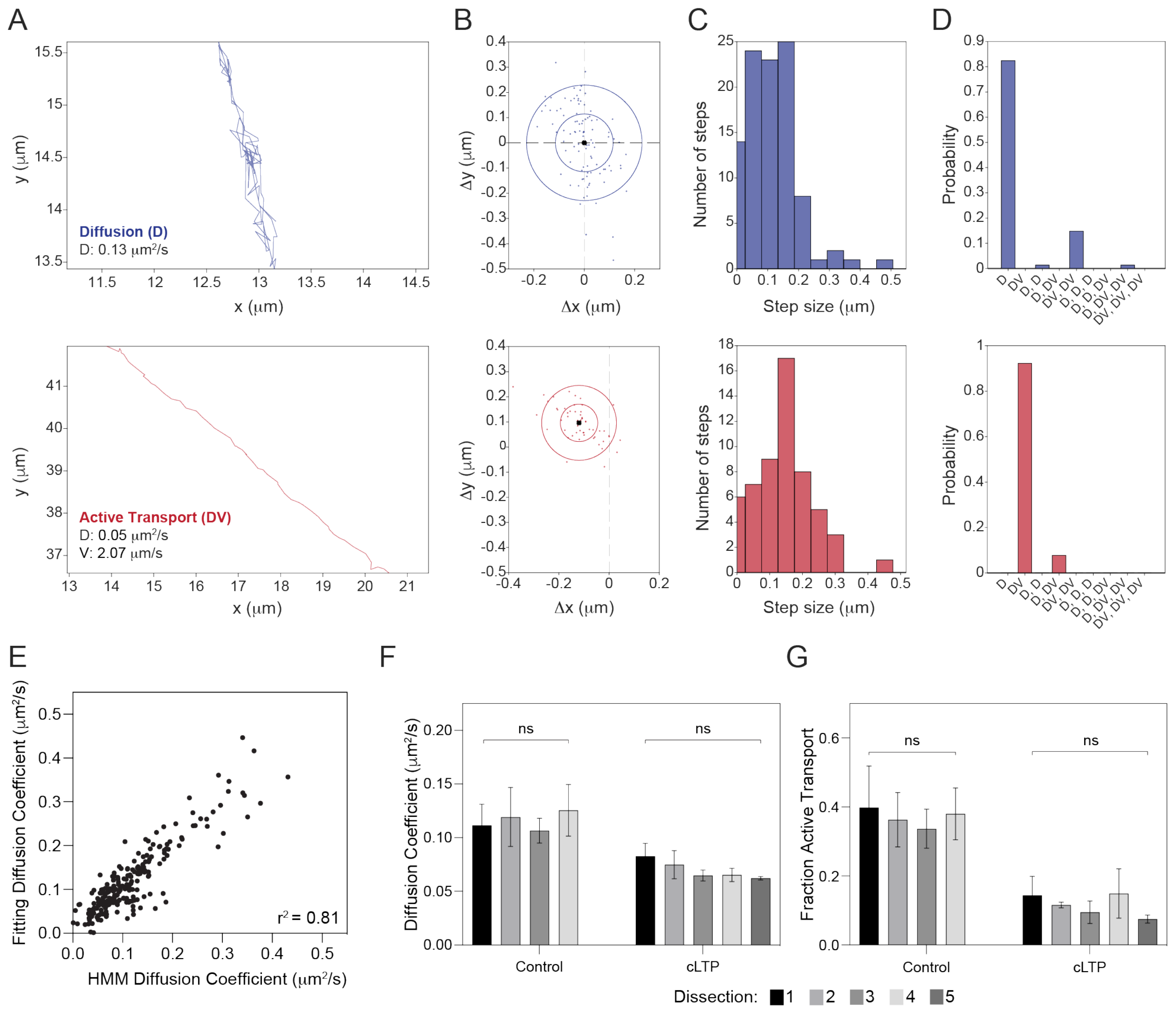

**Figure 2-figure supplement 2. HMM-Bayes analysis can be used to infer the motion states of GluA1-HT vesicles and determine motion parameters.**

(A) Representative trajectories for a GluA1-HT vesicle undergoing diffusion (D, top) and active transport (DV, bottom).

(B) Scatter plot of observed displacements (i.e., the vector between two successive points) along a trajectory for a vesicle exhibiting diffusion (top) or active transport (bottom).

(C) Distributions of step sizes for the displacements presented in B.

(D) Model probabilities for motion states inferred by HMM-Bayes based on the observed displacements and step size distributions in B and C. For a diffusive process, the displacements fit a normal distribution, from which HMM-Bayes can infer important motion parameters, such as the diffusion coefficient and velocity (shown in A). Importantly, HMM-Bayes accounts for active transport by allowing for distributions of displacements with nonzero means (for example, B, bottom).

(E) Scatterplot of diffusion coefficients determined by fitting the linear portion of mean square displacement curves (y-axis) versus diffusion coefficients inferred by HMM-Bayes (x-axis). Diffusion coefficients determined by each method are highly correlated, demonstrating that HMM-Bayes accurately infers diffusion coefficients for diffusing trajectories. n=232 diffusion coefficients.

(F) Diffusion coefficients for 2-3 time lapses from 4-5 different dissections (12-14 time lapses total) under control conditions and cLTP stimulation. Significance determined by Kruskal-Wallis test.

(G) Fractions of GluA1-HT vesicles with active transport for 2-3 time lapses from 4-5 different dissections (12-14 time lapses total) under control conditions and cLTP stimulation. Significance determined by Kruskal-Wallis test. These tests (F-G) demonstrate that the variability observed between different conditions is greater than the variability between dissections.

Figure 2-figure supplement 3

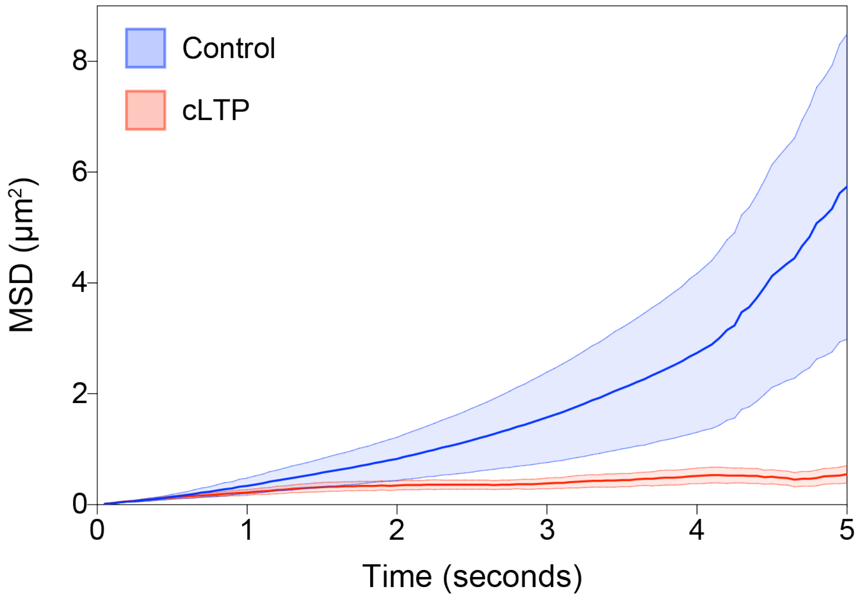

**Figure 2-figure supplement 3. GluA1-HT vesicles exhibit subdiffusive motion during cLTP induction.**

Mean square displacement (MSD) of GluA1-HT vesicles under no treatment control condition (Control, blue) and during cLTP induction (cLTP, red). Thick lines represent mean while area represents standard error. n=41-48 trajectories.

#### Figure 2-figure supplement 4

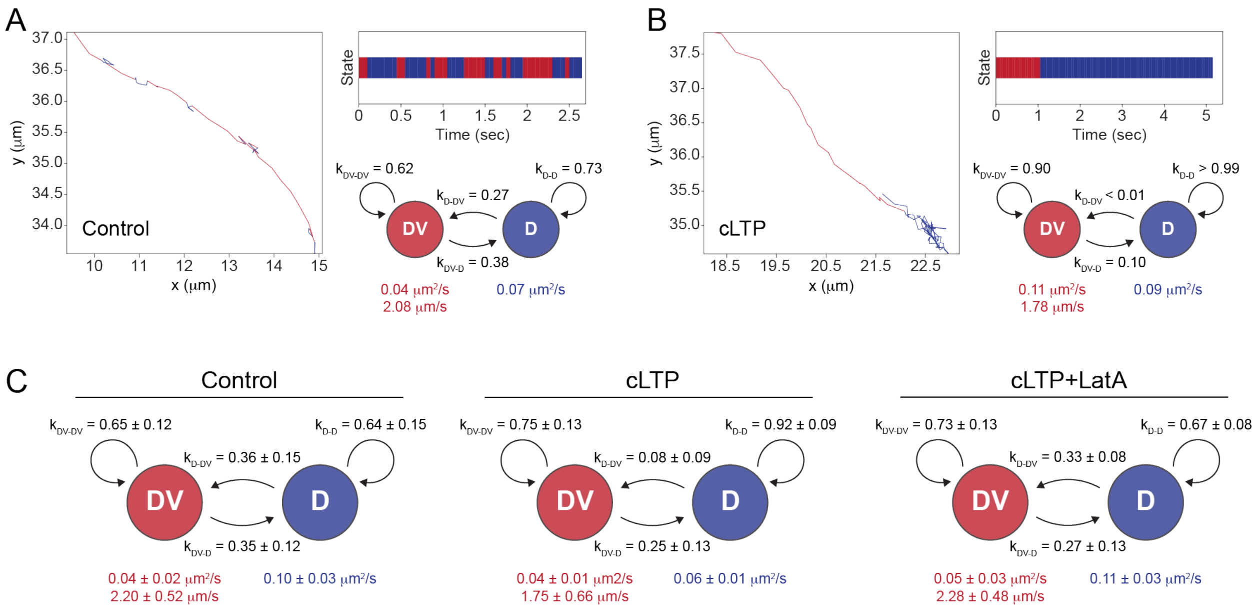

**Figure 2-figure supplement 4. cLTP induction increases the probability GluA1-HT vesicles switch from active transport to diffusion.**

(A) Left: representative trajectory of a GluA1-HT vesicle under control conditions exhibiting stochastic switching between active transport (DV) and diffusive motion (D). Right top: temporal state sequence inferred by HMM-Bayes of the trajectory depicted on the left. Right bottom: transition probabilities and motion parameters for each motion state for the trajectory depicted on the left. $K_{DV\text{-}DV}$, probability that a GluA1-HT vesicle remains in active transport; $K_{DV\text{-}D}$, probability that a vesicle switch from active transport to diffusion; $K_{D\text{-}DV}$, probability that a vesicle switch from diffusion to active transport; $K_{D\text{-}D}$, probability that a vesicle remains in diffusion. Values in red are the velocity and diffusion coefficient of the vesicle during active transport while the value in blue is the diffusion coefficient of the vesicle during diffusion.

(B) Left: representative trajectory of a GluA1-HT vesicle under cLTP induction exhibiting stochastic switching between active transport and diffusive motion. Right top: temporal state sequence inferred by HMM-Bayes of the trajectory depicted on the left. Right bottom: transition probabilities and motion parameters for each motion state for the trajectory depicted on the left.

(C) Average state-switching probabilities for multi-state GluA1-HT vesicles under control conditions (Control), cLTP induction (cLTP), and cLTP induction in the presence of LatA (cLTP+LatA). Control, $k_{DV\text{-}D}$ vs $k_{D\text{-}DV}$, $p=0.8065$; cLTP, $k_{DV\text{-}D}$ vs $k_{D\text{-}DV}$, ****$p<0.0001$; cLTP+LatA, $k_{DV\text{-}D}$ vs $k_{D\text{-}DV}$, *$p=0.0348$. Significance determined by Mann-Whitney test. n=26-30 trajectories for each condition.

Figure 3-figure supplement 1

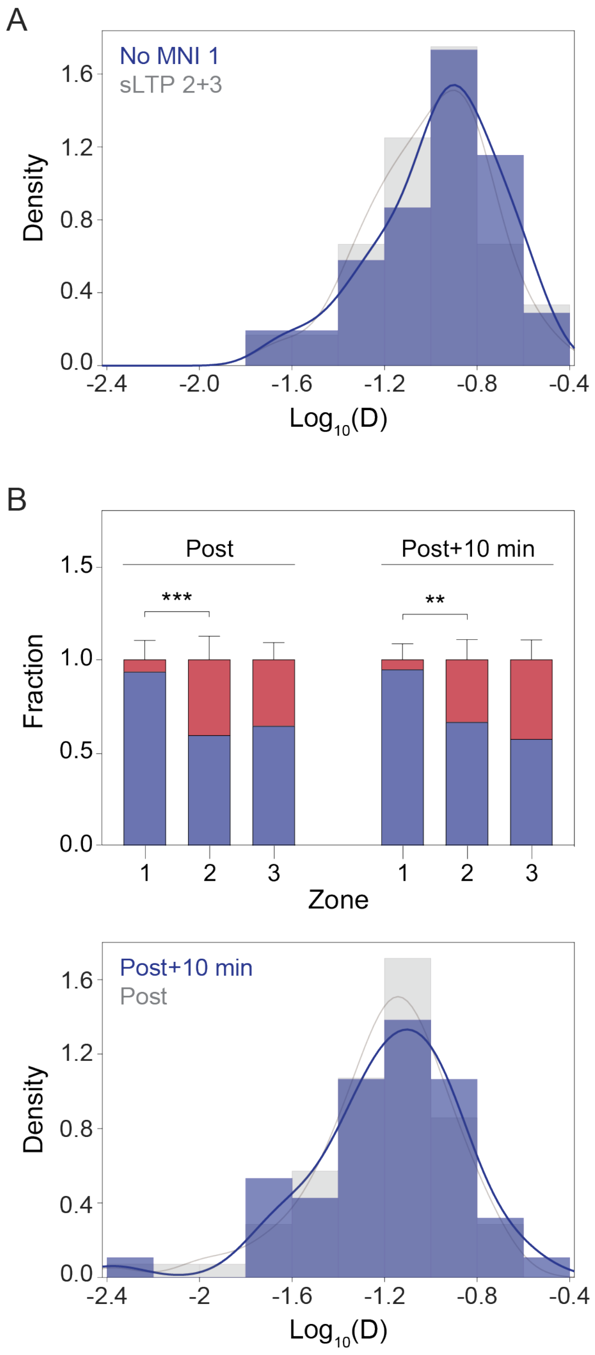

**Figure 3-figure supplement 1. Motion of GluA1-HT vesicles 10 minutes after the cessation of sLTP stimulation is similar to vesicles immediately after sLTP stimulation.**

(A) Distribution of diffusion coefficients in Zone 1 after sLTP stimulation in the absence of MNI (No MNI 1) vs Zone 2+3 in the presence of MNI (sLTP 2+3). n=52-60 trajectories for each condition.

(B) Bar graph: mean fractions of GluA1-HT vesicles exhibiting active transport or diffusion immediately after sLTP (Post) or 10 minutes after sLTP (Post+10 min) in each zone. Error bars represent standard deviation. Post+10 min: Zone 1 vs Zone 2, **p=0.0022 by Mann-Whitney test. n=6-9 experiments for each condition. Histogram: distribution of diffusion coefficients in Zone 1 immediately after sLTP stimulation (Post) vs 10 minutes after sLTP (Post+10 min). n=47-70 trajectories for each condition.

### Figure 3-figure supplement 2

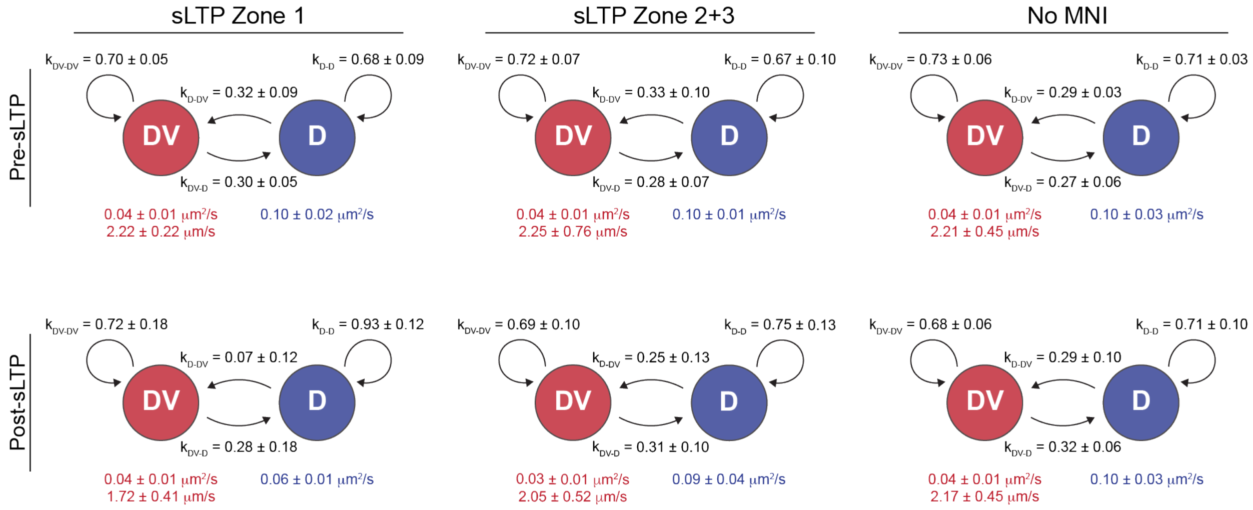

**Figure 3-figure supplement 2. sLTP increases the probability GluA1-HT vesicles switch from active transport to diffusion proximal to the site of stimulation.**

Average state-switching probabilities for multi-state GluA1-HT vesicles before and after sLTP stimulation in Zone 1 (Zone 1), Zone 2+3 (Zone 2+3), and in Zone 1 in the absence of MNI (No MNI). $K_{DV\text{-}DV}$, probability that a GluA1-HT vesicle remains in active transport (DV); $K_{DV\text{-}D}$, probability that a vesicle switch from active transport to diffusion (D); $K_{D\text{-}DV}$, probability that a vesicle switch from diffusion to active transport; $K_{D\text{-}D}$, probability that a vesicle remains in diffusion. Values in red are the velocity and diffusion coefficient of the vesicle during active transport while the value in blue is the diffusion coefficient of the vesicle during diffusion. sLTP Zone 1: pre-sLTP, $k_{DV\text{-}D}$ vs $k_{D\text{-}DV}$, $p=0.7104$; post-sLTP, $k_{DV\text{-}D}$ vs $k_{D\text{-}DV}$, $^*p=0.0195$. sLTP Zone 2+3: pre-sLTP, $k_{DV\text{-}D}$ vs $k_{D\text{-}DV}$, $p=0.6200$; post-sLTP, $k_{DV\text{-}D}$ vs $k_{D\text{-}DV}$, $p=0.5887$. No MNI: pre-sLTP, $k_{DV\text{-}D}$ vs $k_{D\text{-}DV}$, $p=0.3829$; post-sLTP, $k_{DV\text{-}D}$ vs $k_{D\text{-}DV}$, $p=0.8182$. Significance determined by Mann-Whitney test. n=6-7 trajectories for each condition.

Figure 4-figure supplement 1

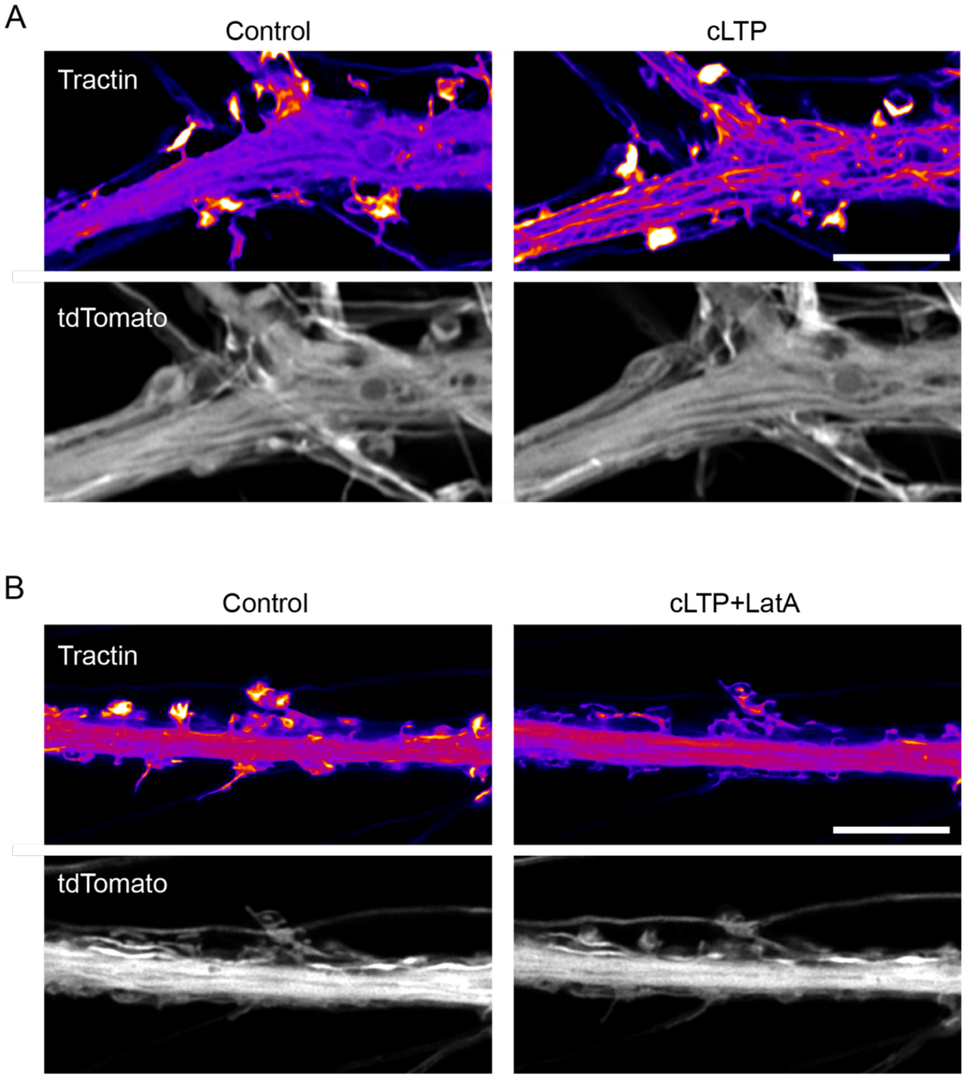

**Figure 4-figure supplement 1. cLTP stimulation leads to the redistribution of tractin without changing the morphology of dendrites.**

(A) Representative Airyscan images of tractin and tdTomato in a dendrite under control conditions and during cLTP. Scale bar, 5 μm.

(B) Representative Airyscan images of tractin and tdTomato in a dendrite under control conditions and during cLTP in the presence of LatA. Scale bar, 5 μm.

Figure 4-figure supplement 2

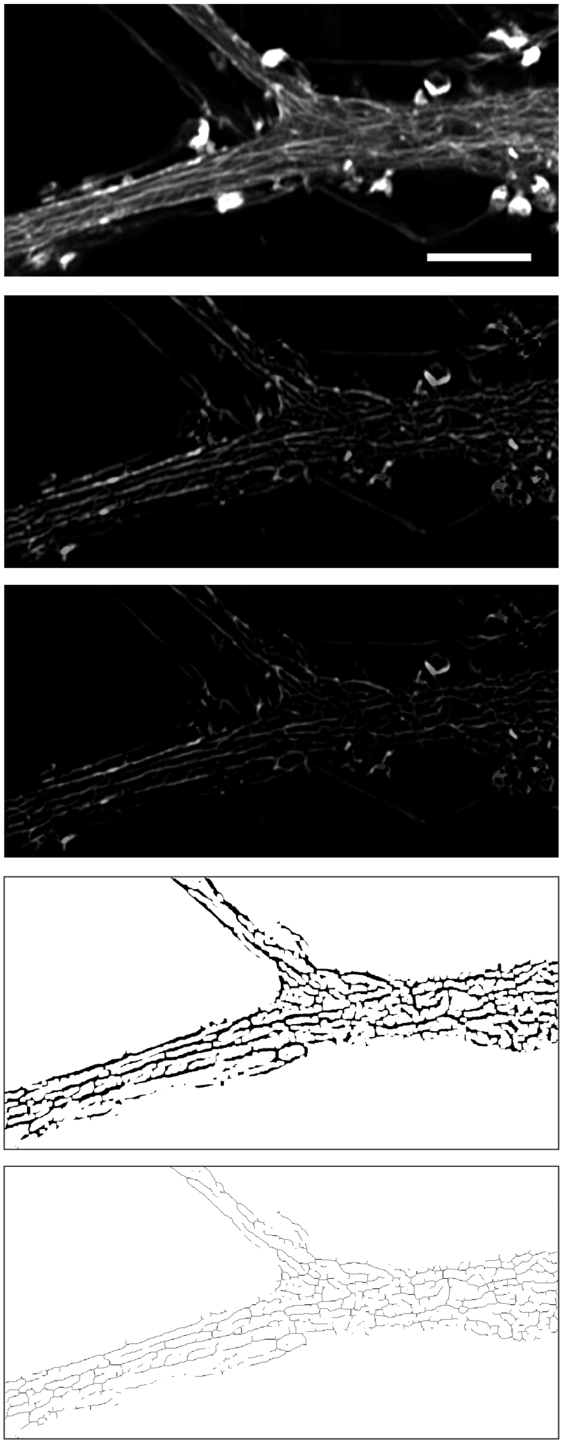

Airyscan Image

Background Subtraction

Minimum Filter

Binary Mask

Network Skeleton

**Figure 4-figure supplement 2. F-actin polymerization quantified by measuring skeleton length of tractin network.**

To create a skeleton of the tractin network, the background from an Airyscan image is first subtracted with a rolling ball radius of 10 pixels. Next, a minimum filter of 1 pixel is applied to the background subtracted image. An Otsu threshold is then applied to the image to create a binary mask. Otsu’s method is an established automatic intensity thresholding method that enables us to minimize subjective analysis between samples and conditions. Spines are removed from the binary mask to analyze tractin in the dendritic shaft. Finally, the binary mask is skeletonized and analyzed using the Analyze Skeleton ImageJ plugin.

Figure 4-figure supplement 3

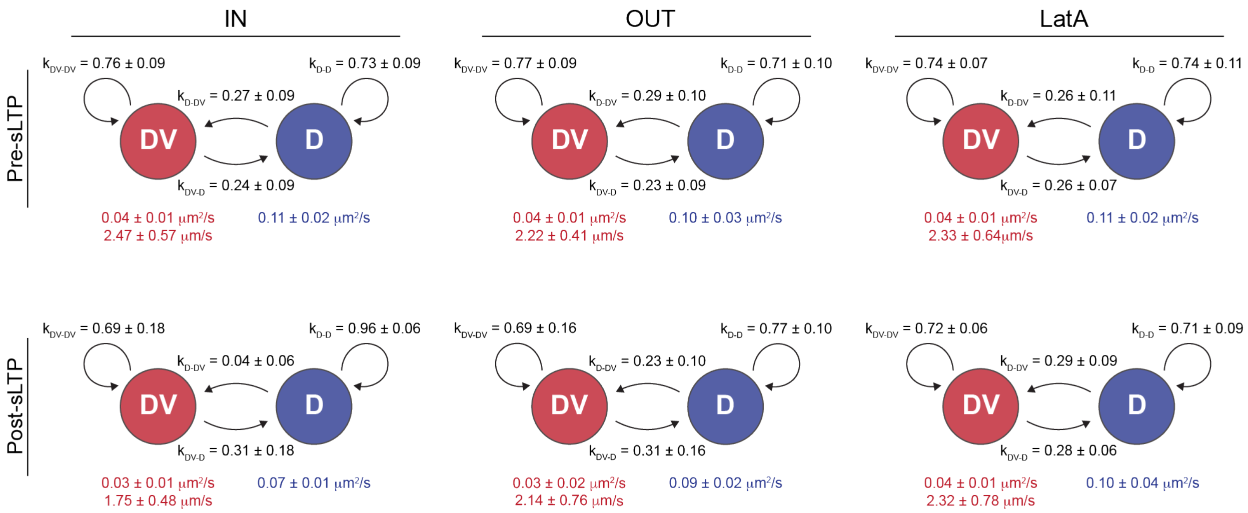

**Figure 4-figure supplement 3. sLTP-induced actin polymerization increases the probability GluA1-HT vesicles switch from active transport to diffusion.**

Average state-switching probabilities for multi-state GluA1-HT vesicles before and after sLTP stimulation inside regions with actin polymerization (IN), outside regions with actin polymerization (OUT), and in the presence of LatA (LatA). $K_{DV-DV}$, probability that a GluA1-HT vesicle remains in active transport (DV); $K_{DV-D}$, probability that a vesicle switch from active transport to diffusion (D); $K_{D-DV}$, probability that a vesicle switch from diffusion to active transport; $K_{D-D}$, probability that a vesicle remains in diffusion. Values in red are the velocity and diffusion coefficient of the vesicle during active transport while the value in blue is the diffusion coefficient of the vesicle during diffusion. IN: pre-sLTP, $k_{DV-D}$ vs $k_{D-DV}$, p=0.4848; post-sLTP, $k_{DV-D}$ vs $k_{D-DV}$, *p=0.0317. OUT: pre-sLTP, $k_{DV-D}$ vs $k_{D-DV}$, p=0.3823; post-sLTP, $k_{DV-D}$ vs $k_{D-DV}$, p=0.4848. LatA: pre-sLTP, $k_{DV-D}$ vs $k_{D-DV}$, p=0.9015; post-sLTP, $k_{DV-D}$ vs $k_{D-DV}$, p>0.9999. Significance determined by Mann-Whitney test. n=5-8 trajectories for each condition.

Figure 5-figure supplement 1

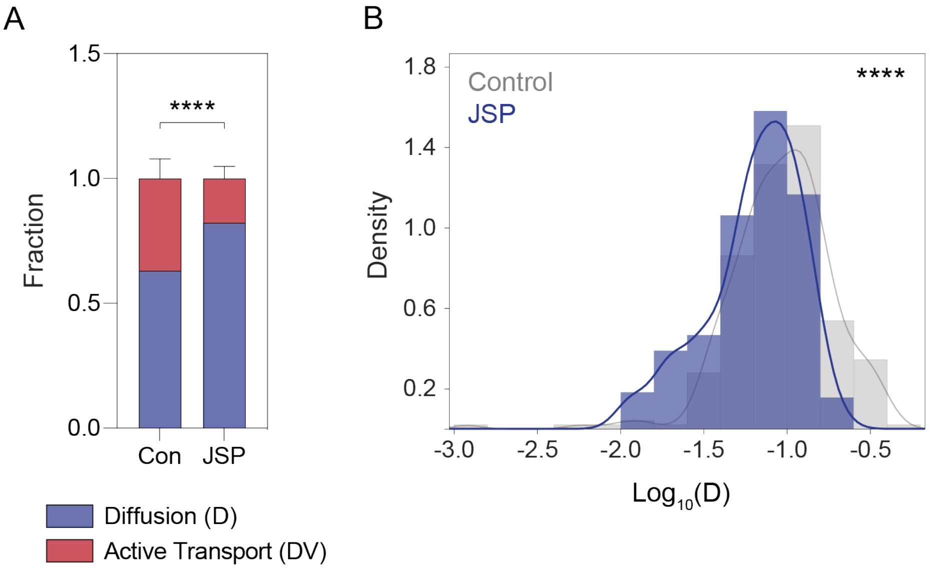

**Figure 5-figure supplement 1. Jasplakinolide treatment reduces the fraction of GluA1-HT vesicles exhibiting active transport and decreases the diffusion coefficient of vesicles exhibiting diffusion.**

(A) Bar graph of the fractions of GluA1-HT vesicles exhibiting active transport (DV) or diffusion (D) in dendritic shafts under control conditions (Con) versus during treatment with Jasplakinolide (JSP). Con vs JSP, ****$p<0.0001$. Significance was determined by Mann-Whitney test. n=9 experiments for each condition. Bars represent mean and standard deviation.

(B) Distributions of diffusion cofficients of GluA1-HT vesicles in dendritic shafts under control conditions (Con) versus during treatment with Jasplakinolide (JSP). Con vs JSP, ****$p<0.0001$. Significance was determined by Kolmogorov-Smirnov test. n=194-232 trajectories for each condition. Lines represent the probability density function of each histogram estimated by kernel density estimation (KDE).

Figure 6-figure supplement 1

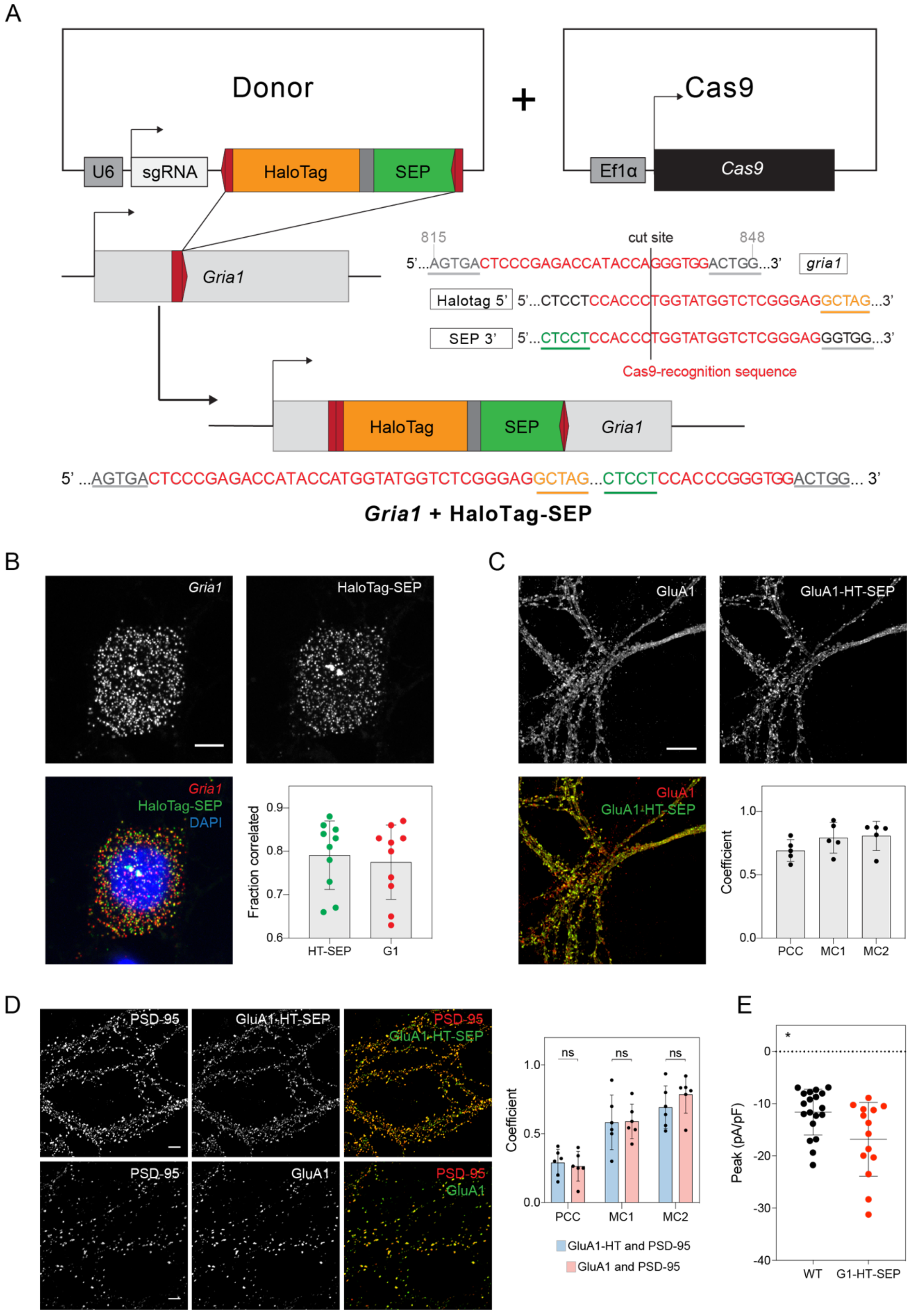

**Figure 6-figure supplement 1. Endogenous GluA1 labeled with HaloTag-SEP reporter via homology-independent targeted integration is trafficked to postsynaptic densities and responds to stimulation.**

(A) Schematic of *Gria1* gene targeting with HaloTag-SEP by homology-independent targeted integration (HITI).

(B) Images: representative confocal images of a two-color HCR RNA-FISH labeling experiment with probes targeting HaloTag-SEP mRNA (HaloTag-SEP) or *Gria1* mRNA (*Gria1*). Scale bar, 5 µm. Bar graph: fraction of HaloTag-SEP mRNA spots located within 0.5 µm of *Gria1* mRNA spots and vice versa.

(C) Images: representative confocal images of a dendrite labeled with anti-HaloTag antibody (GluA1-HT-SEP) and anti-GluA1 antibody (GluA1). Scale bar, 5 µm. Bar graph: Pearson correlation coefficients (PCC) and Manders' coefficients (MC1 and MC2) between HaloTag and GluA1 labeling. Bars represent mean and standard deviation.

(D) Images: representative confocal images of dendrites labeled with anti-HaloTag antibody (GluA1-HT-SEP), anti-GluA1 antibody (GluA1), and anti-PSD-95 antibody (PSD-95) from the same dish. Scale bars, 5 µm. Bar graph: PCCs and MCs between GluA1-HT-SEP and PSD-95 labeling vs GluA1 and PSD-95 labeling. Bars represent mean and standard deviation. Significance was determined by Mann-Whitney test.

(E) Currents elicited by GluA1-HT-SEP and unlabeled neurons in response to locally perfused glutamate. Crosshairs represent mean and standard deviation of peak current densities (pA/pF) for each condition. Significance was determined by Mann-Whitney test. Current densities for GluA1-HT neurons and unlabeled neurons were recorded following a 5 sec 100 µM glutamate stimulation at a -70 mV holding potential. Each dot represents a neuron or recording. *p=0.0192 by Mann-Whitney test. GluA1-HT-SEP exhibits a modest but significant increase in current, possibly reflecting some disruptive properties of the HT-SEP tandem reporter. Nevertheless, GluA1-HT-SEP is trafficked to synapses in a similar manner to GluA1, and its function is not inhibited by the insertion of HT-SEP.

Figure 7-figure supplement 1

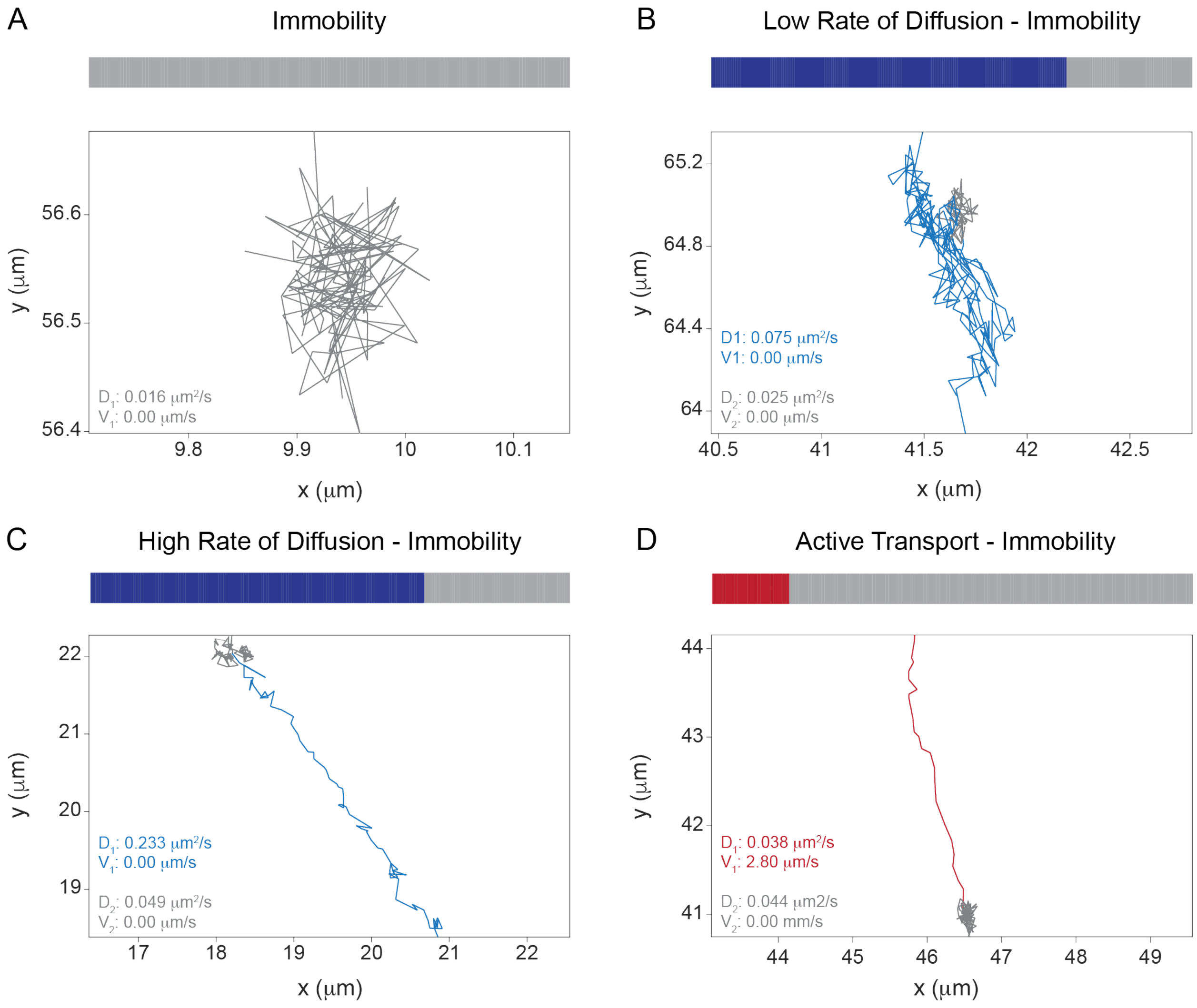

**Figure 7-figure supplement 1. Pre-exocytosis GluA1-HT-SEP vesicles trajectories have two motion states.**

(A-D) Temporal state sequences (top) and trajectories (bottom) for pre-exocytosis GluA1-HT-SEP vesicles exhibiting (A) immobility, (B) low rate of diffusion followed by immobility, (C) high rate of diffusion followed by immobility, and (D) active transport followed by immobility.

Figure 7-figure supplement 2

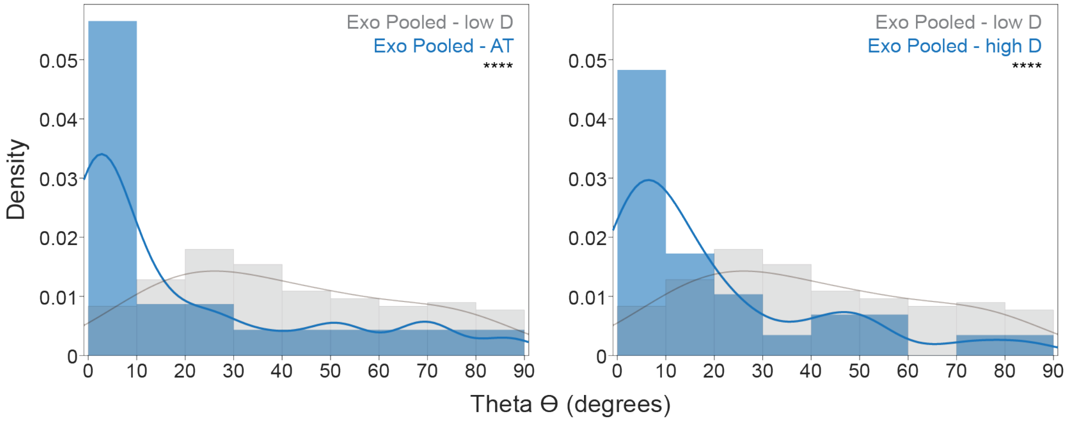

**Figure 7-figure supplement 2. Pre-exocytosis GluA1-HT-SEP vesicle trajectories that exhibit active transport or high rates of diffusion have a longitudinal directional bias.**

Distributions of theta (θ) for pre-exocytosis GluA1-HT-SEP vesicles exhibiting low rates of diffusion or immobility (Exo Pooled – low D) versus pre-exocytosis GluA1-HT-SEP vesicles exhibiting active transport (Exo Pooled – AT, left) or high rates of diffusion (Exo Pooled – high D, right). ****$p < 0.0001$ by Kolmogorov-Smirnov test. n=24-156 trajectories for each condition. Trajectories were pooled from Con-Exo, cLTP-Exo, No MNI-Exo IN, and sLTP-Exo IN. Lines represent the probability density function of each histogram estimated by kernel density estimation (KDE).

#### Movie legends

**Video 1 (related to Figure 2)**

Time-lapse sequence of GluA1-HT-$JF_{549}$ (green) after block-and-chase labeling with $JF_{646}$-HTL and $JF_{549}$-HTL overlaid onto GFP-Homer1c (magenta) in the dendrite of a cultured rat hippocampal neuron. Scale bar, 10 μm.

**Video 2 (related to Figure 2)**

Time-lapse sequence of GluA1-HT-$JF_{549}$ (green) after block-and-chase labeling with $JF_{646}$-HTL and $JF_{549}$-HTL overlaid onto GFP-Homer1c (magenta) under control conditions. Scale bar, 10 μm.

**Video 3 (related to Figure 2)**

Time-lapse sequence of GluA1-HT-$JF_{549}$ (green) after block-and-chase labeling with $JF_{646}$-HTL and $JF_{549}$-HTL overlaid onto GFP-Homer1c (magenta) during cLTP induction. Scale bar, 10 μm.

**Video 4 (related to Figure 3)**

Time-lapse sequences of GluA1-HT-$JF_{549}$ after block-and-chase labeling with $JF_{646}$-HTL and $JF_{549}$-HTL before sLTP (Pre-sLTP, left) and after sLTP (Post-sLTP, right) in the dendrite of a cultured rat hippocampal neuron. Scale bar, 10 μm.

**Video 5 (related to Figure 4)**

Time-lapse sequences of GluA1-HT-$JF_{549}$ after block-and-chase labeling with $JF_{646}$-HTL and $JF_{549}$-HTL before sLTP (Pre-sLTP, left) and after sLTP (Post-sLTP, right) in the dendrite of a cultured rat hippocampal neuron. Scale bar, 10 μm. Performed in neuron also expressing tractin to characterize the motion of GluA1-HT vesicles in areas where actin polymerization occurs.

**Video 6 (related to Figure 5)**

Time-lapse sequence of GEM-HT-ST labeled with $JF_{549}$-HTL in the dendrite of a cultured rat hippocampal neuron under control conditions. Scale bar, 10 μm.

**Video 7 (related to Figure 6)**

Time-lapse sequence of GluA1-HT-SEP exocytosis in the dendrite of a cultured rat hippocampal neuron under control conditions. Scale bar, 5 µm.

**Video 8 (related to Figure 7)**

Time-lapse sequence of GluA1-HT-SEP labeled with $JF_{549}$-HTL (green) – after block-and-chase labeling with $JF_{646}$-HTL and $JF_{549}$-HTL – overlaid with GluA1-HT-SEP exocytosis events from the same imaging experiment (magenta) after sLTP in the dendrite of a cultured rat hippocampal neuron. Scale bar, 5 µm.
